# Discovery of archaeal Fusexins homologous to eukaryotic HAP2/GCS1 gamete fusion proteins

**DOI:** 10.1101/2021.10.13.464100

**Authors:** David Moi, Shunsuke Nishio, Xiaohui Li, Clari Valansi, Mauricio Langleib, Nicolas G. Brukman, Kateryna Flyak, Christophe Dessimoz, Daniele de Sanctis, Kathryn Tunyasuvunakool, John Jumper, Martin Graña, Héctor Romero, Pablo S. Aguilar, Luca Jovine, Benjamin Podbilewicz

**Author notes:** These authors contributed equally to this work (David Moi, Shunsuke Nishio, Xiaohui Li).

## Abstract

Sexual reproduction consists of genome reduction by meiosis and subsequent gamete fusion. Presence of meiotic genes in prokaryotes suggests that DNA repair mechanisms evolved toward meiotic recombination; however, fusogenic proteins resembling those found in eukaryotes were not identified in prokaryotes. Here, we identify archaeal proteins that are homologs of fusexins, a superfamily of fusogens that mediate eukaryotic gamete and somatic cell fusion, as well as virus entry. The crystal structure of a trimeric archaeal Fusexin1 reveals novel features such as a six-helix bundle and an additional globular domain. Ectopically expressed Fusexin1 can fuse mammalian cells, and this process involves the additional domain and a conserved fusion loop. Archaeal fusexin genes exist within integrated mobile elements, potentially linking ancient archaeal gene exchanges and eukaryotic sex.

**One-Sentence Summary:** Cell membrane fusion proteins of viruses and eukaryotes are also present in archaea.

## Main text

How early eukaryotes developed the capacity for gamete fusion is a central question entangled with the origin of the eukaryotic cell itself. The widespread presence of a conserved set of meiosis, gamete and nuclear fusion proteins among extant eukaryotes indicates that meiotic sex emerged once, predating the last eukaryotic common ancestor (LECA) (*1, 2*). The conserved gamete fusion protein HAP2/GCS1 belongs to a superfamily of fusion proteins (fusogens) called fusexins (*3*–*5*). This superfamily encompasses class II viral fusogens (viral fusexins) that fuse the envelope of some animal viruses with the membranes of host cells during infection (*6, 7*); EFF-1 and AFF-1 (somatic fusexins) that promote cell fusion during syncytial organ development (*8*–*11*); and HAP2/GCS1 (sexual fusexins; hereafter referred to as HAP2) that mediate gamete fusion (*12*–*14*). Although it is assumed that sexual fusexins were already present in the LECA (*1, 15*), their shared ancestry with viral fusexins posed a “the virus or the egg” evolutionary dilemma (*16*). In one scenario, fusexins are proper eukaryal innovations that were captured by some viruses and used for host invasion. Alternatively, a viral fusexin gene was transferred to an early eukaryotic cell and then repurposed for gamete fusion.

Here, we identify a fourth family of fusexins in genomes of Archaea and prokaryotic fractions of metagenomes from very diverse environments. We provide structural and functional evidence indicating that these proteins are cellular fusogens. Genomic analyses indicate that archaeal fusexins are carried by integrated mobile genetic elements. Evolutionary analyses of the whole fusexin superfamily reveal new working models for the emergence of meiotic sex during eukaryogenesis.

## Results

### Fusexin genes in Archaea

To search for fusexins we used the crystallographic structures of *C. reinhardtii, A. thaliana*, and *T. cruzi* HAP2 (Cr/At/TcHAP2) (*4, 17, 18*) to build dedicated Hidden Markov Models (HMMs). These were used to scan the Uniclust30 database with HHblits (see Methods in Supplementary Materials). We detected 24 high-confidence candidates in prokaryotes: eight belong to isolated and cultivated archaea, and the remaining sixteen to metagenome-assembled genomes (MAGs, table S1). We then built HMMs of the candidate ectodomains and compared them to HMMs of sexual, somatic and viral fusexins. Fig. 1A shows that the prokaryotic candidates have detectable sequence similarities with HAP2, with E-values below 0.001 and HHblits-derived probabilities higher than 0.95 (fig. S1). We named these proteins Fusexin1 (Fsx1). *fsx1* genes found in cultivated and isolated prokaryotes are restricted to the Halobacteria class (Euryarchaeota superphylum) whereas MAGs containing Fsx1s include all major Archaea superphyla (table S1). Next, we used this Fsx1 sequence set to search the Metaclust database, which comprises 1.59 billion clustered proteins from over 2200 metagenomic/metatranscriptomic datasets. Performing a scan pipeline using PHMMER, PSI-BLAST, HMM-HMM comparisons and topology filtering we found 96 high-confidence *fsx1* genes. The identified *fsx1*s come from different environments (with preeminence of saline samples) and a wide temperature range (−35 to 80ºC, data S1).

**Fig. 1.**
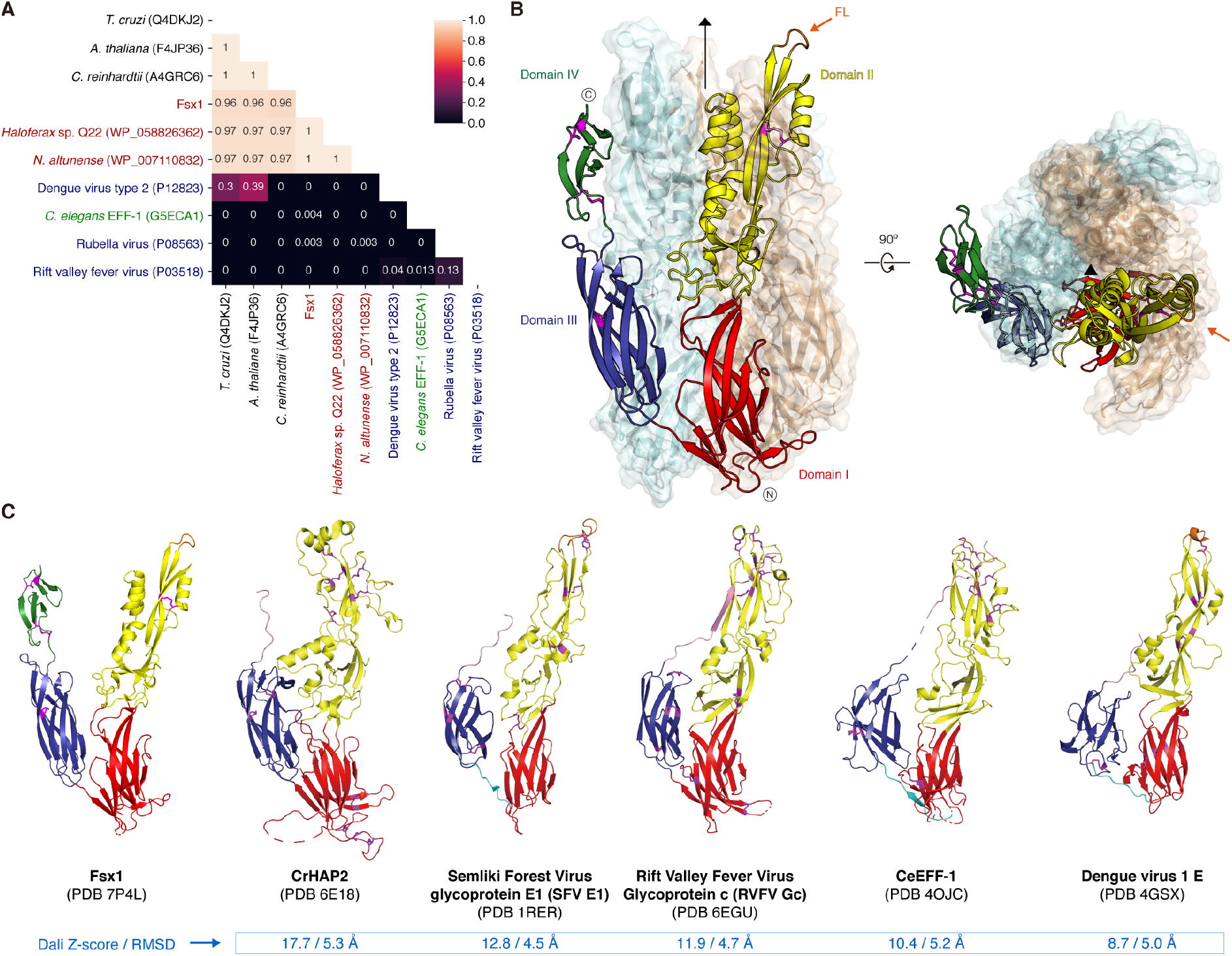
Fsx1 is a member of the fusexin protein superfamily. (**A**) HMM vs HMM homology probabilities of a subset of eukaryotic, viral and archaeal fusexin ectodomains. With exception of Fsx1, which derives from a metagenomic sequence, archaeal fusexins (red), viral fusexins (blue), EFF-1 (green) and HAP2s (black) are indicated by RefSeq/UniProt identifiers. (**B**) Crystal structure of the trimeric ectodomain of Fsx1. The three-fold non-crystallographic axis is indicated. Subunit A is shown as a cartoon coloured by domain, with disulfides and the fusion loop (FL) coloured magenta and orange, respectively; subunits B, C are in mixed cartoon/surface representation. (**C**) Side-by-side comparison of Fsx1_E_ and known class II fusogens. Elements are coloured as in panel b; the stem region and the linker between domains I and III are shown in pink and cyan, respectively.

### Fsx1 is a structural homolog of HAP2/GCS1

To experimentally investigate the presence of fusexin-like proteins in Archaea, a selection of the candidate genes was expressed in mammalian cells (fig. S2A, B). High-level expression was observed for a metagenomic Fsx1 sequence from a hypersaline environment, predicted to encode a ∼55 kDa ectodomain region (Fsx1_E_) followed by three transmembrane domains (TMs) (data S1). Crystals of Fsx1_E_, obtained in the presence of 2.5 M NaCl, 0.2 M CaCl_2_, yielded data to 2.3 Å resolution that was phased by molecular replacement using a combination of fragments from models generated by AlphaFold2 (*19*) (figs. S3 and S4; table S2).

Fsx1_E_ is a monomer in solution but crystallized as a homotrimer of 119×78×75 Å (Fig. 1B and figs. S3 and S5). Each protomer consists of four domains, the first three of which match the approximate dimensions and relative arrangement of domains I-III of known fusexins in their post-fusion conformation (*20*); accordingly, fold and interface similarity searches identify HAP2 as the closest structural homolog of Fsx1_E_, followed by viral fusexins and *C. elegans* EFF-1 (Fig. 1B, C and fig. S6). Fsx1 domains I and III are relatively sequence-conserved among archaeal homologs (figs. S7A and S8A) and closely resemble the corresponding domains of HAP2 (RMSD 2.1 Å over 218 Cα), including the invariant disulfide bond between domain III strands βC and βF (*4*) (C_3_389-C_4_432; figs. S5C and S6). On the other hand, Fsx1 domain II shares the same topology as that of HAP2 but differs significantly in its secondary structure elements and their relative orientation, as well as disulfide bonds (fig. S6C). In particular, Fsx1 domain II is characterized by a four-helix hairpin, whose N-terminal half interacts with the same region of the other two subunits to generate a six-helix bundle around the molecule’s three-fold axis (Figs. 1B and 2, A to C, and figs. S5A and S7, B and C).

**Fig. 2.**
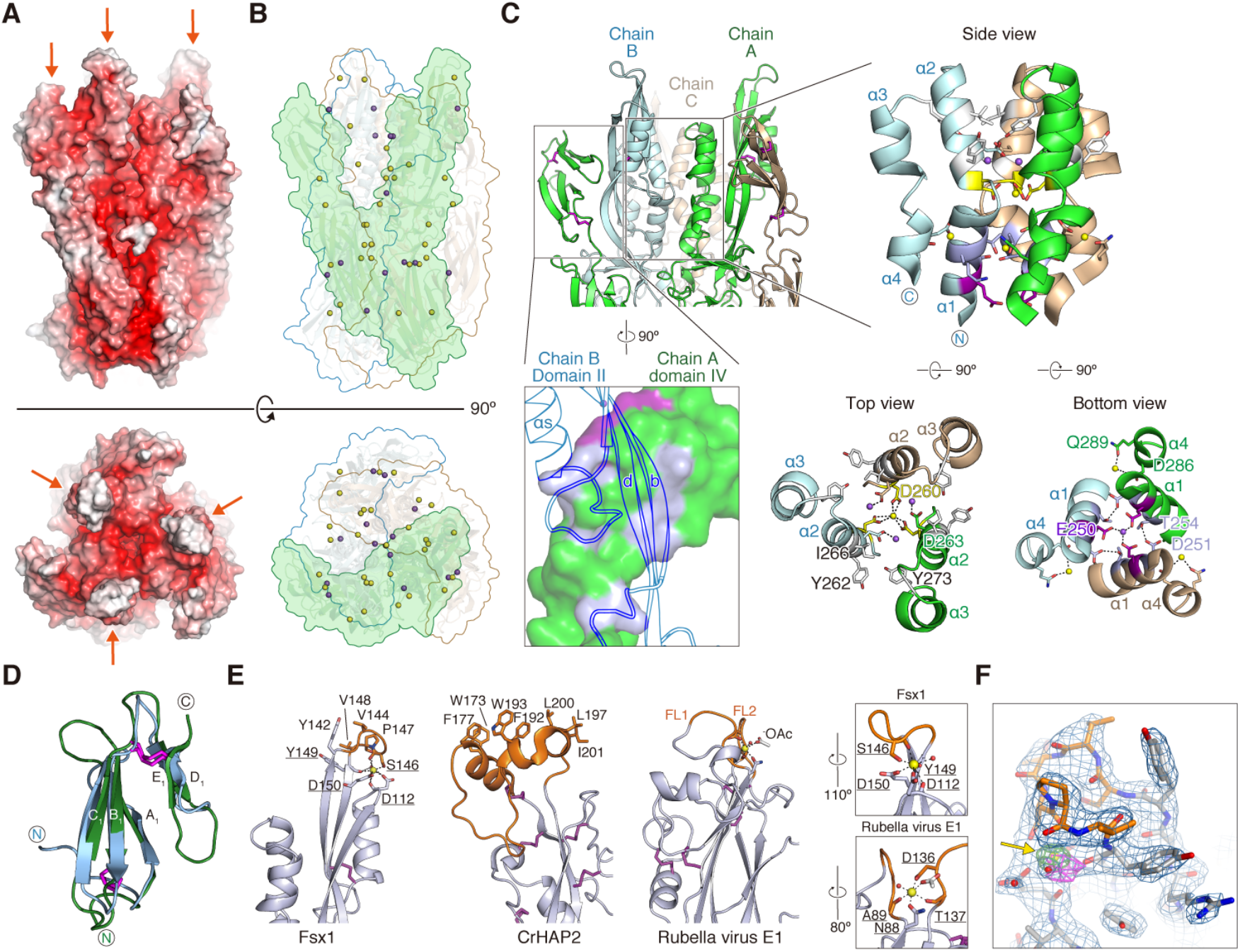
Distinct structural features of Fsx1. (A) Fsx1_E_ surface coloured by electrostatic potential from red (−10 kT/e) to blue (+10 kT/e) through white (0 kT/e). Orange arrows indicate the FLs. (B) Location of the Ca^2+^ and Na^+^ ions (depicted as yellow and purple spheres, respectively; see also figs. S5 and S7) stabilizing the Fsx1_E_ trimer. The molecular surface of a protomer is shaded green and the outline of the other two subunits is coloured cyan and wheat. (C) Details of interactions at the level of the six-helical bundle made by domain II of the Fsx1 subunits (right subpanels) and the domain IV/domain II interface (bottom left subpanel). Selected side chains are colored by the type of inter-chain contacts in which they are involved (gray: hydrophobic interaction; blue bell: hydrogen bonding; yellow: Ca^2+^ coordination; fuchsia: Na^+^ coordination), with dashed lines indicating hydrogen bonds. Note that the helical bundle of Fsx1 is not a leucine-zipper coiled-coil structure, such as those found in class I/III viral fusion proteins or in the SNARE four-helix bundles, and see also fig. S7B. (D) Superposition of Fsx1 domain IV (green) and Der p 23 (PDB 4CZE, blue) (Dali Z-score 3.6, RMSD 2.2 Å). (E) Comparison of the Fsx1 region that includes the FL and the corresponding parts of CrHAP2 and Rubella virus E1 protein (PDB 4B3V). Residues coordinating the Ca^2+^ ion that stabilizes the Fsx1 FL are underlined, and compared to the E1 protein Ca^2+^-binding region in the boxed panels on the far right. (F) The Fsx1 FL adopts a highly ordered conformation stabilized by a Ca^2+^ ion. Presence and identity of the latter, indicated by a yellow arrow, are supported by two other maps shown in addition to the *2mFo-DFc* map (blue mesh, contoured at 1.0 σ): a difference map calculated upon omitting all metal ions from the model (thick green(+)/red(−) mesh, 6.0 σ) and a phased anomalous difference map calculated from a 2.9 Å-resolution dataset collected at 7.1 KeV (thick magenta mesh, 3.2 σ).

Notably, unlike previously characterized viral and eukaryotic fusexins, Fsx1 also contains a fourth globular domain conserved among archaeal homologs (Fig. 1, B and C, and figs. S5D, S6 and S8). Its antiparallel β-sandwich, which includes the two C-terminal disulfides of Fsx1, resembles the carbohydrate-binding fold of dust mite allergen Der p 23 and related chitin-binding proteins (*21*) (Fig. 2D); accordingly, it is also structurally similar to a high-confidence AlphaFold2 model of the C-terminal domain of acidic mammalian chitinase (*22*). In addition to being coaxially stacked with domain III as a result of a loop/loop interaction stabilized by the C_5_457-C_6_477 disulfide, domain IV contributes to the quaternary structure of the protein by interacting with domain II of the adjacent subunit to which domain III also binds (Figs. 1B and 2C).

The Fsx1_E_ monomer has a net charge of -67, and another feature stabilizing its homotrimeric assembly is a set of Ca^2+^ and Na^+^ ions that interacts with negatively charged residues at the interface between subunits (Fig. 2A, B). Additional metal ions bind to sites located within individual subunits; in particular, a Ca^2+^ ion shapes the conformation of the domain II c-d loop (S143-V148) so that its uncharged surface protrudes from the rest of the molecule (Fig. 2 and fig. S7D). Strikingly, the position of this element matches that of the fusion loops (FLs) of other fusexins, including the Ca^2+^-binding fusion surface of rubella virus E1 protein (*23, 24*) (Fig. 2E). Moreover, as previously observed in the case of CrHAP2 (*18*), the loops of each trimer interact with those of another trimer within the Fsx1 crystal lattice.

In summary, despite significant differences in the fold of domain II, the unprecedented presence of a domain IV, and extreme electrostatic properties, the overall structural similarity between Fsx1 and viral or eukaryotic fusexins suggests that this prokaryotic molecule also functions to fuse membranes.

### Fsx1 can fuse eukaryotic cells

To test the fusogenic activities of the candidate archaeal fusexins we studied their fusion activity upon transfection in eukaryotic cells (*3, 10, 11*). Cells with either red or green nuclei are mixed with each other and fusion is measured by the formation of hybrid cells with both red and green nuclei revealing merger of their cytoplasms. For this, we co-cultured two batches of Baby Hamster Kidney (BHK) cells independently transfected with Fsx1 and co-expressing either nuclear H2B-RFP or H2B-GFP (*3*). We then performed immunofluorescence against a V5 tag fused to the cytoplasmic tail of Fsx1 (Fig. 3A and B, and fig. S9). We observed a five-fold increase in the mixing of the nuclear H2B-GFP and H2B-RFP compared to vector control, showing that Fsx1 is a *bona fide* fusogen, as efficient as the fusexin AtHAP2 (Fig. 3C). To determine whether Fsx1 expression is required in both fusing cells or, alternatively, it suffices in one of the fusing partners, we mixed BHK-Fsx1 coexpressing cytoplasmic GFP with BHK cells expressing only nuclear RFP. We found increased multinucleation of GFP+ cells (revealing cell-cell fusion) but very low mixing with RFP+ cells not expressing Fsx1. In contrast, the vesicular stomatitis virus G-glycoprotein (VSVG) fusogen induced efficient unilateral fusion (*11*) (Fig. 3D-F). Thus, Fsx1 acts in a bilateral way, similarly to EFF-1 and AFF-1 fusexins (*11, 25, 26*). We then performed live imaging using spinning disk confocal microscopy and observed cell-cell fusion of BHK-Fsx1 cells (Fig. 3, G and H).

**Fig. 3.**
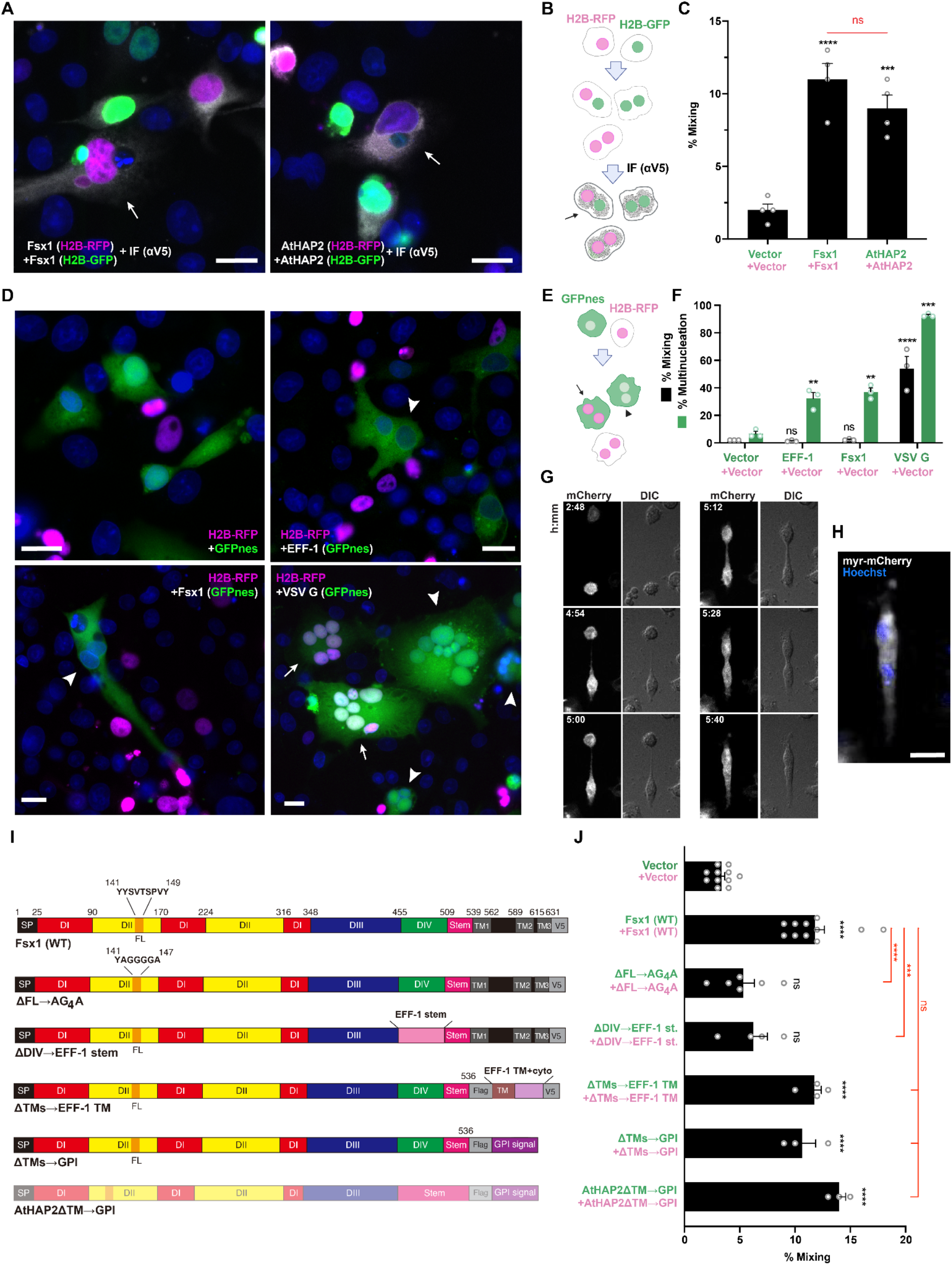
Fsx1 mediates bilateral cell-cell fusion. (A to C) Cell-cell fusion was measured by content-mixing, indicated by the appearance of multinucleated cells containing green nuclei (H2B-GFP) and magenta nuclei (H2B-RFP). Immunofluorescence against the V5 tag was also performed (gray). (A) Images of mixed cells. DAPI, blue. See also fig. S9A. (B) Scheme of experimental design. (C) Quantification of content-mixing. The mixing indices presented as means ± SEM of four independent experiments. Comparisons by one-way ANOVA followed by Bonferroni’s test. ns = non-significant, *** p < 0.001, **** p< 0.0001. (D to F) Unilateral fusion was evaluated by mixing control cells expressing nuclear H2B-RFP (magenta) with cells expressing GFP with a nuclear export signal (GFPnes, green cytoplasm) only or together with Fsx1, EFF-1 or VSV G. (D) Images of cells transfected with empty::GFPnes vector, Fsx1::GFPnes, EFF-1::GFPnes or VSV G::GFPnes. Fsx1 and EFF-1 show multinucleated GFPnes positive cells (arrowheads). VSV G multinucleated cells are found with GFP only (arrowheads) or with both markers (arrows). Scale Bars, 20 µm. See also fig. S9E. (E) Scheme of experimental design. (F) Quantification of content-mixing experiments in which only the GFP population of cells express Fsx1, EFF-1, VSV G or none of them (vector). Bar chart showing means ± SEM of three independent experiments. Comparisons by one-way ANOVA followed by Dunett’s test against the vector. ns = non-significant, ** p < 0.01, *** p < 0.001, **** p < 0.0001. (G) Spinning disk microscopy time-lapse images indicating the merging of two cells expressing myr-mCherry and Fsx1. Time in hours:minutes. The red channel (mCherry, white) and the DIC are shown. Refer to Movie S1. (H) For the last point a Z-projection showing the myr-mCherry fluorescence (white) and the nuclei Hoechst (blue; Movie S2). Scale bar, 20 µm. (I to J) Structure function analysis of Fsx1. (I) Schematic diagram of wild type Fsx1 four mutants and AtHAP2ΔTM→GPI. SP, signal peptide; FL, fusion loop (Fig. 2E). (J) Quantification of content-mixing in populations of cells expressing vectors (n=7), Fsx1 (wt) (n=7), its mutants (ΔFL→AG4A (n=6), ΔDIV→EFF-1 stem (n=4), ΔTMs→EFF-1 TM (n=4), ΔTMs→GPI (n=3), or AtHAP2ΔTM→GPI (n=3) (see fig. S10). Bar chart showing means ± SEM. Comparisons by one-way ANOVA followed by Bonferroni’s test against the vector (black) and against Fsx1 (red). ns=non-significant, *** p < 0.001, **** p< 0.0001.

### Structure-function analysis of Fsx1

To compare archaeal Fsx1 activity with fusexins from eukaryotes and viruses, we introduced mutations into three different structural domains of Fsx1 and tested surface expression and fusogenic activity in mammalian cells.

First, to test whether the putative FL of Fsx1 (143-SVTSPV-148) is involved in fusion, we replaced it with a linker of 4G between Y142A and Y149A (Figs. 2E and 3I, and fig. S8; ΔFL→AG_4_A). This replacement does not affect surface expression yet reduces content mixing to levels similar to those of the negative control (Fig. 3J and fig. S10).

Second, we asked whether domain IV, which is only present in archaeal fusexins, has a function in the fusion process. For this, we replaced the entire domain with the stem region of EFF-1 (Fig. 3I; ΔDIV→EFF-1 stem). While this mutant Fsx1 reaches the cell surface, suggesting that it folds normally, it shows a significantly reduced activity compared to wildtype Fsx1 (Fig. 3J and fig. S10).

Third, to test whether the three TMs of Fsx1 are required for fusion, we replaced them with the TM and cytoplasmic domains of EFF-1 (Fig. 3I; ΔTMs→EFF-1 TM) or a glycosylphosphatidylinositol (GPI) anchor signal (Fig. 3I; ΔTMs→GPI). We found that both Fsx1 mutants remained active (Fig. 3J), indicating that the Fsx1 TMs are not essential for fusion. Finally, we also replaced the TM and cytoplasmic domains of AtHAP2 with a signal for GPI and found that the protein also maintained its fusogenic activity (Fig. 3, C, I, and J). Thus, contrary to some viral fusogens in which the GPI-anchored glycoproteins fail to drive complete fusion (*27*–*29*), lipid-anchored Fsx1 or eukaryotic HAP2 promote syncytia formation when expressed on the surface of BHK cells.

### Fsx1s are ancient fusogens associated with integrated mobile elements

The sparse pattern of Fsx1 presence in Archaea led us to perform genomic comparisons of related species with and without the *fsx1* gene. These revealed >50 kbp DNA insertions in the genomes of species with *fsx1* genes (fig. S12). To investigate them, we performed k-mer spectrum analysis on *fsx1*-containing Pure Culture Genomes (PCGs) and found divergent regions containing the *fsx1* ORF (Fig. 4A and fig. S13). Gene content analyses of *fsx1*-containing regions (fig. S14) show that they share a portion of their genes (fig. S15) and display conserved synteny (Fig. 4B and fig. S16), suggesting common ancestry. These regions are enriched in ORFs homologous to proteins involved in DNA mobilization and integration (Fig. 4B and table S3). Thus, our results indicate that *fsx1* genes are contained in integrated mobile elements (IMEs) that can be mobilized by a conjugative-like, cell fusion-dependent mechanism.

**Fig. 4.**
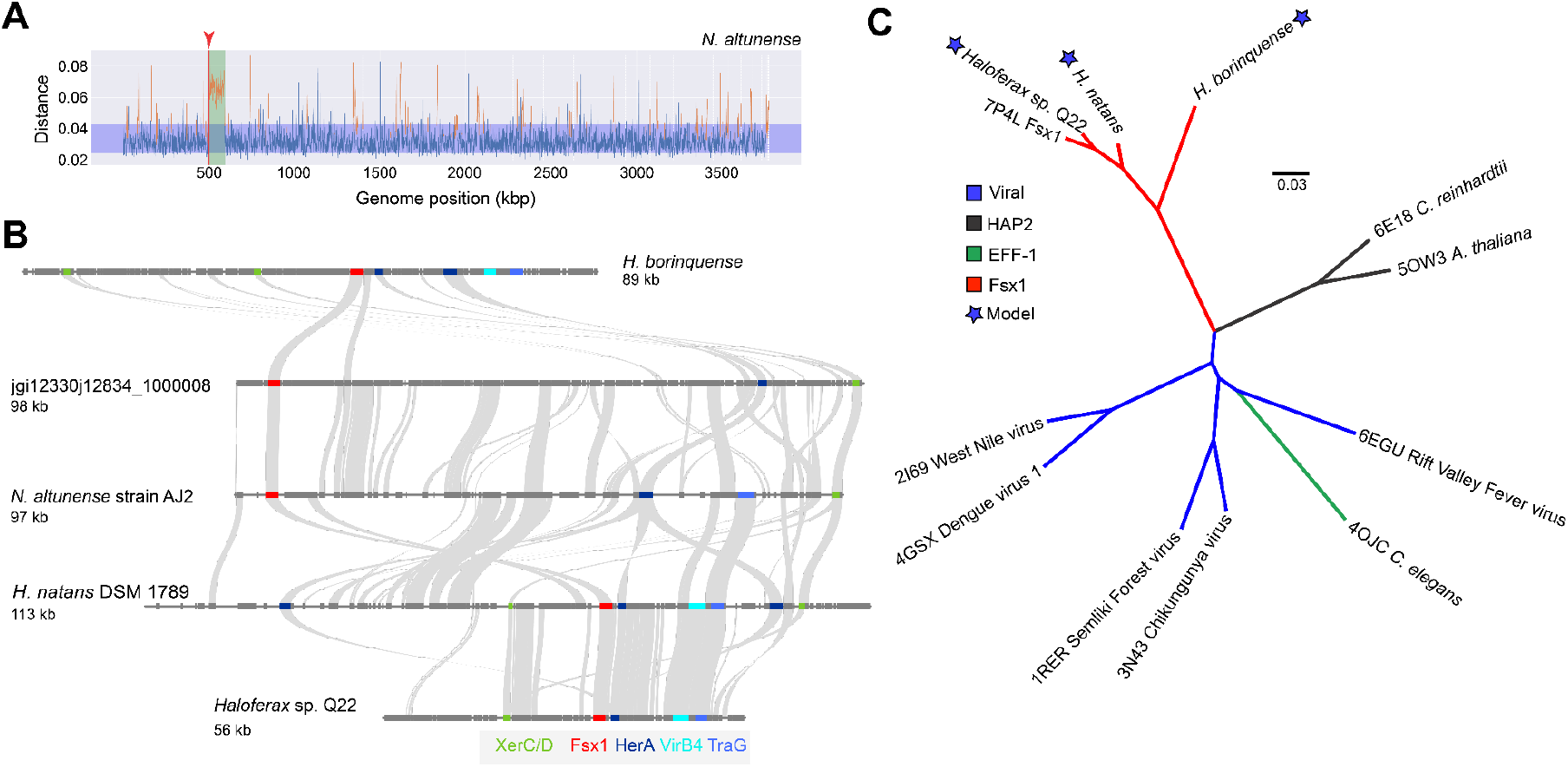
Genomic features of Fsx1s and evolutionary history of fusexins. (A) Fsx1s are embedded in integrated mobile elements (IMEs). *Natrinema altunense* AJ2 complete genome k-mer spectrum deviation from centroid. Blue region shows the standard deviation. Locus of *fsx1* is indicated by the red arrowhead; the mobile element containing *fsx1* is in green. (B) Four IMEs from PCG sequences and the metagenomic sequence from which the crystallized Fsx1 was obtained (second from top) are represented by their annotated genes (thick segments). Gray links are drawn between homologs. The *fsx1* gene is marked in red and selected ORFs homologous to IME signature genes are labeled and color-coded. XerC/XerD recombinases from family integrase (green); HerA helicase (dark blue); VirB4, Type IV secretory (T4SS) pathway (cyan); TraG/TraD/VirD4 family enzyme (light blue). See also fig. S16, and tables S1 and S3. (C) Computational models based on the Fsx1_E_ crystal structure and experimental structures of other fusexins were compared using flexible structural alignment to build a minimum evolutionary tree. Scale bar represents distance as 1-TM_score_ (see Methods). These distances indicate that all structures are homologous.

To describe Fsx1’s tempo and mode of evolution we first compared archaeal and sexual fusexins, which share enough sequence conservation to apply standard phylogenetic analyses, not possible for somatic and viral fusexins (Fig. 1A and fig. S1). We built maximum likelihood (ML) phylogenies for a set of Fsx1 sequences derived from isolated species and metagenomes, and a subset of HAP2s encompassing a wide breadth of eukaryotic lineages. Two major findings come from these phylogenies. First, the branching pattern of Fsx1 sequences from complete genomes is incompatible with their species tree (fig. S17), supporting a history of Horizontal Gene Transfer (HGT) events within Archaea, in line with *fsx1s* presence in IMEs. Second, eukaryal fusexins still cluster into a strongly supported clade suggesting they diverged before LECA.

To place *fsx1* in the broader fusexin superfamily context, we performed structural phylogenetic analysis comparing crystal structures from viral, somatic and eukaryotic gamete fusogens (Fig. 4C). This structure-based tree supports a viral origin of somatic fusexins (EFF-1) (*10*) and is also compatible with archaeal fusexins appearing before the radiation of eukaryotes.

## Discussion

The archaeal proteins herein identified place fusexins in yet another domain of life, with different membrane chemistry and along a wide niche landscape, from cold hypersaline lakes to hot springs and hydrothermal vents (fig. S17).

Our structural and functional analyses show that Fsx1 has both conserved and diverged properties when compared to eukaryotic and viral fusexins (Fig. 3 and fig. S6). Like its viral counterparts, Fsx1 has an uncharged loop required for fusion. However, unlike previously known fusexins, Fsx1 harbors an additional domain (IV) involved in fusogenic activity that may bind sugars (Figs. 2D and 3J). Considering that cell surface glycosylation was found to be important for fusion-based mating of halophilic archaea (*30*), this domain may actively promote fusion by interacting with carbohydrates attached to lipids or proteins such as S-layer glycoproteins (*31*). Unlike HAP2s, Fsx1 homologs have 1-4 TMs and a variable Cys number (5 to 30, see data S1). Like eukaryotic fusexins, Fsx1 mediates BHK cell fusion in a bilateral fashion (Fig. 3F). However, in contrast to viral fusogens (*27*–*29, 32, 33*), the fusion activity of Fsx1 is maintained following substitution of its three TMs with a single TM or a GPI anchor. Since GPI-anchored AtHAP2 is also fusogenic, other fusexins may also drive complete cell fusion without a specific involvement of TMs. Future studies will address the function of the six-helix bundle formed by Fsx1 domain II, which is unprecedented among fusexins and raises an unexpected structural connection with class I viral fusogens (*6, 7*).

The “virus or the egg” dilemma (*16*) posits that fusexins may have been viral innovations (class II fusogens), later acquired by eukaryotes, or vice versa. Archaeal fusexins expand this dilemma: the origins of gamete fusion may have had a prokaryotic player. Both structure- and sequence-based trees (Fig. 4C and fig. S17) do not solve but provide insights to address this predicament, in which we distinguish three main alternative origins: viral, eukaryotic and archaeal. Although exaptation of viral genes by eukaryotic organisms is well documented (*34*), known viral fusexins are restricted to very few divergent lineages infecting multicellular eukaryotes, suggesting that different viruses captured fusexin genes in different occasions. The Virus-first hypothesis requires more than one HGT event from an ancient viral lineage into different domains. Support for the existence of such fusexin-containing viral lineage infecting both archaeal and eukaryotic hosts is currently lacking (*35*).

The ubiquity of sexual fusexins in Eukarya strongly suggests evolutionary success, in line with the Eukarya-first hypothesis. However, examples of interdomain gene transfers from eukaryotes to archaeas are rare (*36*). The presence of Fsx1s in Halobacteria IMEs is consistent with HGT in the opposite direction. In addition to the Asgard- and alphaproteobacterial endosymbiont-related inherited genes, the pre-LECA may have received hundreds of archaeal genes from other lineages, including Euryarchaeota (*37*). Halophilic archaea are polyploid (*38*) and undergo HGT events that overcome species and genera barriers (*39, 40*). Fsx1s seem absent in some archaeal species known to undergo fusion-based gene exchange events (*41*). As genetic exchange in extant prokaryotes is dominated by conjugation, transduction and transformation, we hypothesize that Fsx1s are molecular fossils still playing a role in archaeal ‘sex-like’ events.

Overall, the pre-LECA divergence between archaeal and eukaryal fusexins, their presence in mobile elements and our current understanding of the role played by HGT during eukaryogenesis (*42*–*44*) prompt us to consider the Archaea-first as a plausible hypothesis.

Discovery of the Asgard superphylum (*45*) and the recent cultivation of one of its members (*46*) support eukaryogenesis models where populations of bacteria and archaea lived in syntrophy, transferring metabolites and genes (*47*). Acquisition of a *fsx1* gene during the FECA to LECA transition could have enabled pre-LECA cells to undergo genome expansion, explore syncytial forms (*48*) and evolve into mononucleated cells fully equipped for meiosis and gamete fusion (*49*). Our findings raise the possibility that gamete fusion is the product of over two billion years of evolution of this ancient archaeal cell fusion machine.

## Acknowledgments

We thank Sonja-Verena Albers, Dan Cassel, Peter Walter, Alejandro Colman-Lerner, Uri Gophna, Yael Iosilevskii, Shahar Lavid, and members of our laboratories for discussion and comments on the manuscript; Jose Flores for advice on searches of fusexins in metagenomes; Olivier Gascuel for discussions on phylogenetics; Pablo Dans for advice on molecular dynamics; Yoav Henis for the GPI-BHA plasmid. Part of the computations for this work was performed at the Vital-IT Center for high-performance computing of the SIB Swiss Institute of Bioinformatics.

## Funding

Comisión Sectorial de Investigación Científica grant CSIC I+D-2020-682, Uruguay (HR, MG)

Fondo para la Investigación Científica y Tecnológica grant, Argentina (PICT-2017-0854 (PSA)

Beca de Doctorado Consejo Nacional de Investigaciones Científicas y Técnicas, Argentina (DM)

Knut and Alice Wallenberg Foundation grant 2018.0042, Sweden (LJ)

Novartis Foundation for Medical-Biological Research grant 17B111, Switzerland (CD)

Swedish Research Council grants 2016-03999 and 2020-04936 (LJ)

Swiss Leading House for the Latin American Region (CD and PSA)

Swiss National Science Foundation grant 183723 (CD)

Israel Science Foundation grants 257/17, 2462/18, 2327/19, and 178/20 (BP)

European Union’s Horizon 2020 research and innovation programme under the Marie Skłodowska-Curie grant agreement No 844807 (NGB)

FOCEM, Fondo para la Convergencia Estructural del Mercosur grant COF 03/11 (MG)

Programa de Desarrollo de las Ciencias Básicas, Uruguay (HR, MG, and ML)

## Author contributions

BP conceived the experiments; performed some imaging work; designed, supervised and analyzed cell fusion experiments. CD helped devise analysis strategies for k-mer and phylogenetic analysis. CV designed, performed and analyzed cell fusion assays. DdeS collected X-ray data, took part in structural analysis and validated metal substructure. DM carried out deep homology detection of Fsx1; designed gene content analysis strategies and phylogenetic analysis, collected sequence data; performed k-mer, functional, structural and phylogenomics analyses, built homology models, coded analysis routine pipelines. HR supervised the bioinformatics part of the work, estimated relative acquisition times, designed and performed, non-reversible substitution models analyses, structural phylogenetic analyses and phylogenomic surveys. KF made constructs of Fsx1 mutants. KT and JJ generated AlphaFold2 models of monomeric Fsx1. LJ supervised the biochemical and structural part of the work; collected X-ray data; solved the Fsx1 structure, refined and analyzed it; and generated AlphaFold-Multimer models of trimeric Fsx1. MG designed bioinformatic strategies, supervised bioinformatic aspects of the work, analyzed sequence and structural data. ML performed IME synteny analyses, phylogenetic and phylogenomic surveys. NGB carried out live imaging and surface biotinylation experiments, assisted with the preparation and design of plasmids; analyzed data. PSA supervised the bioinformatics part of the work, analyzed data. SN expressed, purified and crystallized Fsx1; performed SEC-MALS experiments; analyzed SAXS data; collected X-ray data and took part in structure determination, model building and structure analysis. XL carried out immunofluorescence and western blots for archaeal fusexins in mammalian cells, designed and constructed plasmids, performed imaging work. DM, SN, XL, NGB, MG, HR, PSA, LJ and BP made figures and tables. DM, SN, XL, MG, HR, PSA, LJ and BP wrote the manuscript. All authors reviewed the manuscript.

## Data and materials availability

Crystallographic structure factors and atomic coordinates have been deposited in the Protein Data Bank under accession code 7P4L. All relevant codes, notebooks and datasets necessary for: HHblits (v3.3.0) and HMMER searches and comparisons (Fig. 1A, fig. S1 and data S1); raw data and statistical information for functional analyses (Fig. 3, fig. S2 and data S4); sequences of *fsx1* genes synthesized (Data S5); K-mer spectra analyses (Fig. 4A and fig. S13); IMEs clustering, content and synteny analyses (Fig. 4B, figs. S14, S15 and S16, data S2 and S3), protein structure- and sequence-based comparisons (Fig. 4C and fig. S17), are available at https://github.com/DessimozLab/Archaeal-Fusexins.

## Supplementary Materials

Materials and Methods

Figs. S1-S19

Tables S1-S7

References (50-145)

Movies S1-S2

Data S1-S5

