## Supplementary methods, figures and tables for "Discovery of archaeal Fusexins homologous to eukaryotic HAP2/GCS1 gamete fusion proteins"

**This PDF file includes:**

**Materials and Methods**

***Fusexin genes in Archaea***

**Initial fusexin search using structurally-guided MSAs**

###### HMM-based distance matrices

###### Metaclust database search pipeline

##### *Fsx1 is a structural homolog of HAP2/GCS1*

###### DNA constructs

###### Protein expression and purification

###### Size exclusion chromatography-multiangle light scattering (SEC-MALS)

###### Small-angle X-ray scattering (SAXS)

###### Crystallization and X-ray diffraction data collection

###### Data reduction and non-crystallographic symmetry analysis

###### Structure determination by molecular replacement with AlphaFold2 models

###### Model building, refinement and validation

###### Sequence-structure analysis

###### Structural modeling of trimeric Fsx1

##### *Fsx1 can fuse eukaryotic cells and Structure-function analysis of Fsx1*

###### Cells and reagents

###### Immunofluorescence

###### Western blots

###### Content mixing assays with immunofluorescence

**Cell fusion assay by content mixing with nuclear and cytoplasmic markers**

###### Live imaging of fusing cells

###### Surface biotinylation

###### Data analysis

###### Statistical tests

##### *Fsx1s are ancient fusogens associated with integrated mobile elements*

###### Integrated Mobile Element (IME) identification by k-mer spectra analysis and

###### comparative genomics

###### IME homology analyses

**Supplementary Figs. S1 - S19**

**Supplementary Tables S1 - S7**

**Captions for Movies S1 - S2**

**Captions for Data S1 - S5**

**Materials and Methods**

###### No statistical methods were used to predetermine sample size. The experiments were not randomized. All software versions numbers and databases used are available through the project’s GitHub: https://github.com/DessimozLab/Archaeal-Fusexins

***Fusexin genes in Archaea***

###### Initial fusexin search using structurally-guided MSAs

HMMs were prepared using structurally guided multiple sequence alignments (MSAs) of known eukaryotic HAP2 sequences (ectodomains only). Structural MSAs were derived using I-TASSER (v5.1) (*50*) generated models of HAP2 homologues for *Erythranthe guttata* (A0A022QRC8), *Phytomonas* sp. isolate EM1 (W6KUI1), *Plasmodium falciparum* (A0A1C3KGX6), *Chlorella variabilis* (E1Z455) and the HAP2 crystal structures for *Chlamydomonas reinhardtii* (PDB 6E18 (*51*) and 6DBS (*18*)) and *Arabidopsis thaliana* (PDB 5OW3 (*17*)).

Searches for fusexin homologues using structurally guided MSAs were performed for 3 iterations on the Uniclust database (*52*) using default HHblits (v3.3.0) parameters (*53*).

###### HMM-based distance matrices

A taxonomically representative list of known viral and eukaryotic fusexin homologues, covering major lineages, was manually curated. A MSA was built for each homologue by using the sequence as a query on the Uniclust database with HHblits (*53*) for 3 iterations (see Supplementary Information). This set of MSAs was compiled into an HH-suite (v3.3.0) database and each MSA was used as a query against this database to establish a profile-based distance matrix using the probability of homology (Fig. 1A and fig. S1).

###### Metaclust database search pipeline

We searched the Metaclust (*54*) dataset (nr50) using an HMM made of Fsx1 sequences found in PCGs and MAGs (data S1; Auxiliary Supplementary Materials). Fsx1 sequences were aligned using ClustalO (v1.2.2) (*55*) with default settings for 3 iterations and the resulting MSA was used as a query with hmmsearch (hmmer suite v3.3.2 ) (*56*) against the Metaclust50 dataset (*54*). All returned sequences with an E-value < 0.0001 with a match length greater than 100 residues were selected for further analysis. PSIBLAST (v2.12.0) was also used on the Metaclust (nr90) with Fsx1 sequences found in PCGs and MAGs with default parameters for 3 iterations. All returned sequences with an E-value < 0.0001 and an alignment length greater than 100 were added to the pool of candidates. Manual curation was performed using membrane protein topology predictor TOPCONS(v2.0) (*57*) and distant homology searches using HHBlits (*58*) against the PDB70.

##### *Fsx1 is a structural homolog of HAP2/GCS1*

###### DNA constructs

Ten archaeal genes were synthesized (GenScript) and cloned into pGene/V5-His vectors (table S5). Details of nucleotides used for synthesis and protein sequences are described in data S5.

For structural studies, a synthetic gene fragment encoding the extracellular region of a metagenomic Fsx1 ORF (IMG genome 3300000868, scaffold JGI12330J12834_ 1000008, ORF 8; data S1) (GenScript) was subcloned by PCR in frame with the 5’ chicken Crypα signal peptide- and 3’ 8xHis-tag-encoding sequences of pLJ6, a mammalian expression vector derived from pHLsec3(*59*). The protein construct that yielded the final high-resolution dataset included residues D25-S535 and contained a T369C substitution, introduced by PCR mutagenesis with the aim of facilitating heavy atom derivatization for experimental phasing. Oligonucleotides were from Sigma-Aldrich or IDT and all constructs were verified by DNA sequencing (Eurofins Genomics or Macrogen).

To generate pCI::GFPnes plasmid (see list of plasmids in table S6), an oligo DNA encoding for the nuclear export signal (LQKKLEELELD) was cloned downstream the region encoding EGFP of the pCAGIG plasmid using the enzyme BsrGI. Then, the GFPnes coding sequence was amplified, cut with BmgBI and BglII and used to replace the H2B-GFP coding sequence of the pCI::H2B-GFP plasmid (see list of primers in table S7). Fsx1-V5, AtHAP2-V5(*3*), EFF-1-V5, VSV-G (*11*) and other archaeal fusexins (NaFsx1, HQ22Fsx1, HnFsx1) were subcloned into corresponding pCI::H2B-RFP/H2B-GFP/GFPnes vectors separately. For mutagenesis of Fsx1, i) Fsx1-ΔFL-AG_4_A: The mutation of Y142A, Y149A and four glycines inserted between them were achieved using PCR with overlapping primers. ii) Fsx1-ΔDIV-EFF-1 stem: The stem region of EFF-1(E510-D561) was amplified from pGene::EFF-1-V5 and fused to the upstream and downstream regions of Fsx1-DIV with overlapping primers. iii) Fsx1ΔTMs→EFF-1 TM: The TM and cytoplasmic tail of EFF-1 (I562-I658) were amplified from pGene::EFF-1-V5 and fused to the ectodomain of Fsx1 to replace its TMs. iv) Fsx1ΔTMs→GPI: The Fsx1 TMs were replaced with the carboxy-terminal 37 amino acids of decay accelerating factor (DAF) which confer the signal for GPI anchor (*60*). Similarly, the TM and cytoplasmic tail of AtHAP2 were replaced with the GPI signal from DAF to get AtHAP2ΔTM→GPI. All mutants were ligated into pCI::H2B-RFP and pCI::GFPnes vectors for mixing assay. Additional details are found in tables S6 and S7.

###### Protein expression and purification

HEK293T cells (*61*) were transiently transfected using 25 kDa branched polyethyleneimine and cultured in DMEM media (Invitrogen) supplemented with 2% (v/v) foetal bovine serum (Biological Industries). 90-96 hours after transfection, the conditioned media from HEK293T cells was harvested, 0.2 µm-filtered (Pall) and adjusted to 20 mM Na-HEPES pH 7.8, 2.5 M NaCl, 5 mM imidazole. 10 ml Ni Sepharose excel beads (GE Healthcare) pre-equilibrated with immobilized metal affinity chromatography (IMAC) buffer (20 mM Na-HEPES pH 7.8, 2.5 M NaCl, 10 mM imidazole) were added to 1 L adjusted conditioned media and incubated overnight at 4ºC. After washing the beads with 100 column volumes IMAC buffer, captured Fsx1_E_ was batch-eluted with 30 mL 20 mM Na-HEPES pH 7.8, 2.5 M NaCl, 500 mM imidazole and concentrated with 30 kDa-cutoff centrifugal filtration devices (Amicon). The material was then further purified by SEC at 4ºC, using an ÄKTAfplc chromatography system (GE Healthcare) equipped with a Superdex 200 Increase 10/300 GL column (GE Healthcare) pre-equilibrated with 20 mM Na-HEPES pH 7.8, 2.5 M NaCl. Peak fractions were pooled and concentrated to 5 mg mL^-1^ (fig. S3, A and B).

###### Size exclusion chromatography-multiangle light scattering (SEC-MALS)

Purified Fsx1_E_ samples (120-150 µg) were measured using an Ettan LC high-performance liquid chromatography system with a UV-900 detector (Amersham Pharmacia Biotech; λ = 280 nm), coupled with a miniDawn Treos MALS detector (Wyatt Technology; λ = 658 nm) and an Optilab T-rEX dRI detector (Wyatt Technology; λ = 660 nm). Separation was performed at 20ºC using a Superdex 200 Increase 10/300 GL column (GE Healthcare) with a flow rate of 0.5 mL min^−1^ and mobile phases consisting of 20 mM Tris-HCl pH 8.5, 150 mM NaCl (normal salt condition) or 20 mM Tris-HCl pH 8.5, 2.0 M NaCl and 0.2 M CaCl_2_ (high salt condition) (fig. S3C). Data processing and weight-averaged molecular mass calculations were performed using ASTRA (v7.1.3) (Wyatt Technology). BSA (150 µg) was used as a control.

###### Small-angle X-ray scattering (SAXS)

SAXS experiments were performed at beamline BM29 of the European Synchrotron Radiation Facility (ESRF) ((*62*), using Fsx1_E_ (4.5 mg mL-1) in 20 mM Na-HEPES pH 7.8, 150 mM NaCl. Sample delivery and measurements were performed using a 1 mm thick quartz capillary, which is part of the BM29 BioSAXS automated sample changer unit (*63*). Data were collected at 1 Å wavelength in 10 frames of 1 s at 20ºC, using an estimated beam size of 1 mm x 100 µm; buffer blank measurements were carried out under the same conditions, both before and after sample measurement. Data were averaged and subtracted using PRIMUS (*64*) from the ATSAS package (v3.0.3) (*65*), which was also used to calculate the pair-distance distribution function, as well as the radius of gyration and the Porod volume. Theoretical scattering curves for monomeric and trimeric Fsx1_E_ were calculated and compared with the experimental data using CRYSOL (ATSAS v3.0.3) (*66*). *Ab initio* envelope reconstruction was performed using the program DAMMIF (ATSAS v3.0.2) ((*67*), resulting in twenty models that were superimposed and averaged with DAMAVER (ATSAS v3.0.2) (*68*). Chain A of the refined Fsx1_E_ model was fitted into the SAXS reconstruction using UCSF ChimeraX (v1.2.4) (*69*) (fig. S3D).

###### Crystallization and X-ray diffraction data collection

Two similar initial hits obtained from extensive screening using a mosquito crystallization robot (TTP Labtech) were manually optimized by setting up vapour diffusion experiments at 20ºC in 24-well plates. To grow diffraction-quality crystals, 1 µl purified Fsx1_E_ was mixed with 1 µL 23% (w/v) PEG 4000, 0.1 M Tris-HCl pH 8.5, 0.2 M CaCl_2_ and equilibrated against 1 mL of the same solution. Rhomboidal plates of Fsx1_E_ grew in 1-3 months from protein precipitate that appeared after overnight equilibration of the crystallization drops (fig. S3E). For data collection, specimens were freed from the precipitate by micromanipulation with MicroMounts (MiTeGen) and flash frozen in liquid nitrogen. More than a hundred crystals were screened at beamlines ID23-1 of the ESRF (*70*) and I04 of Diamond Light Source, yielding datasets of highly variable quality. The final X-ray diffraction dataset at 2.3 Å resolution was collected at ESRF ID23-1.

###### Data reduction and non-crystallographic symmetry analysis

Datasets were processed in space group *C*2 with XDS (version Jan 31, 2020 BUILT=20200417) (*71*) (table S2). By revealing a strong non-origin peak at chi=120 (fig. S3F), self rotation functions calculated with MOLREP (v11.7.03) (*72*) or POLARRFN (v7.1.010) (*73*) clearly indicated the presence of three-fold non-crystallographic symmetry (NCS) within the asymmetric unit of the centred monoclinic crystals. Combined with Matthews coefficient calculations (*74*, *75*), this strongly suggested that Fsx1_E_ crystallized as a homotrimer.

###### Structure determination by molecular replacement with AlphaFold2 models

Multiple attempts to experimentally determine the structure of Fsx1_E_ using a variety of heavy atoms failed, probably because the high-salt mother liquor composition hindered heavy atom binding. Because molecular replacement (MR) with HAP2-derived homology models also failed, we took advantage of the recent significant advances in protein 3D structure prediction using machine learning (*76*) to phase the data by MR (*77*) (fig. S4). To do so, we used AlphaFold2 (*19*) (with default monomer prediction parameters) to generate five independent models of Fsx1 ectodomain residues D25-S535, with per-residue pseudo-B factors corresponding to 100-(per-residue confidence (pLDDT (*19*))). These models had relative root-mean-square deviations (RMSD) of 1.4-3.3 Å, or 0.7-1.9 Å after excluding 26 C-terminal residues predicted with low-confidence. Initial attempts to solve the structure with Phaser (v2.8.3) (*78*), using an ensemble including these models (further truncated to Q453, the predicted C-terminal end of domain III), yielded 4 solutions (with top Log Likelihood Gain (LLG) 188, final Translation Function Z score (TFZ) 9.6) that were retrospectively correct in terms of domain I/II placement, but completely wrong in the positioning of domain III. Because of the latter, automatic refinement of these solutions did not progress beyond R_free_ ~0.53. On the other hand, a parallel consecutive search for three copies of a domain I/II ensemble (D25-A335; RMSD 0.3-0.9 Å) followed by three copies of domain III (P350-Q453; RMSD 0.1-0.3 Å), using a model RMSD variance of 1 Å, yielded a clear single solution (LLG 876, TFZ 23.1) that could be automatically refined to initial R 0.45, R_free_ 0.46.

Remarkably, although a single copy of domain 3 corresponds to only 7% of the total scattering mass in the asymmetric unit of the Fsx1_E_ crystal, the very high accuracy of its AlphaFold2 model (reflected by *a posteriori*-calculated global RMSD and Distance Test Total Score (GDT_TS) of 0.7 Å and 97.6, respectively) allowed Phaser to also find a correct MR solution using just this part of the structure. Specifically, a consecutive search for three copies of the domain resulted in a trimeric model with LLG 275 and TFZ 15.1, which could be refined to starting R 0.51, R_free_ 0.51.

Also worth mentioning is the observation that the same domain I/II + domain III MR strategy used to phase the 2.3 Å resolution data could also be successfully applied to an initial dataset at much lower resolution (3.5 Å, with outer shell mean I/σI 0.6 and CC_1/2_ 0.31); in this case, the Phaser LLG and TFZ values for the solution were 361 and 13.5, respectively, and initial automatic refinement of the corresponding model yielded R 0.44, R_free_ 0.48.

###### Model building, refinement and validation

The initial model of Fsx1_E_ was first automatically rebuilt using PHENIX AutoBuild (v1.19.2) ((*79*) (1083 residues; R 0.34, R_free_ 0.38) and then significantly improved with the machine-learning-based sequence-docking method of ARP/wARP (v8.0 patch 1) ((*80*), as implemented in CCP4 (v7.1.012) ((*73*) (1390 residues; REFMAC (v5.8.0267) ((*81*) R 0.23). The resulting set of coordinates was subsequently subjected to alternating cycles of manual rebuilding with Coot (versions 0.8.9.3-0.9.6) ((*82*)/ISOLDE (v1.1.0) ((*83*) and refinement with phenix.refine (versions 1.19.2 and dev-4282) ((*84*), using torsion-based NCS restraints and three Translation-Libration-Screw-rotation groups per chain. Metal ions were assigned based on electron density level; difference Fourier maps generated using alternative atom types; correspondence with peaks in phased anomalous difference maps, calculated with PHENIX (v1.19.2) ((*85*) or ANODE (v2013/1) ((*86*) from data collected at low energy; and coordination properties (*87*). Protein geometry was validated using MolProbity (v4.5.1) ((*88*) (table S2).

###### Sequence-structure analysis

Transmembrane helices were predicted using TMHMM (v2.0) ((*89*). GDT_TS scores were calculated using LGA (v09/2019) ((*90*) and structural similarities were assessed with Dali (v5) ((*91*) and PDBeFold (v2.59) ((*92*). Secondary structure was assigned using DSSP (v4.0-67) ((*93*). Subunit interfaces were analyzed using PDBsum (*94*), PIC (*95*) and PDBePISA (v1.52) ((*96*). Molecular charge was calculated using the YASARA2 force field of YASARA Structure (v.21.7.1) ((*97*) and electrostatic surface potential calculations were performed with PDB2PQR (v2.1.2) ((*98*) and APBS (v1.5) ((*99*), via the APBS Tools plugin of PyMOL (v2.4.2) (Schrödinger, LLC). Mapping of amino acid conservation onto the 3D structure of Fsx1_E_ was carried out by analyzing a sequence alignment of archaeal homologues with ConSurf (*100*). Structural figures were generated with PyMOL ( v2.5.2 ) and assembled in Illustrator CC 2020 (Adobe).

###### Structural modeling of trimeric Fsx1

Models of homotrimeric Fsx1 were generated using a local copy of AlphaFold-Multimer (v2.1.1) ((*101*), installed using the open‐source code and instructions available at <https://github.com/deepmind/alphafold>.

###### *Fsx1 can fuse eukaryotic cells and Structure-function analysis of Fsx1*

###### Cells and reagents

Baby Hamster Kidney (BHK-21) cells (kindly obtained from Judith White, University of Virginia) were maintained in DMEM supplemented with 10% FBS (Biological Industries), 100 U/ml penicillin, 100 µg/ml streptomycin (Biological Industries), 2 mM L-glutamine (Biological Industries), 1 mM sodium pyruvate (Gibco), and 30 mM HEPES buffer, pH 7.3, at 37°C with 5% CO_2_. Transfections were performed using Fugene HD (Promega) or jetPRIME (Polyplus) according to the manufacturer’s instructions.

###### Immunofluorescence

BHK cells were grown on 24-well tissue-culture plates with glass coverslips. Permeabilized cells were fixed with 4% paraformaldehyde (EM grade, Bar Naor, Israel) in PBS, followed by incubation in 40 mM NH_4_Cl to block free aldehydes, permeabilized in 0.1% Triton X-100 in PBS and blocked in 1% FBS in PBS. After fixation, the coverslips were incubated 1 h with mouse anti–V5 antibody (Invitrogen, 1:500) and 1 h with the secondary antibody which was donkey anti–mouse coupled to Alexa Fluor 488 (Invitrogen, 1:500). Alternatively, for immunofluorescence without permeabilization, cells were blocked on ice in PBS with 1% FBS for 20 minutes, and then stained with Monoclonal ANTI-FLAG M2 antibody (Sigma, 1:1000) on ice for 1h. After anti-FLAG staining, cells were washed and fixed with 4% PFA in PBS. Cells were blocked again and stained with the secondary antibody (donkey anti–mouse coupled to Alexa Fluor 488; Invitrogen) diluted 1:500 in PBS for 1 h. In all cases, nuclei were stained with 1 µg/ml DAPI. Images were captured using a Nikon Eclipse E800 with a 60X/1.40 Plan Apochromat objective and an optical zoom lens (Nikon) using a Hamamatsu ORCA-ER camera controlled by Micro-Manager (v1.4.22) software (*102*) (fig. S10D).

###### Western blots

24 h post-transfection, cells were treated with Lysis Buffer (50 mM Tris-HCl pH 8.0, 100 mM NaCl, 5 mM EDTA, 1% Triton X-100 supplemented with chymostatin, leupeptin, antipain and pepstatin) on ice for 10 min. After 10 min centrifugation at 14,000 rpm at 4 °C, supernatants of lysates were mixed with reducing sample buffer (+ DTT) and incubated 5 min at 95°C. Samples were loaded on a 10% SDS-PAGE gel and transferred to PVDF membrane. After blocking, membranes were incubated with primary antibody anti–V5 mouse monoclonal antibody (1:5,000; Invitrogen) or anti-actin (1:2,000; MP Biomedicals) at 4 °C overnight and HRP-conjugated goat anti-mouse secondary antibody 1 h at room temperature. Membranes were imaged by the ECL detection system using FUSION-PULSE.6 (VILBER).

###### Content mixing assays with immunofluorescence

BHK-21 cells at 70% confluence were transfected (using JetPrime; Polyplus at a ratio of 1:2 DNA:transfection reagent) with 1 µg pCI::Fsx1-V5::H2B-eGFP, pCI::Fsx1-V5::H2B-RFP, pCI::AtHAP2-V5::H2B-eGFP, pCI::AtHAP2-V5::H2B-RFP, respectively. Control cells were co-transfected with pCI::H2B-eGFP and pRFPnes or pCI::H2B-RFP and pRFPnes. 4 h after transfection, the cells were washed 4 times with DMEM with 10% serum (Invitrogen), 4 times with PBS and detached using Trypsin (Biological Industries). The transfected cells were collected in Eppendorf tubes, resuspended in DMEM with 10% serum, and counted. Equal amounts of H2B-RFP and H2B-eGFP cells were mixed and seeded on glass-bottom plates (12-well black, glass-bottom #1.5H; Cellvis) and incubated at 37^o^C and 5% CO_2_. 18 h after mixing, 20 µM 5-fluoro-2’-deoxyuridine (FdUrd) was added to the plates to arrest the cell cycle and 24 h later, the cells were fixed with 4% PFA in PBS and processed for immunofluorescence. To assay mixed cells and detect the transfected proteins (Fsx1-V5 or AtHAP2-V5), we stained cells with anti-V5 mAb (Life Science). The secondary antibody was Alexa Fluor 488 goat anti-mouse, with 1 µg/ml DAPI (*3*). Micrographs were obtained using wide-field illumination using an ELYRA system S.1 microscope (Plan-Apochromat 20X NA 0.8; Zeiss) and recorded with a iXon+ EMCCD camera (Andor). The GFP + RFP mixing index was calculated as the number of Red and Green nuclei in mixed cells out of the total number of nuclei of fluorescent cells in contact (Fig. 3B).

**Cell fusion assay by content mixing with nuclear and cytoplasmic markers**

For the unilateral setup, BHK-21 cells were transfected (as explained above) with 1 µg pCI::GFPnes; pCI::Fsx1-V5::GFPnes; 0.25 µg pCI::EFF-1-V5::GFPnes; 1 µg pCI::VSV-G::GFPnes in respective 35 mm plates. The cells were incubated, washed, and mixed with pCI::H2B-RFP (empty vector) transfected cells (Fig. 3E). For evaluating the mutants, BHK-21 cells were transfected with 1 µg pCI::Fsx1-V5::GFPnes or pCI::Fsx1-V5::H2B-RFP or the plasmids encoding for each mutant: ΔFL→AG_4_A, ΔDIV→EFF-1 stem, ΔTMs→EFF-1 TM, Fsx1ΔTMs→GPI or AtHAP2ΔTM→GPI (Fig. 3I). Empty pCI::GFPnes or pCI::H2B-RFP were used as negative controls. 4 h after transfection, the cells were washed, counted, mixed, and incubated as previously described. In all cases, 18 h after mixing, 20 µM FdUrd was added to the plates, and 24 h later, the cells were fixed with 4% paraformaldehyde diluted in PBS. Nuclei were stained with 1 µg/ml DAPI. Images were obtained using wide-field illumination with an ELYRA system S.1 microscope as described above.

The GFP + RFP mixing index was calculated as the number of nuclei in mixed cells, green cytoplasm (GFPnes) with red (H2B-RFP) and blue (DAPI) nuclei out of the total number of nuclei in fluorescent cells in contact (Fig. 3, F and J). The multinucleation indexes were defined as the ratio between the number of nuclei in multinucleated cells (Nm) and the total number of nuclei in multinucleated cells and expressing cells that were in contact (Nc) but did not fuse, using the following equation: % multinucleation = Nm/(Nc+Nm)×100. The percentage of multinucleation was calculated for GFPnes cells with RFP and DAPI nuclei. For the unilateral assay, multinucleation was determined as the ratio between the number of nuclei in multinucleated green cells and the total number of nuclei in green multinucleated cells and GFPnes expressing cells that were in contact but did not fuse (Fig. 3F).

###### Live imaging of fusing cells

BHK cells were plated on 15 mm glass bottom plates (Wuxi NEST Biotechnology Co., Ltd.) and transfected with 1 µg pCI::Fsx1-V5::H2B-GFP together with 0.5 µg myristoylated-mCherry (myr-palm-mCherry; kindly provided by Valentin Dunsing and Salvatore Chiantia (*103*)). 18 h after transfection, the cells were incubated with 2 μg/ml Hoechst dye for 10 min at 37°C and washed once with fresh medium. Time-lapse microscopy to identify fusing cells was performed using a spinning disc confocal microscope (CSU-X; Yokogawa Electric Corporation) with an Eclipse Ti and a Plan-Apochromat 20X (NA, 0.75; Nikon) objective. Images in differential interference contrast and red channels were recorded every 4 min in different positions of the plate using high gain and minimum laser exposure. Time lapse images were captured with an iXon 3 EMCCD camera (Andor Technology). After 5 h, confocal z-series, including detection of the DAPI channel, were obtained to confirm the formation of multinucleated cells. Image analyses were performed in MetaMorph (v7.8) (Molecular Devices) and ImageJ (v1.53c) (*104*) (National Institutes of Health).

###### Surface biotinylation

Proteins localizing on the surface were detected as previously described (*3*). Briefly, BHK cells were transfected with 1 µg pCAGGS, pCAGGS::EFF-1-V5, pCAGGS::Fsx1-V5, pCAGGS::ΔFL→AG_4_A-V5 ,pCAGGS::ΔDIV→EFF-1 stem-V5 or pCAGGS::Fsx1ΔTMs→EFF-1 TM-V5. 24 h later, cells were washed twice with ice-cold PBS^2+^ (with Ca^2+^ and Mg^2+^) and incubated with 0.5 mg/ml EZ-Link Sulfo NHS-Biotin (Thermo Fisher Scientific) for 30 min on ice. The cells were washed four times with ice-cold PBS^2+^, once with DMEM with 10% FBS (to quench residual biotin), followed by two more washes with PBS^2+^. To each plate 300 µl of Lysis Buffer supplemented with 10 mM iodoacetamide were added and the cells detached using a scrapper. The insoluble debris was separated by centrifugation (10 min at 21,000 *g*), and the lysate was mixed with NeutrAvidin Agarose Resin (Thermo Fisher Scientific) and 0.3% SDS. After an incubation of 12 h at 4°C the resin was separated by centrifugation (2 min at 21,000 *g*), washed three times with lysis buffer and then mixed with SDS-PAGE loading solution with freshly added 5% b-mercaptoethanol and incubated 5 min at 100°C. After pelleting by centrifugation, the samples were separated by SDS-PAGE gel and analyzed by Western blotting as described above using anti–V5 mouse monoclonal antibody. Loading was controlled using anti-actin C4 monoclonal (1:2,000; MP Biomedicals).

###### Data analysis

Counting of content mixing and multinucleation was made blind for the experiments included in Fig. 3, F and J. Interobserver error was estimated for counting of multinucleated cells, cells in contact, and content-mixing experiments: the differences in percentages of multinucleation and content mixing obtained by two observers was <10%.

###### Statistical tests

Results are presented as means ± SEM. For each experiment we performed at least three independent biological repetitions. To evaluate the significance of differences between the averages we used one-way ANOVA as described in the legends (GraphPad Prism 9).

***Fsx1s are ancient fusogens associated with integrated mobile elements***

##### Integrated Mobile Element (IME) identification by k-mer spectra analysis and comparative genomics

Comparison between close species with presence (*fsx1+)* or absence (*fsx1-)* of archaeal fusexins to detect insertion sites was done performing sequence similarity searches in complete genomes from the closest relatives available in the PATRIC database (*105*) (fig. S12 and table S1). Coordinates of *fsx1*-containing IMEs present in pure culture genomes (PCGs) are annotated in table S4**.**

Among different methodologies that rely on DNA composition to identify horizontally transferred genomic regions (*106*), k-mer spectrum analysis is a standard tool for this purpose (*107*, *108*). Normalized k-mer spectra for DNA sequences of arbitrary length were generated by counting occurrences of all k-mers and normalizing by the total amount of words counted. k-mer sizes from 3 bp to 8 bp were tested with no effect on results. A length of 4 bp was selected. To detect possible horizontally transferred regions, an average spectrum for each genome was calculated. A spectrum was calculated for a sliding window of 1 kb using 500 bp steps and subtracted from the genomic average at each window position **(**Fig. 4A and fig. S13**)**. The absolute value of the difference between the genomic average and window spectra is represented over the entire genome. Gaussian mixture models using two distributions were fitted (*109*) to the k-mer content of all windows, to classify these as belonging to either the core genome or transferred elements. This deviation in k-mer spectra has been explored in the context of the archaeal mobilome and contains information on the ecological niche and evolutionary history of DNA sequences (*110*).

### **IME gene content and homology analyses**

We followed the pipeline depicted in figure S14. Briefly, PCGs’ IMEs were determined by a combination of k-mer spectra and genomic alignments (see table S4). We initially inspected *fsx1*-containing scaffolds and kept only sequences that were 20 kb or longer for downstream analyses. We generated an enriched annotation for each IME. Then, we obtained an initial set of groups of homologous sequences, and each of these groups was enriched by means of HMM searches. Subsequently, the enriched homology groups showing similarity between them, as judged by HMM-HMM comparisons, were collapsed into unique groups.

In detail, first, we re-annotated the identified mobile elements combining the corresponding segment of the PATRIC (*105*) GFF annotation file with in-house ORF predictions (minimum ORF length of 30 nucleotides, option by default). ORF inference was done by means of getorf of the EMBOSS package v6.6.0.0(*111*), specifying genetic code by Table 11 (Bacteria and Archaea) and other parameters running by default. The similarity of inferred ORFs and annotated features in these mobile elements (i.e. features in their GFF annotation file) was established by means of BLASTP reciprocal searches (*112*). We kept all the predicted ORFs and homologs that were at least annotated in one genome, in this way we tried to recover missanotated conserved ORFs.

Initial sets of homologs were generated with get_homologs v20210305 (*113*). Sequence identity and query coverage thresholds were set to 35% and 70%, respectively. In-paralogues were not allowed within these groups (option ‘-e’), and remaining parameters were run by default.

HMM profiles were constructed for each homolog group. To this aim, homologous sequences were retrieved for members of each group from the UniRef50 database (*114*) with jackhmmer (HMMER package v3.1b2; <http://hmmer.org> (*56*)) running with one iteration (‘-N 1’ parameter). MSAs were then generated for each group and its relevant hits with MAFFT (*115*) (v7.310) running under ‘--auto’ parameter, and HMMs were created with hmmbuild (HMMER (*56*)). Homolog groups were enriched by means of HMM searches with hmmsearch (HMMER (*56*)), using each HMM as a query against a database comprising all predicted ORFs described above. Hits showing an e-value < 1e-10 and covering at least 50% of the HMM were added to the groups.

Enriched homology groups showing homology were collapsed. For this purpose, HMM-vs-HMM comparisons were performed with HHalign v3.3.0 from the HHsuite (*116*). A graph was created with the Python library networkx v2.5.1, each node being an enriched group of homologs. An edge was established between nodes if their HMM-HMM alignment was significant (i.e. e-value < 1e-10, HMM coverage of longest HMM >= 50%). Groups of interconnected nodes were established with the ‘connected_components()’ routine, creating a collapsed homology group in each case.

Finally, we assessed the gene content similarity between mobile elements using a Jaccard Index based on the homology groups defined above. Usual Jaccard index of two sets is defined as

(# of the intersection)/(# of the union). In this case:

$J (MEA,MEB) =\frac{Nhomology groups shared between ME A \& ME B}{N homol. groups MEA + N homol. groups MEB - Nhomol. groups shared between ME A \& ME B}$

We performed a hierarchical clustering of the MEs based on a distance matrix obtained from the pairwise Jaccard Indexes (distance(A,B) = 1 - J_A,B_). This was done in Python with seaborn v0.11.1 (*117*), employing the clustermap function. A subset of 11 mobile elements (fig. S15, in red), which included ME from PCGs and JGI12330J12834-1000008 (data S1 to S3), was selected for synteny conservation analysis. Plots depicting synteny in gene content between homolog groups were generated employing the MCscan tool (*118*).

HMMER (*56*) and Pfam (*119*) were used on default parameters to assign domains and their associated arCOG (*120*, *121*) identifiers to ORFs (data S2).

These analyses, including collapsed clusters, can be found at <https://github.com/DessimozLab/Archaeal-Fusexins>.

###### Sequence and Structure phylogenies

Maximum likelihood phylogenetic trees were generated with sequences aligned with MAFFT (v7.310) (*115*) (L-INS-i option) as input for IQ-TREE (v1.6.12) (*122*) and selecting the best evolutionary model with ModelFinder (*123*). Homology trimeric models of archaeal homologs of Fsx1_E_ (fig. S17) were built with MODELLER (v10.2) (*124*) using our crystal structure as template.

Protein folds preserve deeper evolutionary signals than sequences (*125*–*127*). Fsx1 models and crystal structures of Fsx1_E_ and eukaryotic and viral fusexins were all-vs-all compared with FATCAT (v2.0) (*128*) to establish their structural distances between them. The following experimental crystal structures from other works were used: Flavivirus E: West Nile virus (2I69) (*129*); Dengue virus serotype 1 (4GSX) (*130*); Alphavirus E1: Semliki Forest virus (1RER) (*131*); Chikungunya virus (3N43) (*132*)*; C. elegans* EFF-1 (4OJC) ((*10*); Bunyavirus Gc Rift Valley fever virus (6EGU) (*133*); eukaryotic HAP2/GCS1 from *A. thaliana* (5OW3) (*17*) and *C. reinhardtii* (6E18) (*51*). The PDB files produced by flexible alignment with FATCAT (v2.0) were compared with TMalign (v20210224) (*134*) to build a TM_score_ (*135*) distance matrix (distance = 1-TM_score_). This distance matrix was the basis to compute a minimum evolution tree with FastME (v2.1.6.2) (*136*) on default parameters.

### **Supplementary Figures**

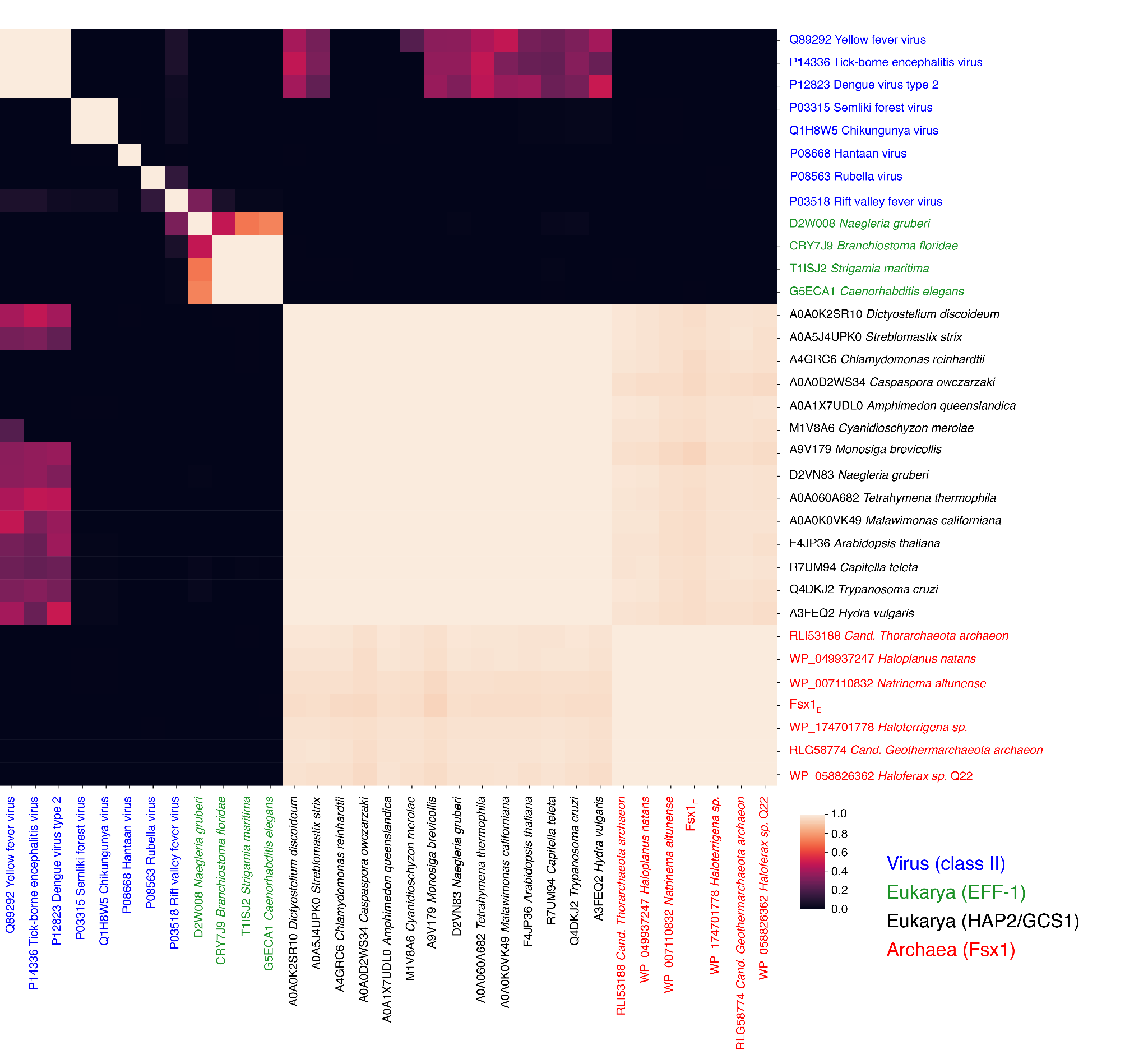

**Fig. S1. Fsx1s are members of the fusexins superfamily.** HMM homology probabilities of fusexins and archaeal candidates ectodomains evidence sequence similarity between archaeal candidates and sexual (HAP2/GCS1) fusexins. HMMs were constructed for each ectodomain sequence (UniProt and NCBI identifiers shown) (*53*). HAP2/GCS1 and EFF-1 sequences were chosen from representative species of the major eukaryotic lineages where these fusexins are present. Flavi-, alpha-, rubi- and bunyaviruses encompass all currently known viral fusexins. All vs all probabilities of homology as determined by HHblits (*53*) were clustered along rows and columns using UPGMA with Hamming distance. Several sequences selected for this analysis have corresponding crystal structures: Yellow fever virus (UniProt: Q89292, PDB: 6IW5(*137*)), Chikungunya virus (UniProt: Q1H8W5, PDB: 3N43(*132*)), Dengue virus (UniProt: P12823, PDB: 1OAN(*138*)), Semliki forest virus (UniProt: P03315, PDB: 1RER(*131*)), Tick-borne encephalitis virus (UniProt: P14336, PDB: 1SVB(*139*)), *Arabidopsis thaliana* (UniProt: F4JP36, PDB: 5OW3(*4*)), Rubella virus (UniProt: P08563, PDB: 4ADG(*23*)), *Chlamydomonas reinhardtii* (UniProt: A4GRC6, PDB: 5MF1 (*4*)), Hantavirus (UniProt: P08668, PDB: 5LK1 (*140*)) and Rift valley fever virus (UniProt: P03518, PDB: 4HJC (*141*)). Although all of the sequences used as input belong to the fusexin structural superfamily, HMM vs HMM comparisons can only detect homology within subsets of the superfamily.

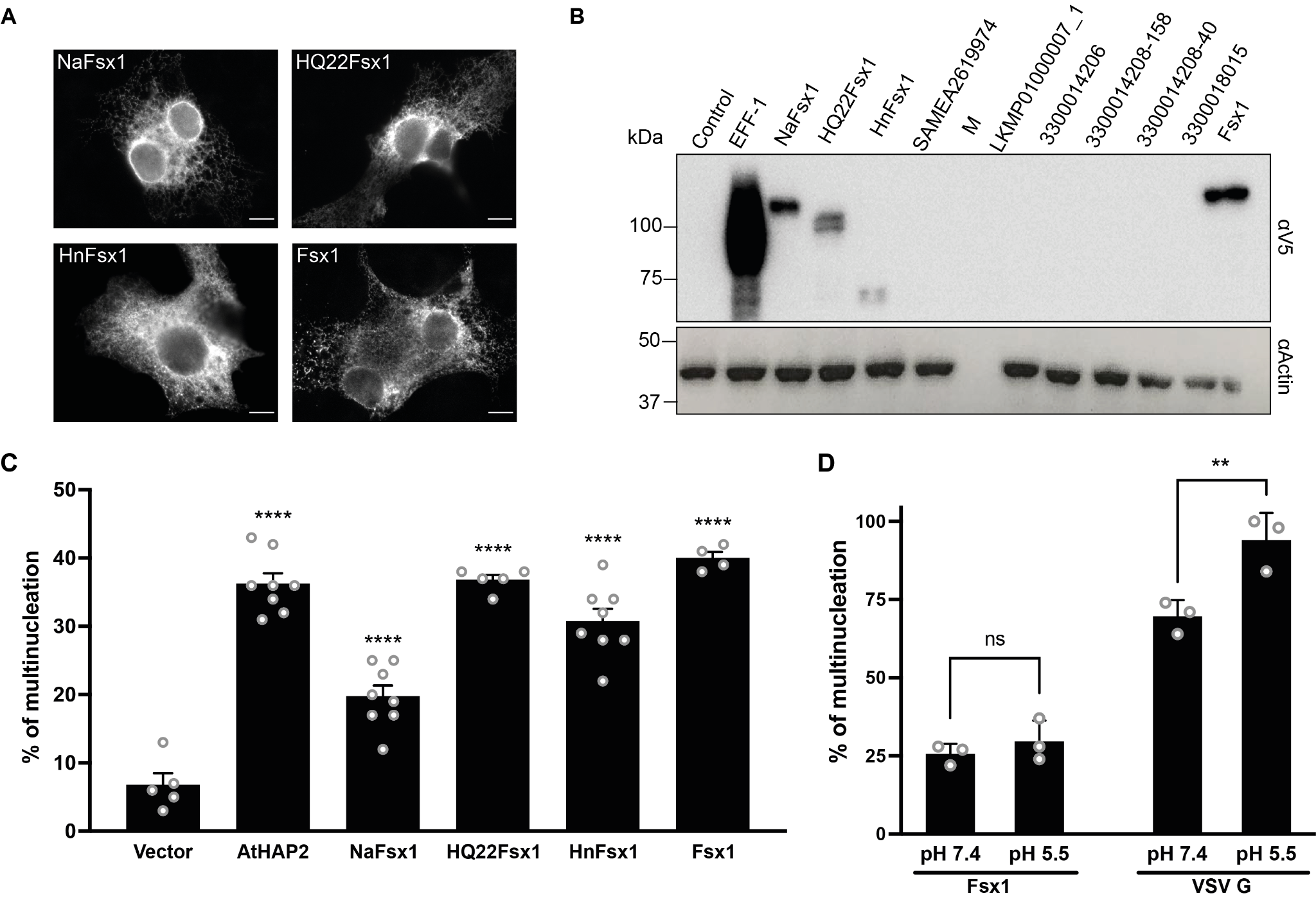

**Fig. S2. Ectopic expression of archaeal fusexins in BHK cells.** (**A** and **B**) Ten archaeal genes were synthesized (table S5) and independently expressed in BHK cells using an inducible promoter. Immunofluorescence (A) and Western blot (B) showing ectopic expression detected with anti-V5 antibody. EFF-1 from *C. elegans* was used as a positive control. NaFsx1, *Natrinema altunense* Fsx1. HQ22Fsx1, *Haloferax* sp. Q22 Fsx1. HnFsx1, *Haloplanus natans* Fsx1. LKMP01000007_1 was obtained from Nanohaloarchaea B1-Br10_U2g21 LB-BRINE-C121. Fsx1 (the protein subsequently characterized), SAMEA2619974 and sequences starting with “330” were obtained from metagenomic databases (see table S5 for complete accession numbers). Scale Bars, 10 µm. (**C**) Quantification of multinucleation in cells expressing archaeal fusexins. Cells were transfected with archaeal fusexins cloned into pCI::H2B-RFP/GFP vectors separately. 48 h post-transfection, immunofluorescence was performed with anti-V5 antibody. Empty vector pCI::H2B-RFP or pCI::H2B-GFP co-transfected with myr-EGFP were the negative controls. AtHAP2 was used as a positive control. Multinucleation was determined as the ratio between the number of nuclei in multinucleated cells and the total number of nuclei in multinucleated cells and expressing cells that were in contact but did not fuse. The percentage of multinucleation is presented as individual data and means ± SEM of independent experiments (n≥4). Total number of nuclei counted in multinucleated cells and in cells in contact n ≥ 1,000 for each experimental condition. Comparisons were made with one-way ANOVA followed by Dunett's test against the empty vector. **** p< 0.0001. **(D)** VSV G activity, but not Fsx1, is enhanced at low pH. Quantification of multinucleation in cells expressing Fsx1 or VSV G. Cells were transfected with pCI::H2B-RFP bearing the coding sequence for Fsx1 or VSV G. 48 h post-transfection, a 5-minute incubation at pH 5.5 buffer was performed to some cells. After 2 h, cells were fixed and immunofluorescence was performed with anti-V5 antibody or anti-G for Fsx1 and VSV G, respectively. Multinucleation was determined as the ratio between the number of nuclei in multinucleated cells and the total number of nuclei in multinucleated cells and expressing cells that were in contact but did not fuse. The percentage of multinucleation is presented as individual data and means ± SEM of three independent experiments. Total number of nuclei counted in multinucleated cells and in cells in contact n ≥ 1,000 for each experimental condition. Comparisons were made with two-way ANOVA. ns, non-significant, ** p < 0.01.

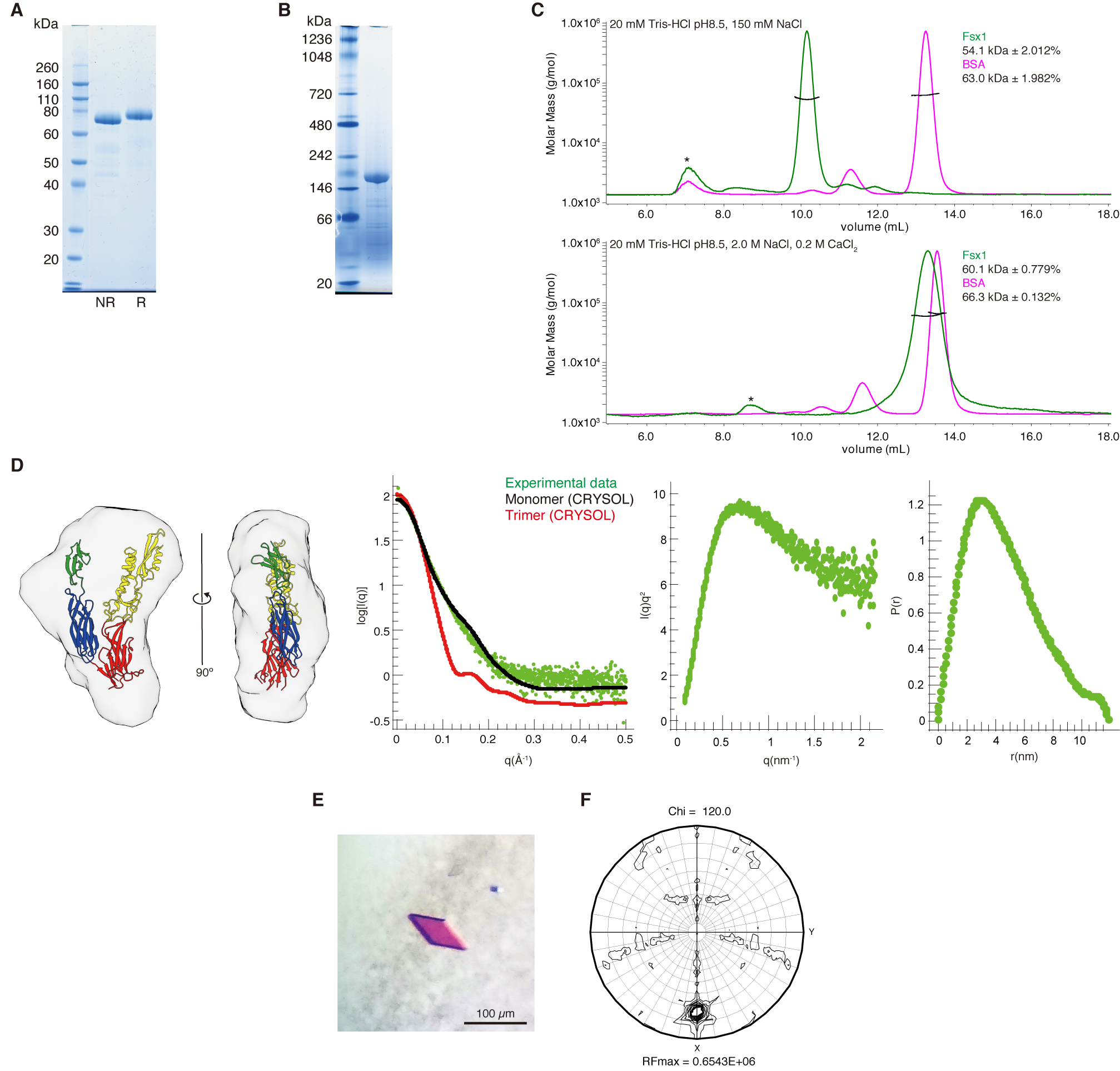

**Fig. S3. The Fsx1 ectodomain is a monomer in solution but crystallizes as a trimer.** (**A** and **B**) SDS-PAGE (A) and blue native PAGE (B) gels of purified Fsx1 ectodomain (Fsx1_E_). NR; non-reducing conditions. R; reducing conditions (see fig. S18). (**C**) Size exclusion chromatography-multiangle light scattering (SEC-MALS) shows that, although Fsx1_E_ has a very different elution volume depending on the salt concentration, it is a monomer in solution in both normal and high salt conditions. BSA, whose elution volume does not change significantly at different salt concentrations, is used as a control. Asterisks indicate the high-molecular weight aggregate. (**D**) SAXS analysis of Fsx1_E_. Left panel, The SAXS envelope of Fsx1_E_, obtained by averaging of *ab initio* shape reconstructions, is consistent with the crystallographic model of Fsx1_E_ chain A. Centre-left panel, Comparison of the experimental SAXS profile of Fsx1_E_ (green dots) and theoretical scattering curves calculated from the refined coordinates of Fsx1_E_ chain A (black dots) or the whole Fsx1_E_ trimer (red dots). Center-right panel, the Kratky plot of Fsx1_E_ suggests the presence of significant flexibility between the domains of the monomeric protein. Right panel, Pairwise interatomic distance distribution of Fsx1_E_. (**E**) Representative rhomboidal plate crystal of Fsx1_E_. (**F**) The Chi=120 section of the self-rotation function of Fsx1_E_ (calculated using a 67.3-2.6 Å resolution range) shows a prominent peak with a height of 72% of the origin peak.

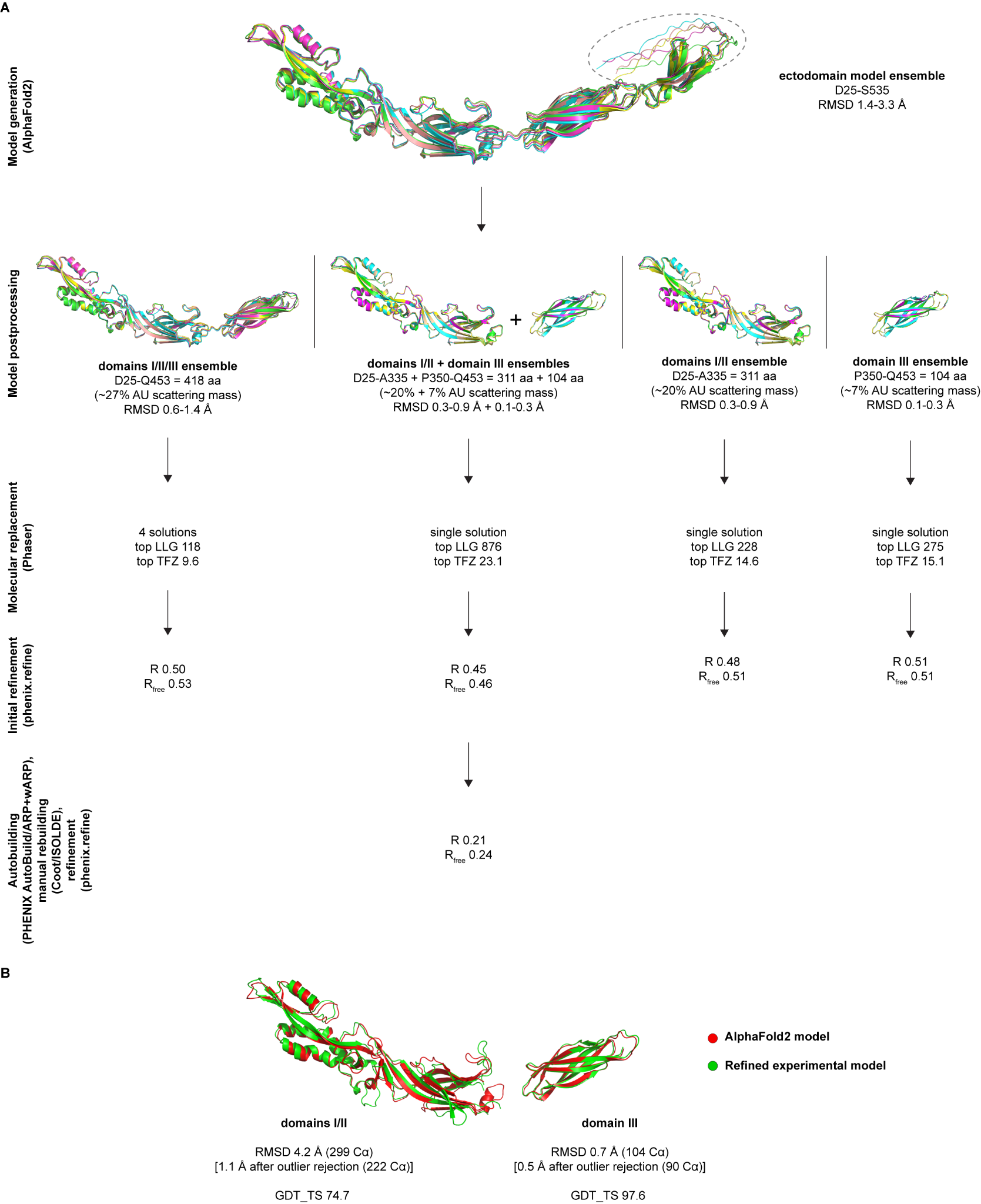

**Fig. S4. AlphaFold2-aided MR phasing of Fsx1_E_.** (**A**) Flowchart of Fsx1_E_ structure determination using different AlphaFold2 model fragments or a combination thereof. The dashed oval in the top panel indicates C-terminal residues D510-S535, predicted with low-confidence by AlphaFold2. aa, amino acid; AU, asymmetric unit. (**B**) Comparison of the AlphaFold2 and final crystallographic models of Fsx1 domains I/II and III.

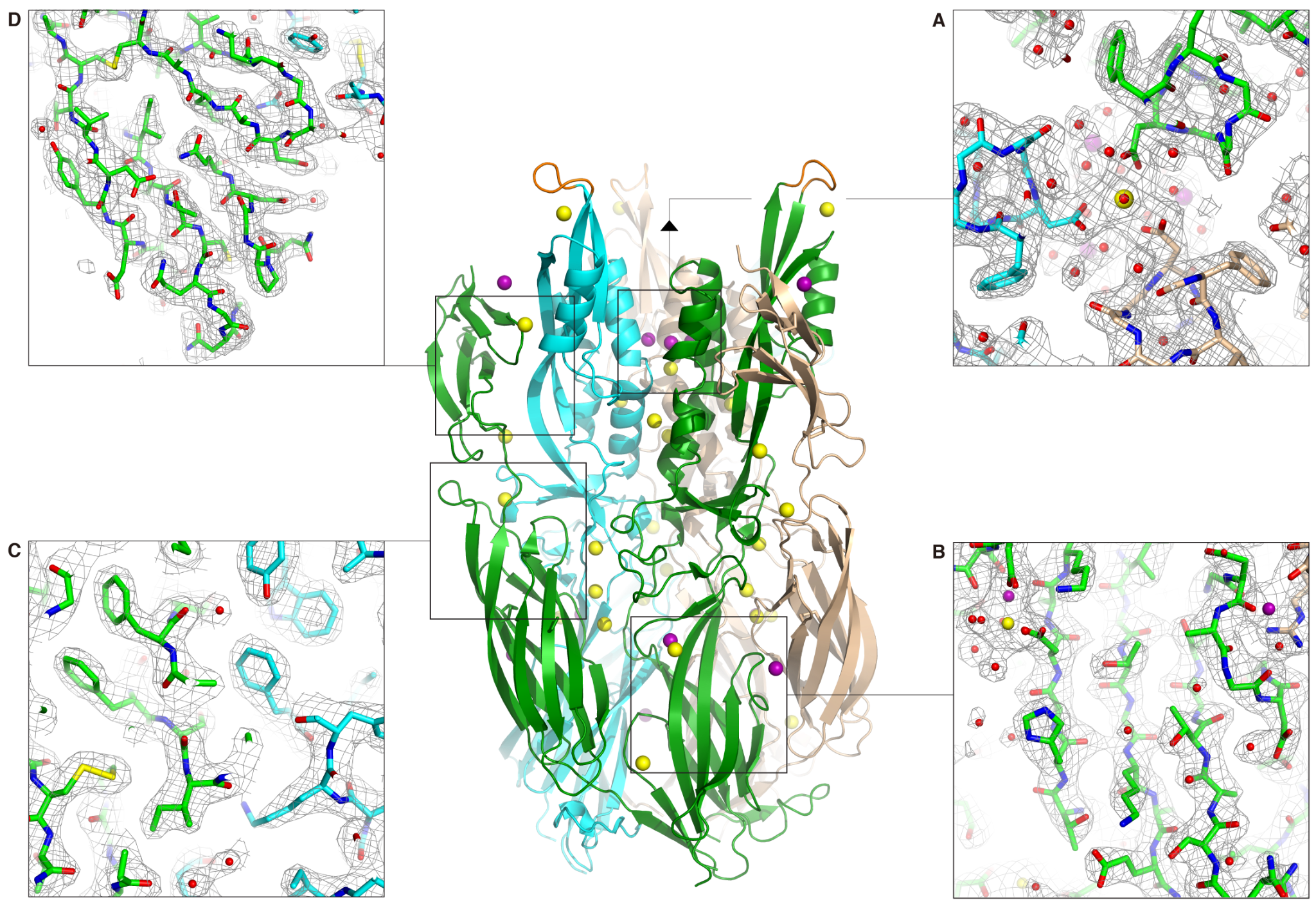

**Fig. S5. Details of the electron density map of Fsx1_E_.** (**A**) View of the domain II helical bundle, looking down the molecular three-fold axis from the center of the structure towards the putative fusion loop end. Domain II α1 helix residues D260 and D263 coordinate a Ca^2+^ ion sitting on the NCS axis and three symmetrically positioned Na^+^ ions, respectively. The refined *2mFo-DFc* electron density map, contoured at 1.0 σ, is shown as a gray mesh superimposed onto the protein model in stick representation. Fsx1 subunits and metal ions are coloured as in Fig. 2C. (**B**) Section of the map centered around β-strands D_0_, E_0_ and F_0_ of domain I. (**C**) Closeup of the map region where domain III of one protein subunit interacts with domains I and II of another. Clear density for the C_3_389-C_4_432 disulfide is visible near the bottom left corner. (**D**) Map of domain IV. The conserved C_7_490-C_8_506 disulfide is at the top left corner, whereas C_6_477 of the C_5_457-C_6_477 disulfide is visible at the bottom and the domain II C_1_125-C_2_155 disulfide of the adjacent subunit can be seen on the top right corner.

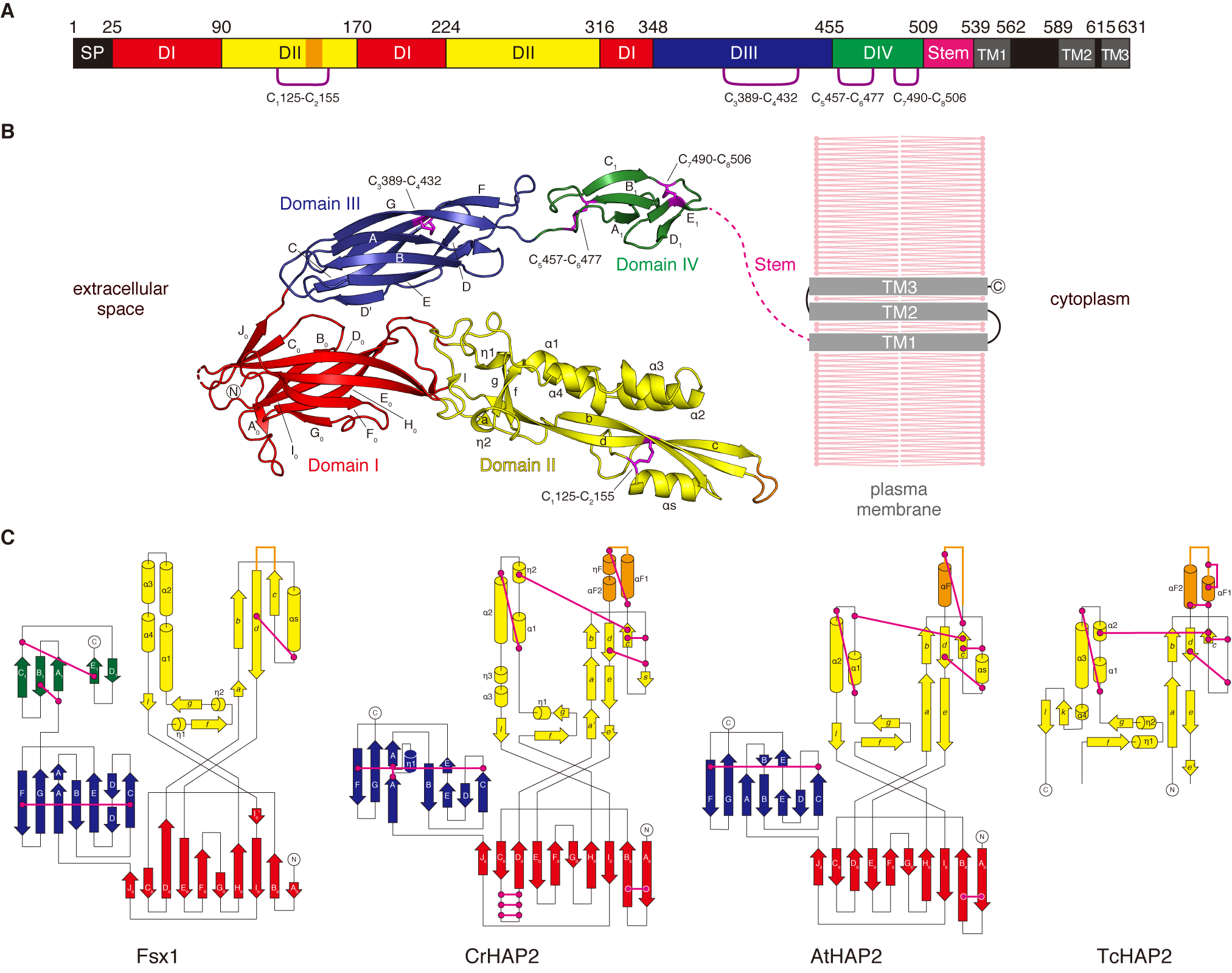

**Fig. S6. Domain architecture of Fsx1 and topological comparison with HAP2.** (**A**) Schematic diagram of the domains of Fsx1. SP, signal peptide; TM, predicted transmembrane helices. Note that Fsx1 is predicted to contain three C-terminal TMs, with a cytoplasmic loop between TM1 and TM2 that lacks Cys residues; on the other hand, HAP2 homologs are characterized by having a single TM, followed by a cytoplasmic tail that often contains Cys implicated in fusion (*142*). (**B**) Crystal structure of the ectodomain of Fsx1 and predicted topology of the full-length protein relative to the plasma membrane. Domains I, II, III and IV are shown in red, yellow, blue and green, respectively; disulfide bonds are indicated and coloured magenta. (**C**) Topology diagrams of the ectodomains of Fsx1, *C. reinhardtii* HAP2 (PDB 6E18 (*51*)), *A. thaliana* HAP2 (PDB 5OW3 (*17*)) and *T. cruzi* HAP2 (PDB 5OW4 (*17*)). Domains and disulfide bonds are coloured as in panel (B). Note how, although domain II of Fsx1 has the same topology as the corresponding domain of HAP2, it contains only one of its conserved disulfide bonds (C_1_125-C_2_155, corresponding to disulfide bond 3 of CrHAP2 (*4*) and disulfide bond 5 of CeEFF-1 (*10*)).

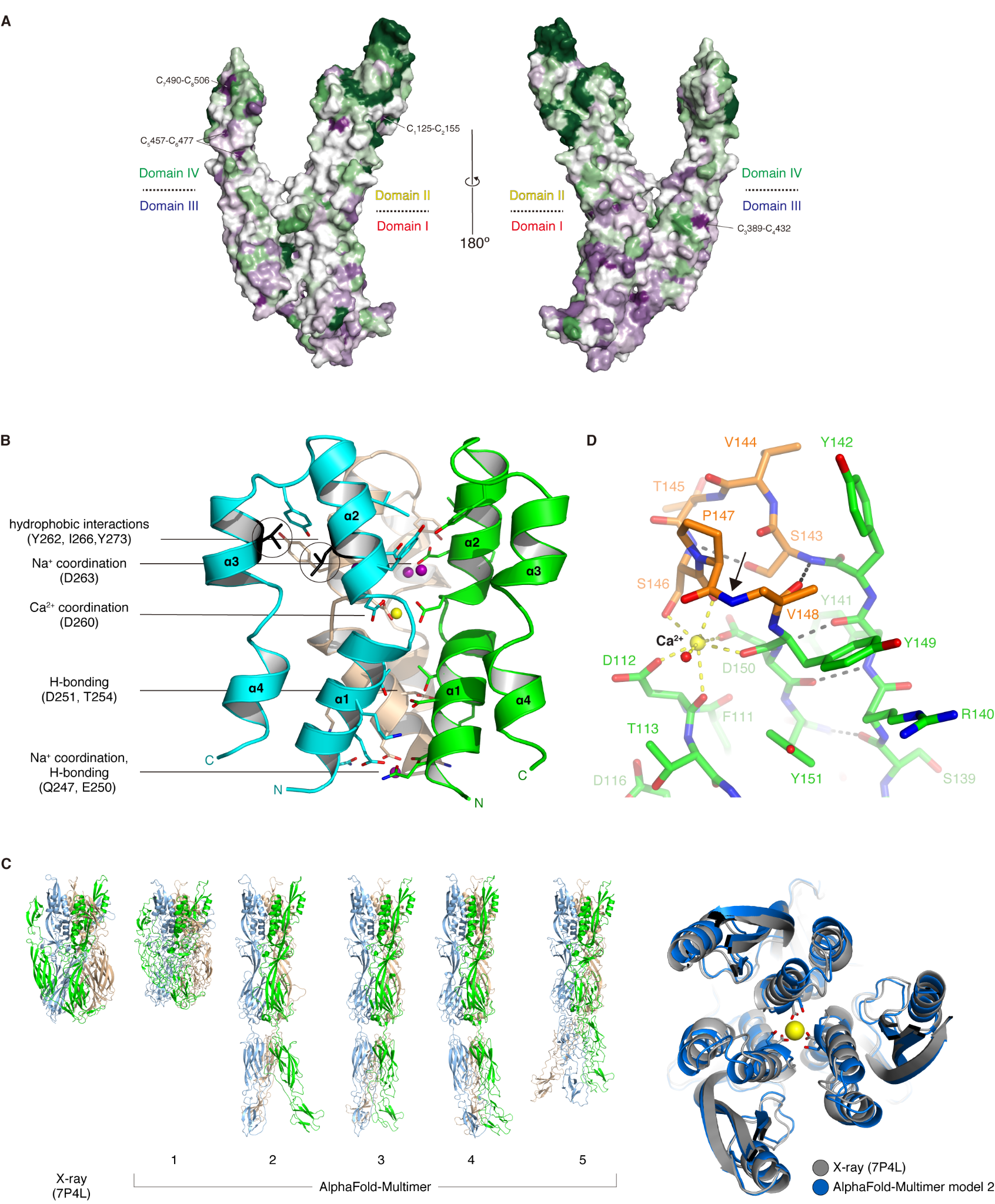

**Fig. S7. 3D mapping of the evolutionary conservation of Fsx1 residues and key structural features of its domain II.** (**A**) Fsx1 domains I and III are more evolutionarily conserved than domains II and IV. Surface representation of the Fsx1_E_ monomer, with residues coloured from green to violet by increasing conservation among archaeal homologs. Approximate domain boundaries are marked, and the position of the four highly conserved disulfides of Fsx1 is indicated. (**B**) Helices α1 and α2 of domain II form a six-helix bundle around the molecular three-fold axis. Subunits are shown in cartoon representation and coloured as in Fig. 2C, with residues mediating direct or ion-mediated interactions between chains depicted in stick representation (for clarity, water-mediated interactions are not shown). The side chains of residues α2 L264 and α3 V278 (black circles), which face each other in the Fsx1_E_ structure, are both replaced by Cys in the sequences of *Halogeometricum borinquense* and *Halobonum sp.* (fig. S8), suggesting that an additional disulfide bond stabilizes the helical bundle of the Fsx1 homologs from these species. (**C**) AlphaFold-Multimer (*101*) prediction of the Fsx1_E_ homotrimer (left panel) generates models that either approximate the overall post-fusion conformation of the crystal structure (albeit with significant interchain clashes; model 1) or adopt an extended conformation that resembles an intermediate state thought to exist before fusion (*7*) (models 2-5). In both cases, the models contain the experimentally observed six-helix bundle and, as shown for the Ca^2+^ coordinated by D260 (yellow sphere), in some instances even reproduce the orientation of side chains that bind ions in the crystal (right panel). Considering that these residues are poorly conserved in other homologs of Fsx1 (fig. S8) and taking into account that AlphaFold2 does not explicitly predict ions (*19*), this suggests that Ca^2+^ and Na^+^ stabilize the trimeric structure of Fsx1 rather than being required for its formation. (**D**) Structure of the fusion loop of Fsx1_E_. The domain II region encompassing the loop is shown in the same orientation as in Fig. 2F, with black and yellow dashes indicating protein hydrogen bonds and the coordination of the Ca^2+^ ion, respectively. Note how binding of the ion locally twists the protein main chain, with the peptide bond between P147 and V148 adopting a *cis* configuration (black arrow).

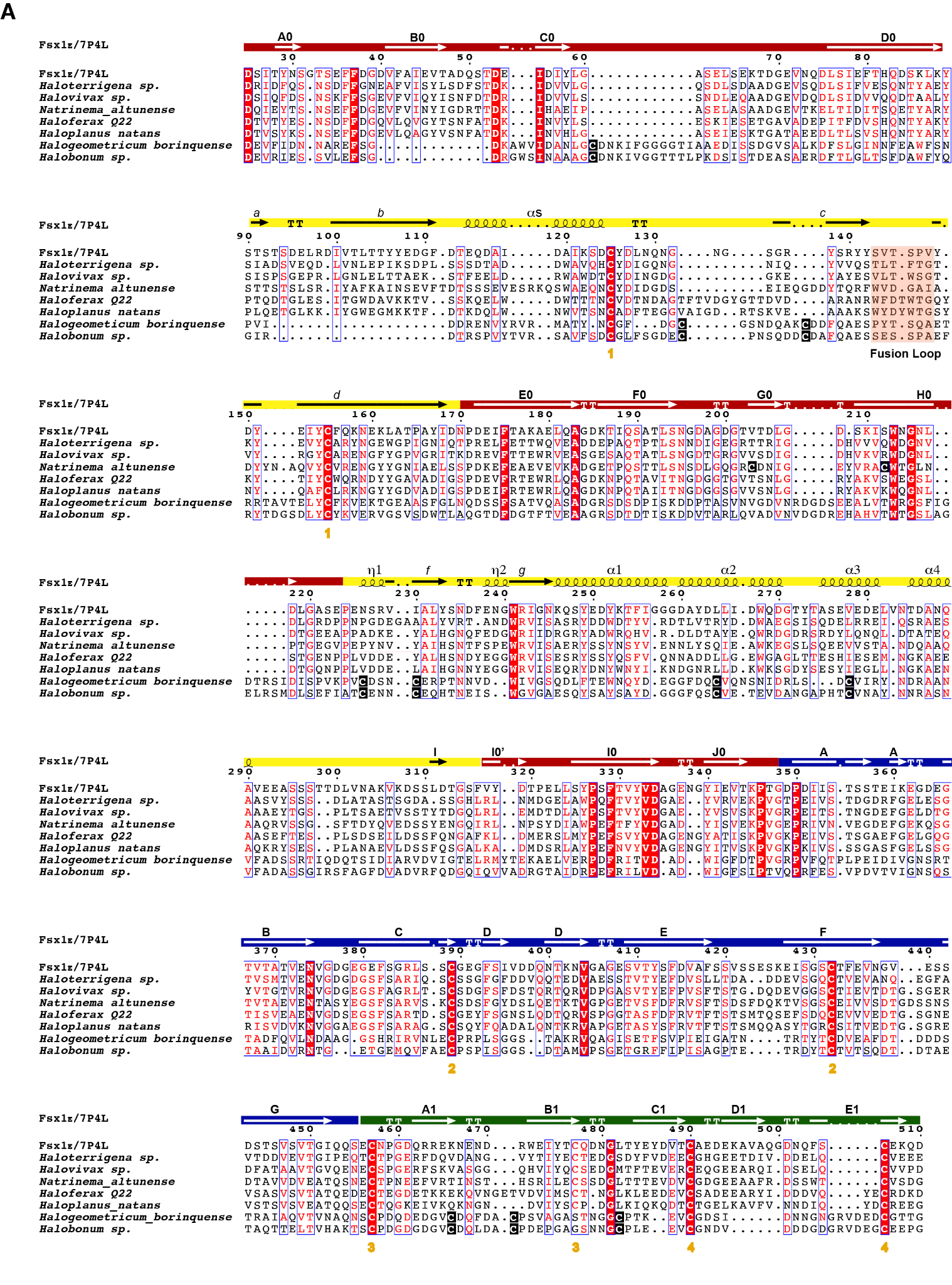

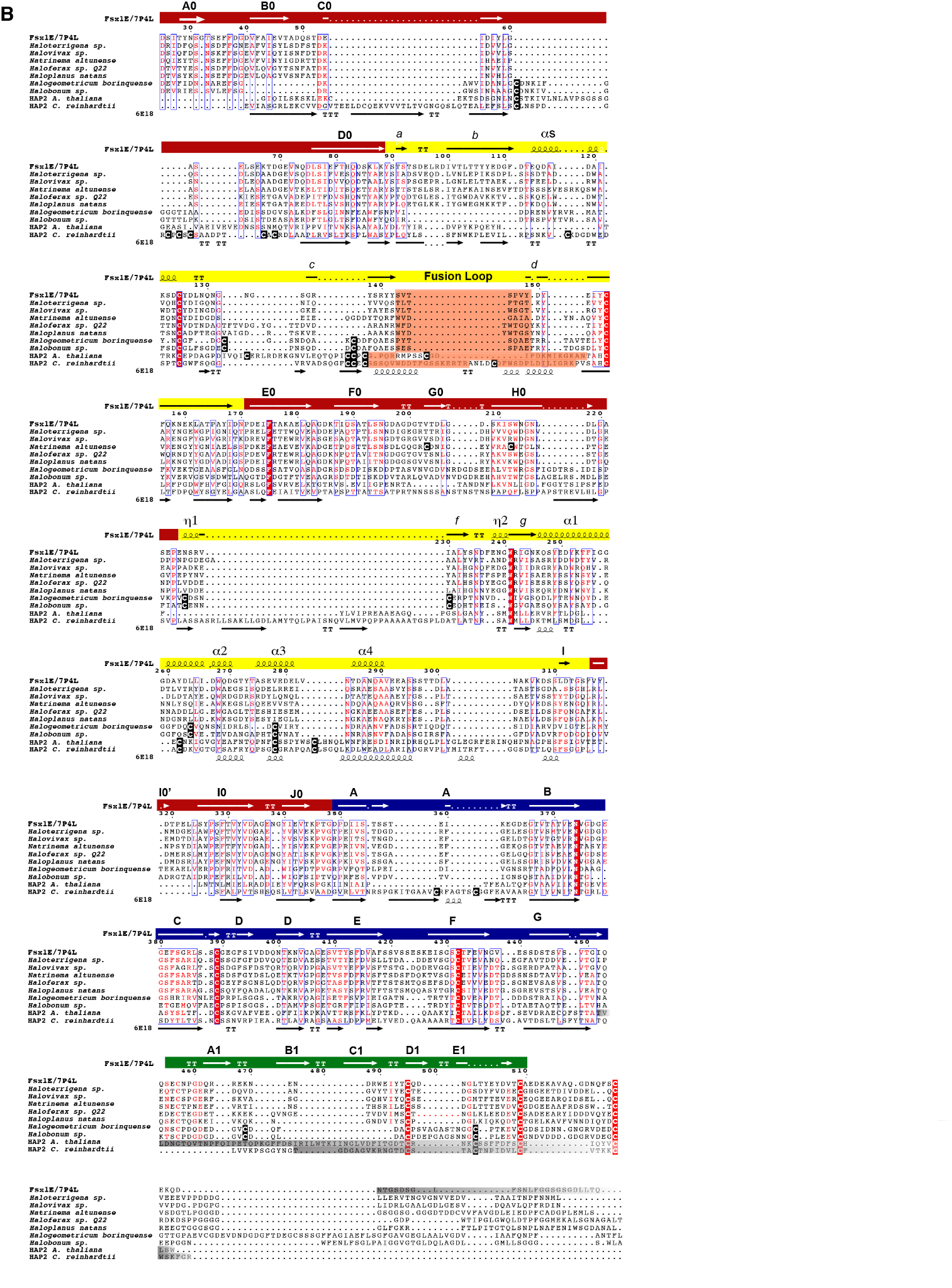

**Fig. S8. Sequence alignment of archaeal and HAP2 fusexin ectodomains.** For reproducibility, no gap or block was altered from alignments. Identical column residues are depicted in bold white on a red background; conserved positions are boxed and labeled red. On top are displayed secondary structure elements and sequence numbering corresponding to Fsx1_E_/7P4L. Made with ESPript (*143*) and manually annotated. (**A**) Archaeal sequences from PCGs, with N- and C-terminal regions cropped to match Fsx1_E_ (PDB: 7P4L). Secondary structure elements are shown within boxed domains, coloured and labeled following the previous nomenclature (domain I, red; domain II, yellow; domain III, blue). Disulfide bonds are indicated by orange numbers below the alignment. Additional, lineage-specific cysteines are black-boxed and depicted in bold white. The fusion (cd) loop is highlighted in light orange within the alignment; as in eukaryotic HAP2, it has poor sequence conservation (see also fig. S7A) but shows a high prevalence of hydrophobic residues (see also fig. S17). Domain IV (green) has relatively poor sequence conservation within archaea (fig. S7A), yet preserves its disulfides. (**B**) Sequence alignment of HAP2s from *A. thaliana* and *C. reinhardtii* (6E18) and archaeal fusexins from cultivated genomes. Of note are the inserts of HAP2 relative to Fsx1s, as well as the absence of domain IV, which seems confined to Archaea. The conserved fusion loop region is orange shaded; lineage-specific cysteines black boxed. C-terminal regions shaded with a gradient from gray to white indicate portions absent from Fsx1_E_ (7P4L) and HAP2 (5OW3 and 6E18) crystal structures, either with no electron density or absent from the expression construct. HAP2s introduce gaps; for clarity, disulfides shown in (A) are omitted and replaced by secondary structure elements from 6E18.

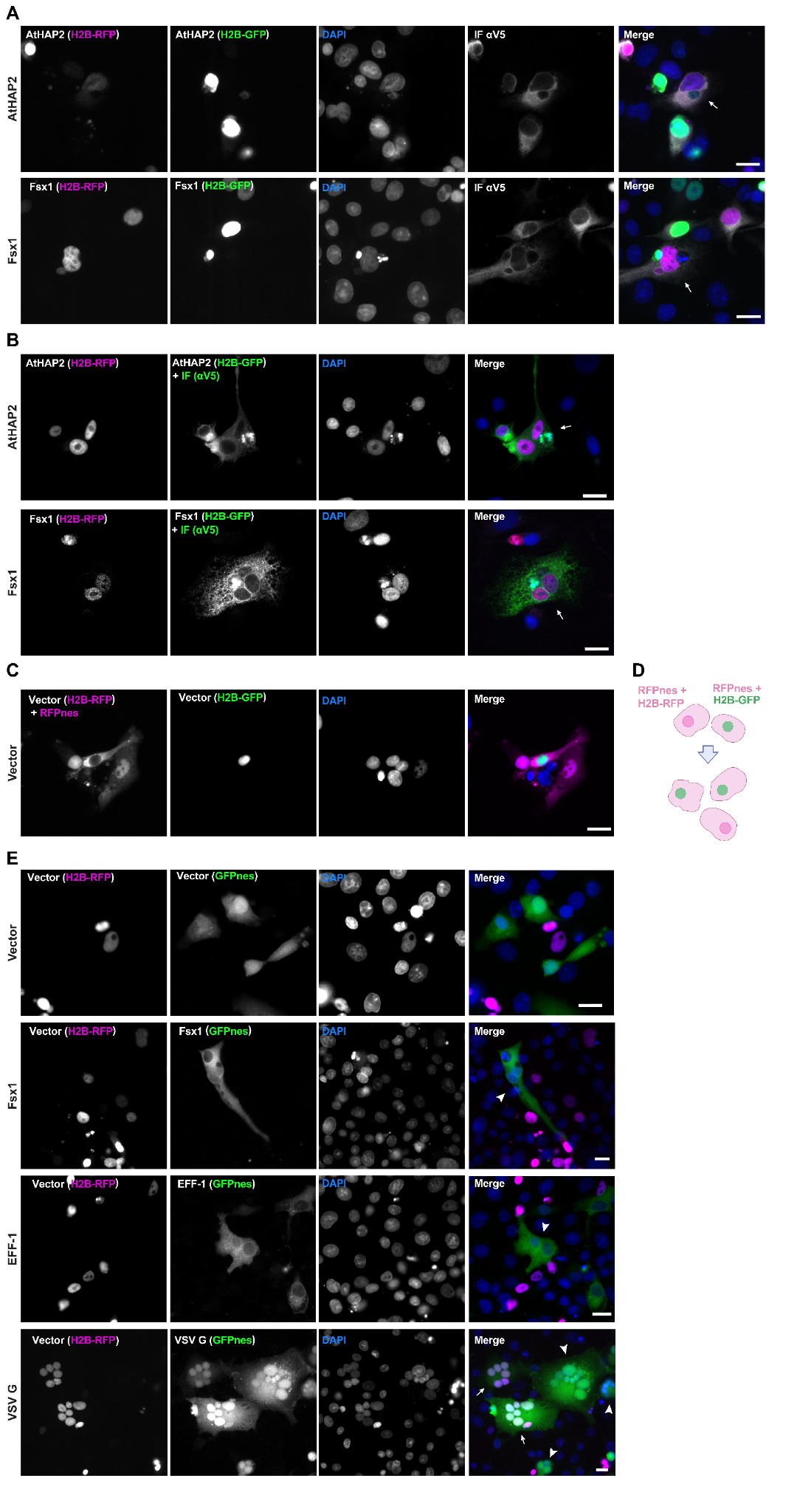

**Fig. S9. Fsx1 mediates bilateral cell-cell fusion.** (**A**) Images from Fig. 3A in each separate channel (red, green, DAPI and far red) and merge. (**B**) Multinucleated cells containing green nuclei (H2B-GFP) and magenta nuclei (H2B-RFP) (arrows). Immunofluorescence against the V5 tag was performed in green to facilitate counting. (**C** and **D**) In the negative control, cell-cell fusion was measured by content-mixing, indicated by the appearance of multinucleated cells containing green nuclei (H2B-GFP) and magenta nuclei (H2B-RFP). To reveal the cytoplasm of the transfected cells, a plasmid encoding for cytoplasmic RFP (RFPnes) was co-transfected. C, Representative images of mononucleated cells with a green or red nucleus. DAPI staining is shown in blue. D, Cartoon showing the experimental design for negative control. (**E**) Images from Fig. 3D in each separate channel (red, green and DAPI) and merge. Multinucleated GFPnes only (arrowheads) or mixed cells (arrows). Scale bars, 20 µm.

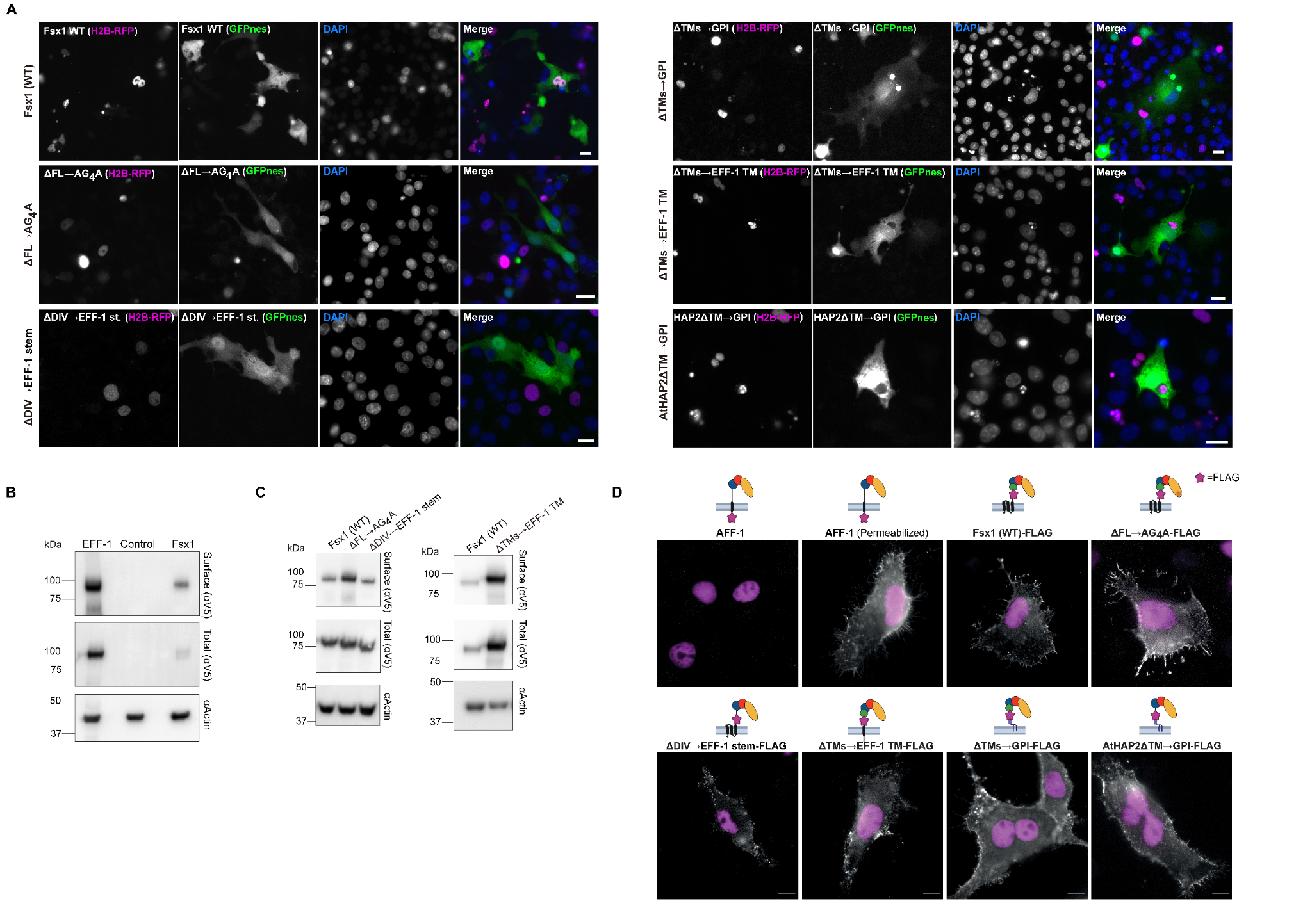

**Fig. S10. Structure-function analysis of Fsx1.** (**A**) Images from Fig. 3J in each separate channel: red (RFP); green (GFP) and blue (DAPI) and the merged images. Scale bars, 20 µm. (**B**) Immunoblot of EFF-1-V5, control (untransfected cells) and Fsx1-V5 expressing cells. “Surface” indicates surface biotinylation followed by affinity purification using neutravidin agarose beads; “Total” indicates the expression in whole cell extracts. Actin is used as a loading control. The amount of initial cells for Fsx1 is 4 times higher than EFF-1 (fig. S19B). (**C**) Surface biotinylation as explained in “B” for cells expressing Fsx1-V5 (WT), ΔFL→AG_4_A-V5, ΔDIV→EFF-1 stem-V5 or ΔTMs→EFF-1 TM-V5 (Fig. 3I and fig. S19C). (**D**) Immunofluorescence images on non-permeabilized cells expressing Fsx1-FLAG (WT), AFF-1-FLAG (negative control, cytotail), Fsx1-ΔFL→AG_4_A-FLAG, Fsx1-ΔDIV→EFF-1 stem-FLAG, AFF-1-FLAG (permeabilized), Fsx1-ΔTMs→EFF-1 TM-FLAG, Fsx1-ΔTMs→GPI and AtHAP2-ΔTM→GPI. The FLAG tag was inserted before the first TM or the GPI signal of each construct except for *C. elegans* AFF-1 in which the FLAG is at C-terminal after the cytoplasmic tail. Transfected BHK cells were incubated with anti-FLAG antibody on ice before fixation. Non-permeabilized staining of FLAG antibody showed the surface expression of Fsx1 and the mutants. *C. elegans* AFF-1 tagged with a cytoplasmic FLAG is a negative control for non-permeabilized staining. Permeabilized staining of CeAFF-1-FLAG shows the localization on plasma membrane and internal compartments (see also fig. S11). Scale bars, 10 µm.

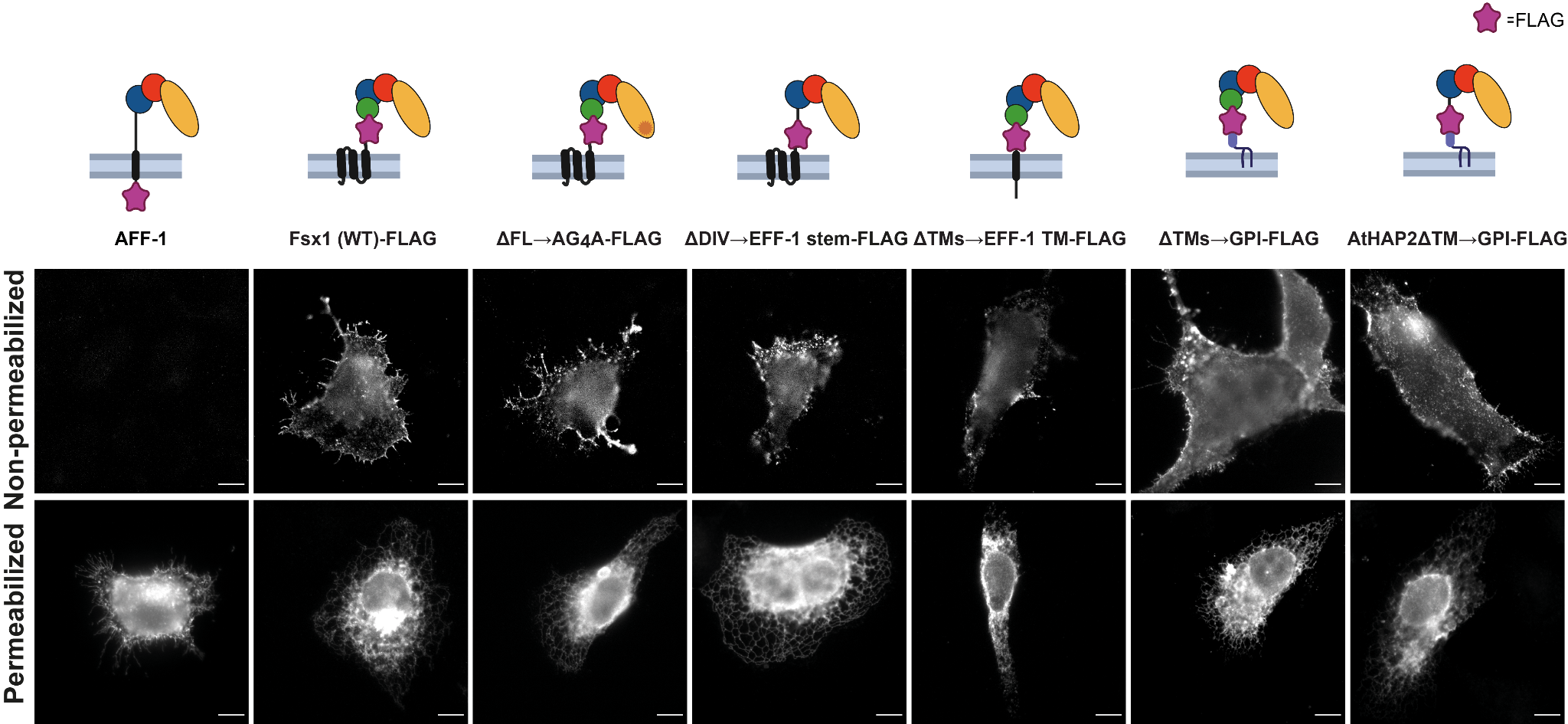

**Fig. S11. Surface expression of Fsx1, mutants and eukaryotic fusexins.** BHK cells were transfected with FLAG-tagged Fsx1 (WT) and the indicated mutants; the FLAG tag was inserted before the membrane anchor (see fig. S10D). Non-permeabilized staining using anti-FLAG antibody showed surface expression of Fsx1 and the various mutants. The proportion of non-permeabilized cells showing surface expression was: AFF-1-FLAG (negative control; 0%, n~1000), Fsx1-FLAG (3.9%, n=1242), Fsx1-ΔFL→AG_4_A-FLAG (4.4%, n=1176), Fsx1-ΔDIV→EFF-1 stem-FLAG (2.6%, n=1263), Fsx1ΔTMs→EFF-1 TM-FLAG (3.3%, n=853), Fsx1ΔTMs→GPI-FLAG (32.6%, n=1045), AtHAP2ΔTM→GPI-FLAG (22.8%，n=997). Another group of transfected BHK cells in parallel were fixed, permeabilized and stained with anti-FLAG antibody. Permeabilized staining showed the main distribution in the cytoplasm (endoplasmic reticulum) of Fsx1 WT, Fsx1 mutants and AtHAP2ΔTM→GPI mutant. *C. elegans* AFF-1 tagged with FLAG at the C terminus (cytoplasmic tail) worked as a negative control for non-permeabilized staining. Scale Bars, 10 µm.

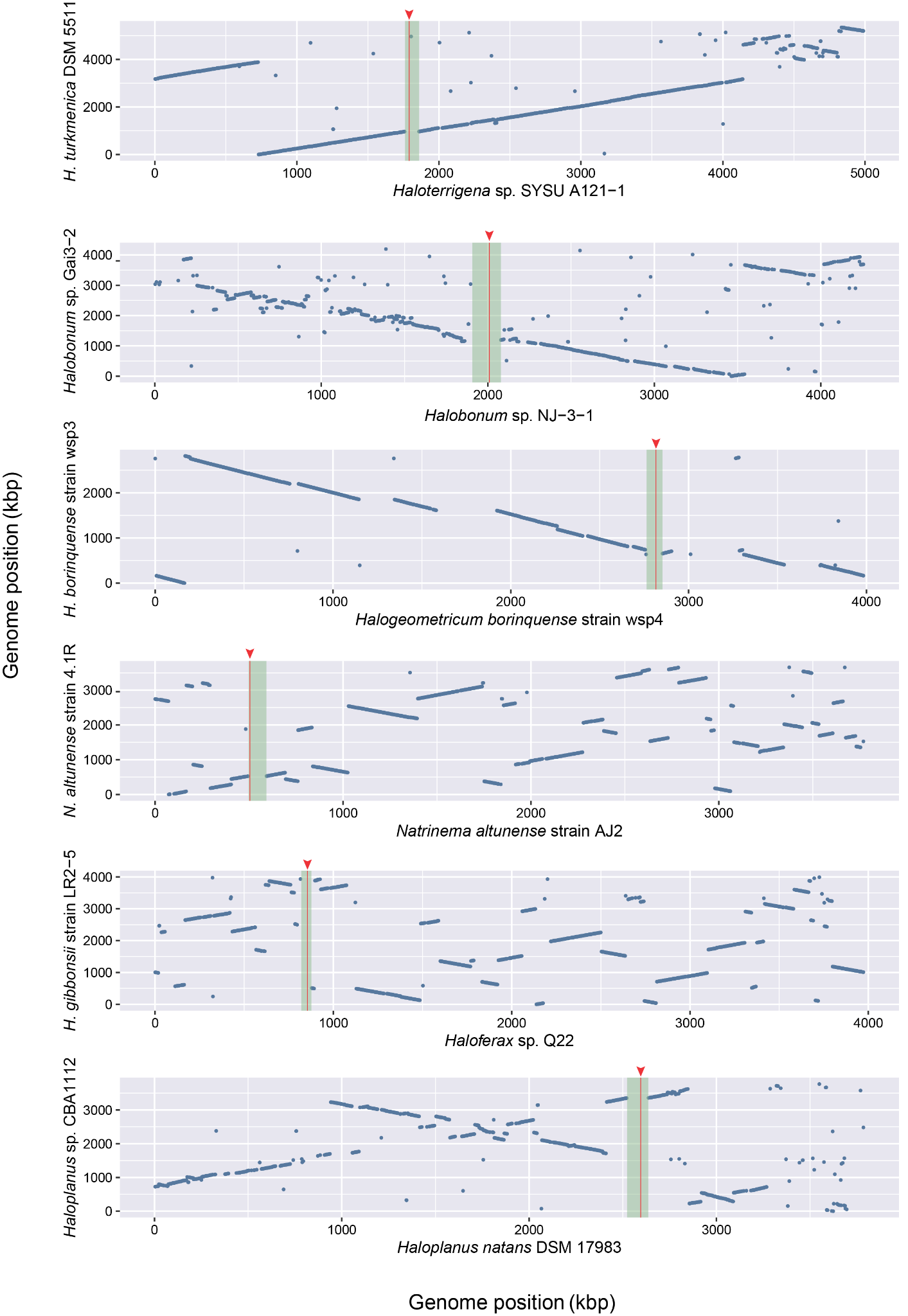

**Fig. S12. Whole genome comparison of species with and without *fsx1*.** Each blue dot represents a segment of 500 bp with more than 80% identity between the species harbouring *fsx1* (e.g. *Haloplanus natans* DSM 17083) and the species with no *fsx1* (e.g. *Haloplanus* sp. CBA1112). Species with *fsx1* are in the x axis, the base of the green rectangles represent the detected IME carrying the *fsx1* gene, locus of *fsx1* is in red vertical line and pointed with a red arrowhead.

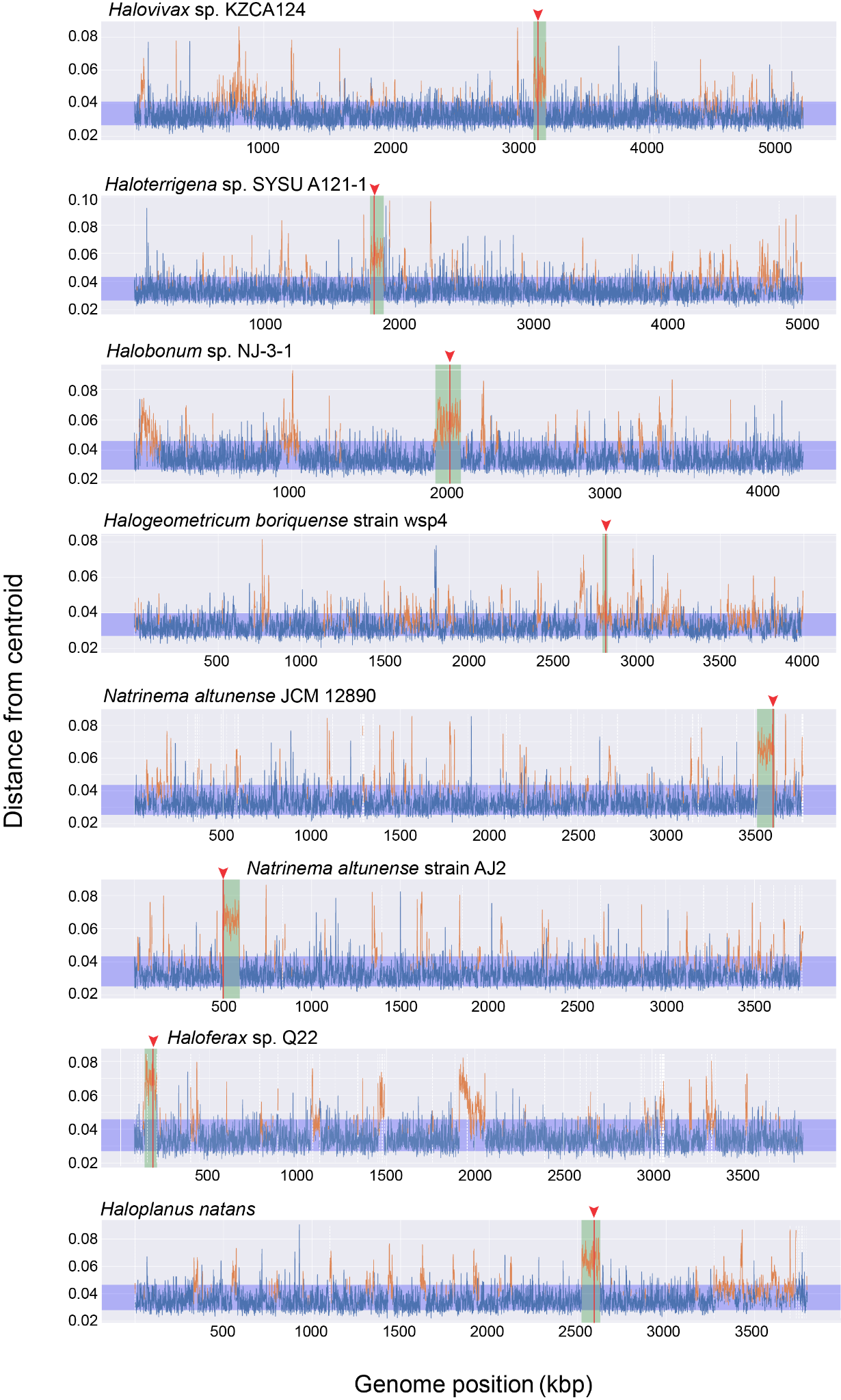

**Fig. S13. K-mer spectra deviation of *fsx1*-containing IMEs.** K-mer spectrum deviation from centroid is shown for each of the PCGs where *fsx1* was detected . Blue region shows the standard deviation. Locus of *fsx1* is in red vertical line and pointed with a red arrowhead, the mobile element containing *fsx1* is in green. Dashed vertical white lines indicate the end of a contig in the genome assembly. *fsx1* is consistently found within regions that deviate from the core genome’s spectrum, indicating they belong to a mobile element.

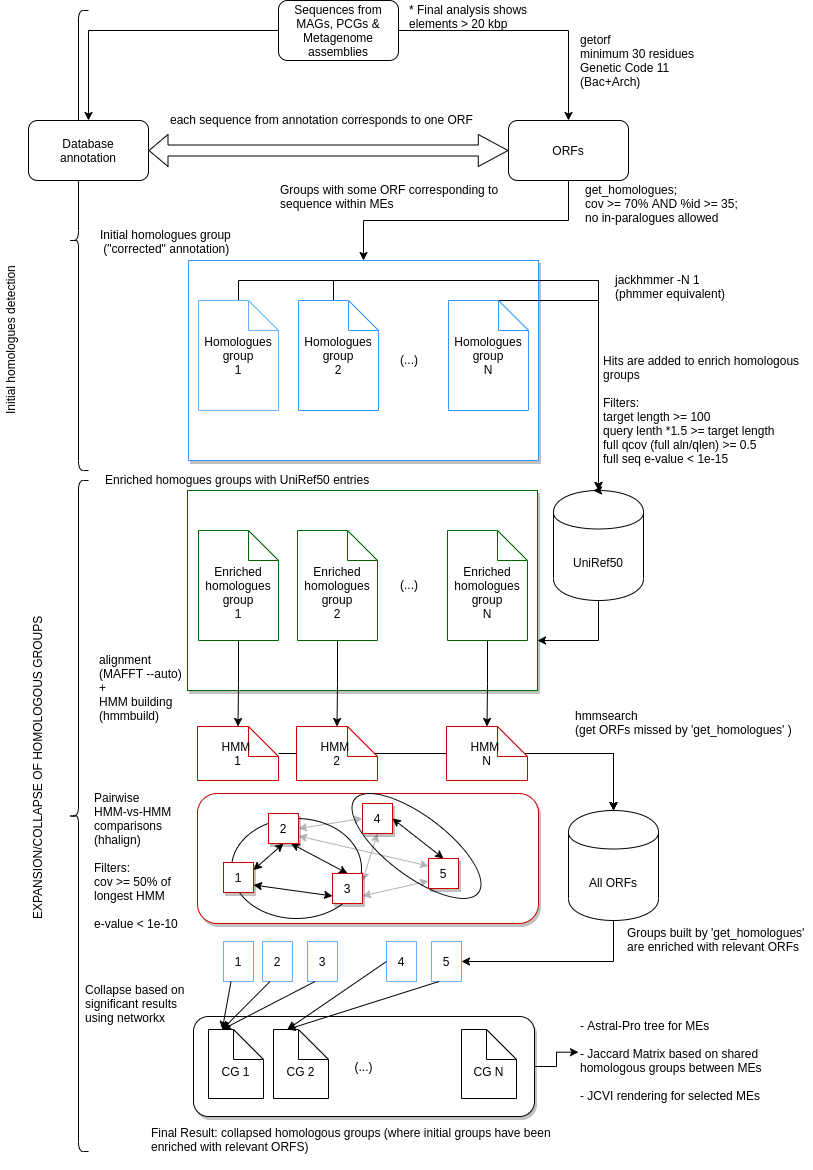

**Fig. S14. Bioinformatics workflow for IMEs.** The general workflow is divided into two parts. First, we re-annotated the ORFs of each IME and searched for potential homologs between them. Then we enriched these initial groups by searching the UNIREF50 database and generated new HMM profiles. With these new HMMs we searched again within each IME to capture any potentially missing homolog. Finally, we performed HMM vs HMM(*58*) search to collapse similar groups into one. CG1, CG2...CGN are final collapsed homologous groups which are the basis for IME clustering, synteny conservation and gene content analyses shown in figs. S15 and S16, and table S3 and data S3, respectively.

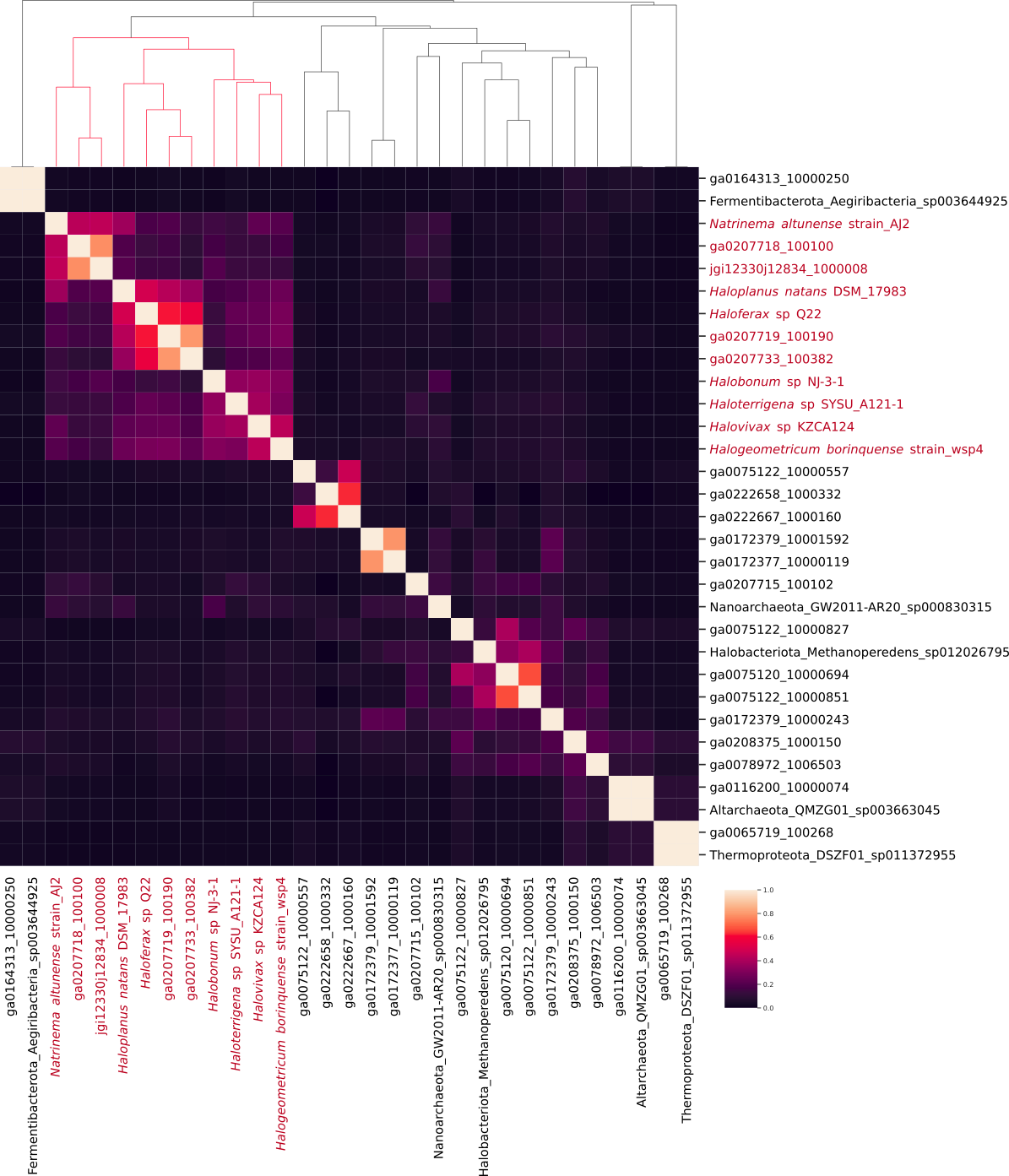

**Fig. S15. Clustering of IMEs from complete genomes, MAGs and metagenomic contigs based on gene content**. A distance metric based on the sharing of homologous genes between all IMEs was computed (see Methods). Then a pairwise distance matrix was built to perform hierarchical clustering. There is a clear cluster, marked in red, which contains all PCG IMEs and the DNA contig containing the crystallized Fsx1 (jgi12330j12834_1000008). This cluster of 11 IMEs was used for the synteny conservation analysis.

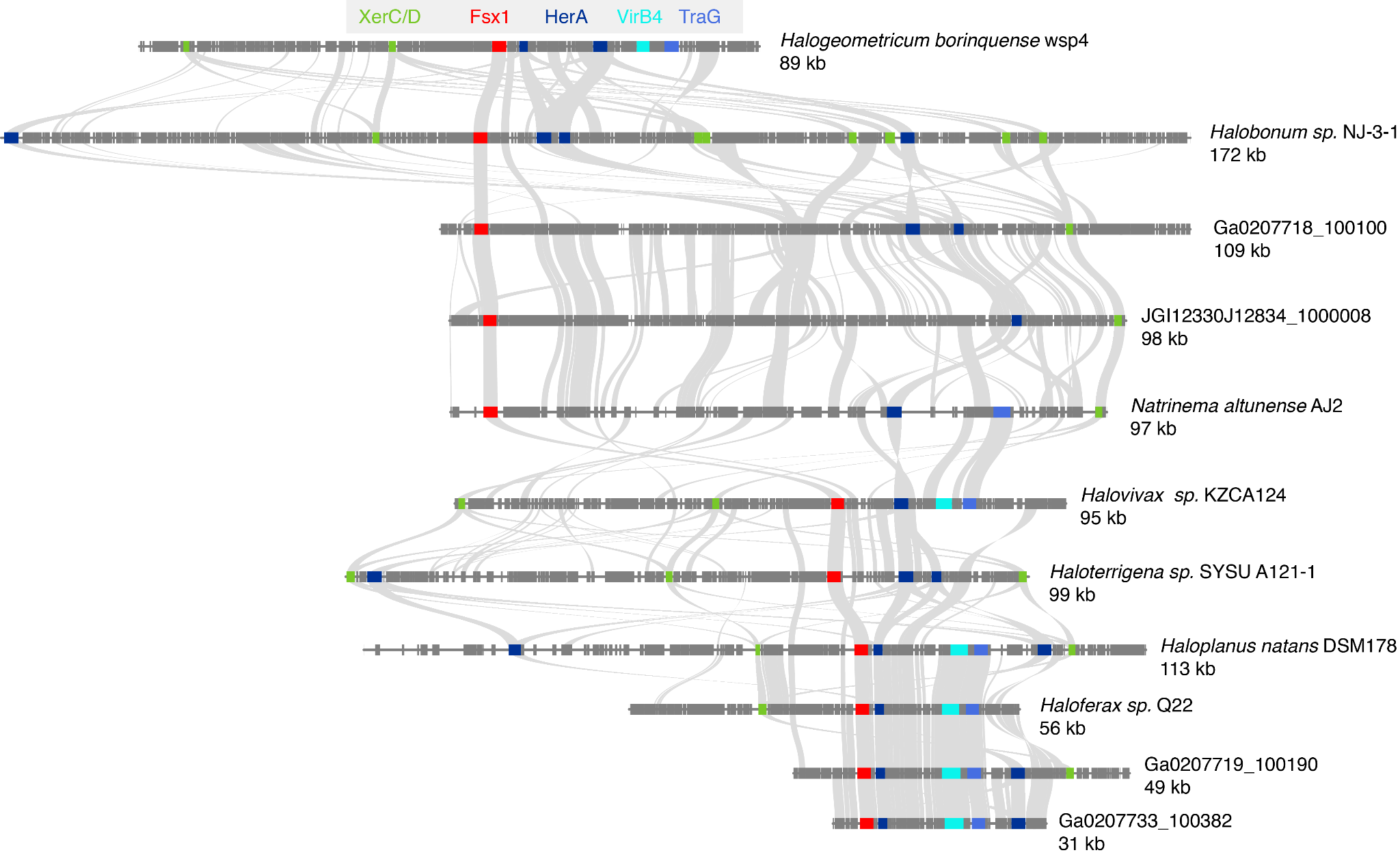

**Fig. S16. Synteny plots for IMEs from complete genomes and metagenomic data.** Annotated regions plus inferred ORFs belonging to homologous clusters identified by the workflow depicted in fig. S14. Homology relationships are represented by gray links. *fsx1* genes are marked in red and selected ORFs homologous to IME signature genes are labeled and color-coded. XerC/XerD recombinases (green); HerA helicase (dark blue); VirB4, Type IV secretory (T4SS) pathway (cyan); TraG/TraD/VirD4 family enzyme, ATPase, T4SS (see table S3 and data S3 for details). The eleven segments analyzed correspond to the cluster marked in red in fig. S15**.**

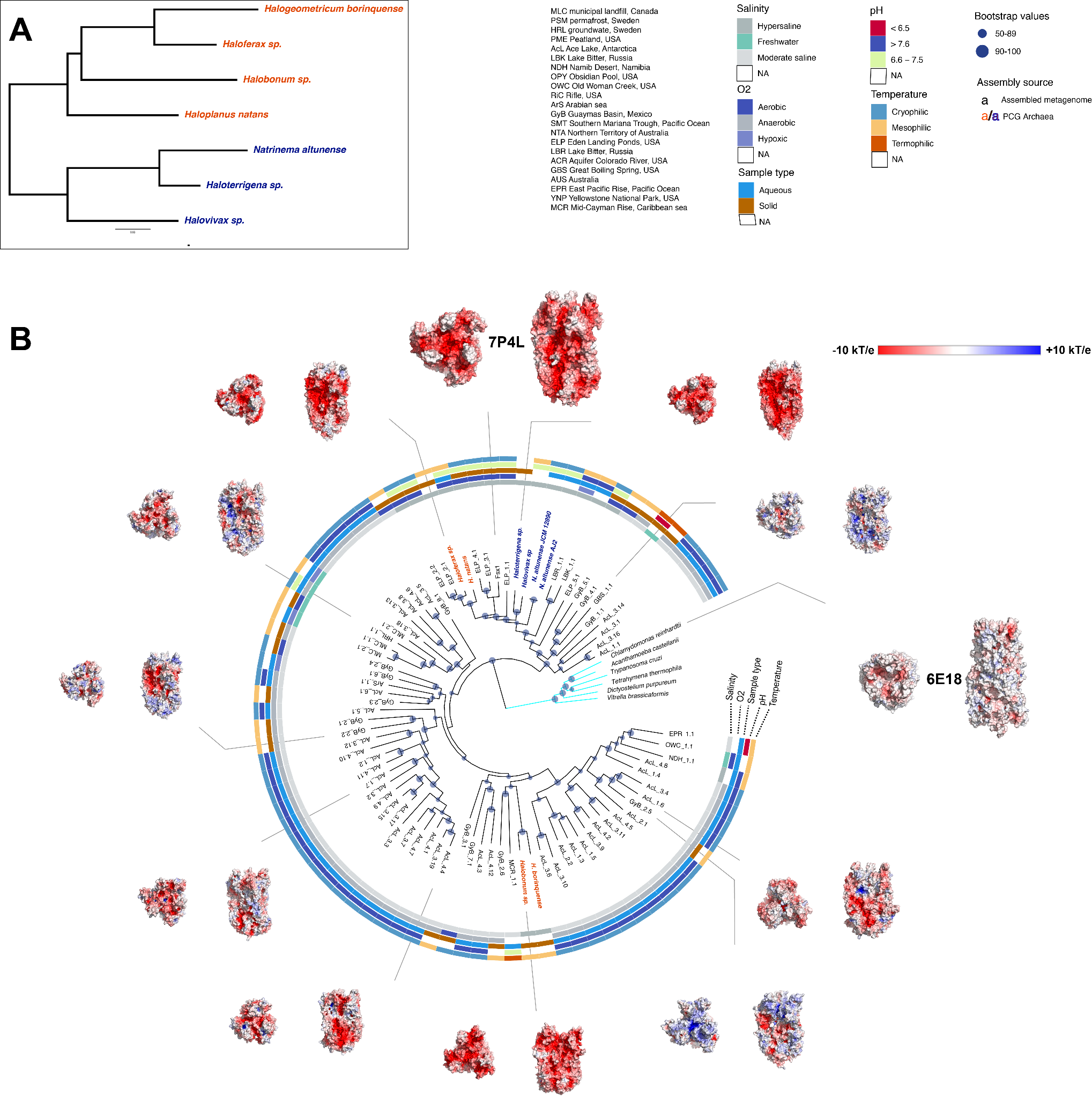

**Fig. S17. Maximum Likelihood phylogenetic tree of Archaea/Eukarya fusexins in context.** (**A**) Panel with phylogenomic tree for *fsx1*-containing cultured archaeal species. (**B**) Unrooted phylogenetic tree represented along with environmental features, a selection of trimers generated by homology modeling (*124*) and two X–ray structures (7P4L and 6E18 (*51*), enlarged). Protein IDs from assembled genomes at tree tips were coded with prefixes indicating sampling site, first number for sample id and last number for the sequence (see data S1). Natrialbales order is shown in blue and Haloferacales order in red, Eukaryotic HAP2 clade is highlighted in cyan with species names. Bootstrap values are indicated with blue dots. Sequences with less than four environmental descriptors were removed from the phylogenetic computation, except if coming from complete genomes. Despite the great diversity of environments where the sequences come from, all of them display uncharged (hydrophobic) tips, even when differing in surface electrostatics for the rest of the molecule. Indeed, for many closely related sequences coming from different environments, surface charge differences are readily apparent. Molecular surfaces coloured by electrostatic potential from negative red (-10 kT/e) to positive blue (+10 kT/e), through neutral/hydrophobic white. All surface representations are oriented as in Fig. 2A. For details on archaeal protein IDs, metagenomics data sources and environmental descriptors refer to data S1.

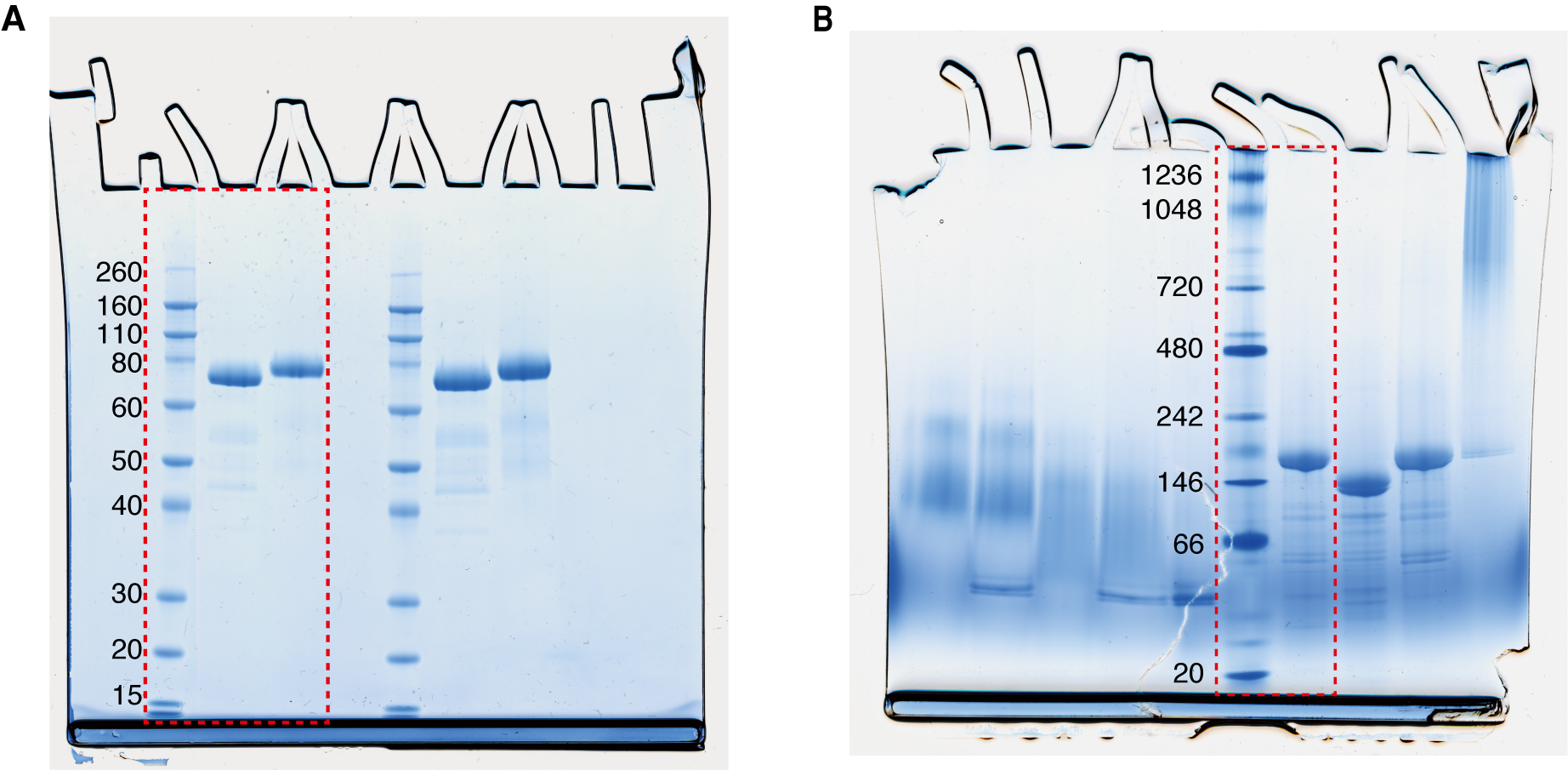

**Fig. S18. Uncropped gel images from fig. S3.** (**A**) SDS-PAGE gel stained with Coomassie G-250. The section indicated by a red dashed square is used in fig. S3A. (**B**) BN-PAGE gel stained with Coomassie G-250. The section indicated by a red dashed square is used in fig. S3B.

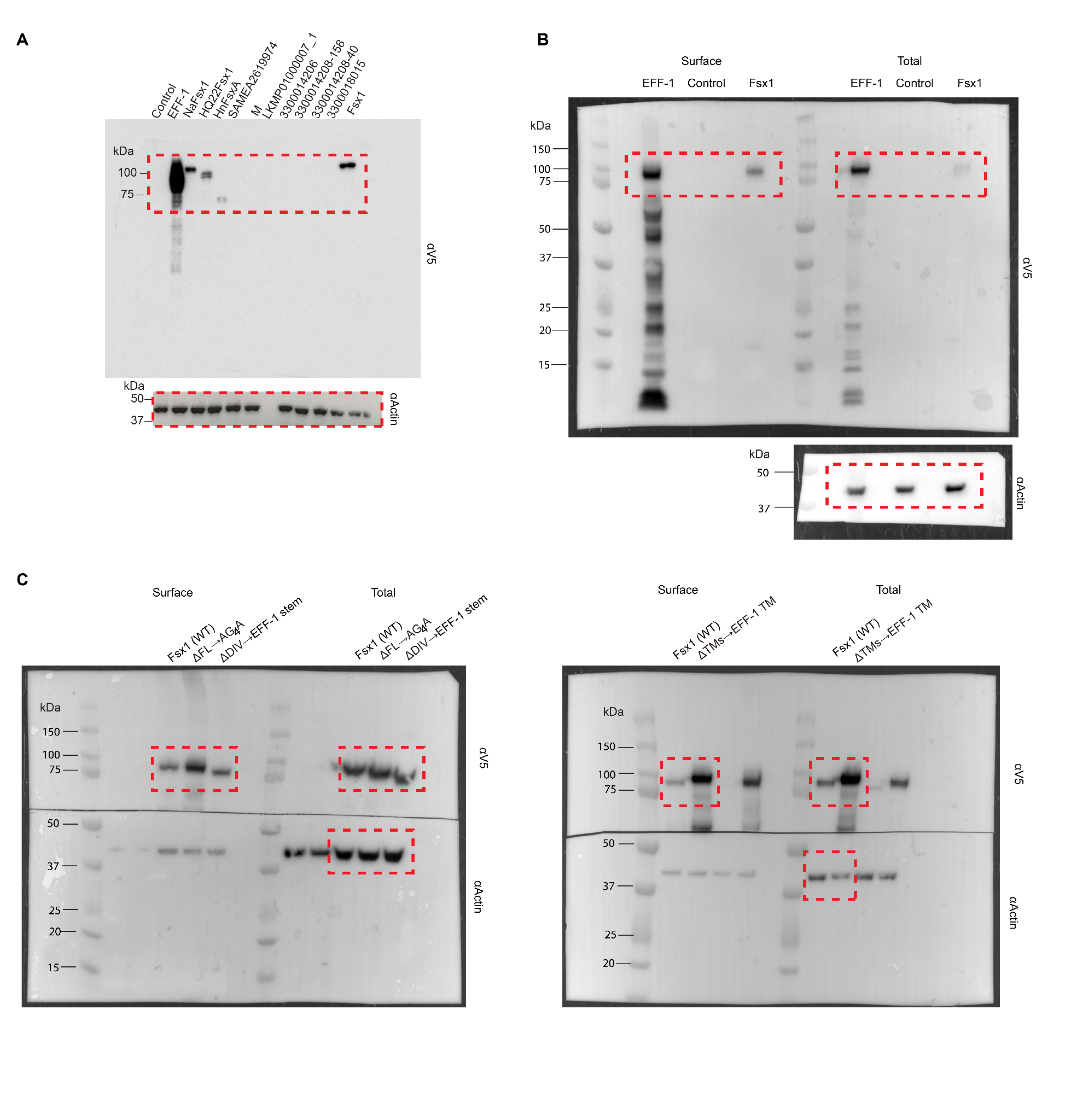

**Fig. S19. Uncropped Western blot images from figs. S2B and S10, B and C.** (**A**) Western blot probed with anti-V5 (upper) and anti-actin (lower) antibodies. The sections indicated by red dashed squares are used in fig. S2B. (**B)** Western blot probed with anti-V5 (upper) and anti-actin (lower) antibodies. The sections indicated by red dashed squares are used in fig. S10B. (**C)** Western blots probed with anti-V5 (upper) and anti-actin (lower) antibodies. The sections indicated by red dashed squares are used in fig. S10C.

### **Supplementary Tables**

| Table S1. Fsx1s in genomes with assigned taxonomy | | |
| --- | --- | --- |
| Fsx1s in Pure Culture Genomes (PCGs) | |  |
| Sequence ID | Species | Taxonomy |
| WP_058826362.1 | *Haloferax* sp*.* Q22 | Archaea › Euryarchaeota › Stenosarchaea group › Halobacteria › Haloferacales › Haloferacaceae |
| WP_144240185.1 | *Natrinema altunense* (AJ2) | Archaea › Euryarchaeota › Stenosarchaea group › Halobacteria › Natrialbales › Natrialbaceae |
| ELY83688.1 | *Natrinema altunense* JCM12890) | Archaea › Euryarchaeota › Stenosarchaea group › Halobacteria › Natrialbales › Natrialbaceae |
| WP_157573584.1 | *Haloplanus natans* DSM 17983 | Archaea › Euryarchaeota › Stenosarchaea group › Halobacteria › Haloferacales › Haloferacaceae |
| WP_174701778.1 | *Haloterrigena* sp*.* SYSU A121-1 | Archaea › Euryarchaeota › Stenosarchaea group › Halobacteria › Natrialbales › Natrialbaceae |
| WP_179268568.1 | *Halobonum* sp*.* NJ-3-1 | Archaea › Euryarchaeota › Stenosarchaea group › Halobacteria › Haloferacales › Halorubraceae |
| WP_163487151.1 | *Halogeometricum borinquense* strain wsp4 | Archaea › Euryarchaeota › Stenosarchaea group › Halobacteria › Haloferacales › Haloferacaceae |
| WP_207587115.1 | *Halovivax* sp. KZCA124 | Archaea › Euryarchaeota › Stenosarchaea group › Halobacteria › Natrialbales › Natrialbaceae |
| Fsx1s in Metagenome-Assembled Genomes (MAGs) | | |
| Sequence ID | Assigned taxon | Taxonomy |
| MGYP000598426430 | Halobacteriales | Archaea › Euryarchaeota › Stenosarchaea group › Halobacteria › Halobacteriales |
| LKMP01000007_1 | Nanohaloarchaea archaeon B1-Br10_U2g21 | Archaea › Euryarchaeota › Stenosarchaea group › Candidatus Nanohaloarchaeota |
| RLG58774.1 | Candidatus Geothermarchaeota B85_G16 | Archaea ›TACK group › Candidatus Geothermarchaeota |
| RLI53188.1 | Candidatus Thorarchaeota archaeon | Archaea › Asgard group › Candidatus Thorarchaeota |
| RKX41251.1 | Thermotogae bacterium | Bacteria › Thermotogae |
| RKZ11204.1 | Candidatus Fermentibacteria bacterium | Bacteria › Candidatus Fermentibacteria |
| RLG94066.1 | Candidatus Bathyarchaeota archaeon | Archaea ›TACK group › Candidatus Bathyarchaeota |
| AJF63093.1 | archaeon GW2011_AR20 | Archaea ›unclassified |
| HEX32987.1 | Candidatus Aenigmarchaeota archaeon | Archaea › DPANN group › Candidatus Aenigmarchaeota |
| HDD44259.1 | Candidatus Desulfofervidus auxilii | Bacteria › Proteobacteria › Deltaproteobacteria › Candidatus Desulfofervidaceae |
| HDI72891.1 | Candidatus Altiarchaeales archaeon | Archaea › DPANN group › Candidatus Altiarchaeota › Candidatus Altiarchaeales |
| HHR27186.1 | Candidatus Bathyarchaeota archaeon | Archaea ›TACK group › Candidatus Bathyarchaeota |
| NJD53946.1 | Candidatus Methanoperedens sp. | Archaea › Euryarchaeota › Stenosarchaea group › Methanomicrobia › Methanosarcinales › Cand. Methanoperedenaceae |
| NOZ47386.1 | Chlorobi bacterium | Bacteria › Chlorobi |
| HGF63239.1 | Candidatus Micrarchaeota archaeon | Archaea › DPANN group › Candidatus Micrarchaeota |
| HID09282.1 | Candidatus Micrarchaeota archaeon | Archaea › DPANN group › Candidatus Micrarchaeota |

###### Table S2. Fsx1_E_ data collection and refinement statistics

| **Data collection** |  |
| --- | --- |
| Space group | *C*2 (5) |
| Cell dimensions  *a*, *b*, *c* (Å)  *α*, *β*, *γ* (°) | 262.51, 111.33, 68.51  90, 100.709, 90 |
| Wavelength (Å) | 1.005 |
| Resolution range (Å) | 67.3-2.3 (2.38-2.30)^*^ |
| Unique reflections | 85618 (8536) |
| Multiplicity | 4.2 (4.4) |
| Completeness (%) | 99.5 (99.9) |
| Mean I / σI | 11.4 (1.4) |
| Wilson B-factor | 42.8 |
| R_merge_  R_meas_  R_pim_ | 0.100 (1.218)  0.115 (1.385)  0.055 (0.651) |
| CC_1/2_ | 1.0 (0.58) |
| CC* | 1.0 (0.86) |
| **Refinement** |  |
| Reflections used in refinement | 85574 (6092)^**^ |
| Reflections used for R_free_ | 2012 (149) |
| R_work_  R_free_ | 0.199 (0.303)  0.243 (0.377) |
| Number of non-H atoms  macromolecules / ligand / solvent | 11733  11051 / 66 / 616 |
| Protein residues | 1432 |
| RMS  bonds (Å)  angles (°) | 0.004  0.58 |
| Ramachandran favoured / allowed / outliers (%) | 98.9 / 1.1 / 0.0 |
| Rotamer outliers (%) | 0.2 |
| Clashscore | 2.5 |
| Average B-factor  macromolecules / ligand / solvent | 53.4  53.5 / 58.8 / 50.0 |

* Values in parenthesis are for the highest resolution shell

** The highest resolution shell used in refinement included reflections between 2.36 and 2.30 Å

| **Table S3. Most common arCOGs from 11 IMEs^a^** | | | | | |
| --- | --- | --- | --- | --- | --- |
| Collapsed Group Name | IMEs count^b^ | arCOG count^c^ | arCOG | Category | Annotation |
| CG_2 | 11 | 22 | arCOG00280, arCOG00285, arCOG06224 | L | HerA helicase |
| CG_17 | 10 | 21 | arCOG01241, arCOG01248, arCOG01250 | X | XerC XerD/XerC family integrase |
| 41684 | 10 | 3 | arCOG01680, arCOG02808, arCOG04362 | K | Transcriptional regulator containing HTH domain |
| 43214 | 9 | 9 | arCOG08903 | S | Uncharacterized protein |
| CG_1 | 8 | 8 | arCOG12186 | S | Uncharacterized protein |
| CG_2 | 7 | 7 | arCOG04816 | U | TraG/TraD/VirD4 family enzyme, ATPase |
| 43797 | 7 | 7 | arCOG12187 | S | Uncharacterized membrane protein |
| 43810 | 7 | 7 | arCOG10296 | S | Uncharacterized membrane protein |
| 43833 | 7 | 7 | arCOG08907 | S | Uncharacterized protein |
| 43868 | 7 | 7 | arCOG10381 | S | Uncharacterized protein |
| CG_2 | 6 | 7 | arCOG00467 | L | Cdc6-related protein, AAA superfamily ATPase |
| CG_2 | 6 | 6 | arCOG07496 | U | VirB4, Type IV secretory pathway |
| 42763 | 6 | 6 | arCOG06216 | K | Transcriptional regulator |
| CG_2 | 5 | 5 | arCOG01308 | O | ATPase of the AAA+ class , CDC48 family |
| CG_21 | 3 | 3 | arCOG07871 | K | Helicase |
| CG_2 | 2 | 2 | arCOG00439 | L | ATPase involved in replication control |
| CG_2 | 2 | 2 | arCOG03779 | V | GTPase subunit of restriction endonuclease |
| CG_2 | 2 | 2 | arCOG05935 | R | Helicase of FtsK superfamily |
| CG_21 | 2 | 2 | arCOG00878 | V | restriction-modification related helicase |
| CG_9 | 1 | 1 | arCOG03600 | E | Transglutaminase-like cysteine protease |
| CG_21 | 1 | 1 | arCOG04818 | K | Superfamily II DNA/RNA helicase, SNF2 family |

a. Collapsed Homology Groups (fig. S14) corresponding to the 11 IME clusters (fig. S15) were analyzed using HMMER against the arCOGs database. Collapsed Homology Groups with zero identified arCOGs are not shown. Full dataset with results is in data S3.

b. Number of IMEs where the indicated arCOGs are present.

c. Total number of ORFs belonging to arCOGs in the next column identified in the set of 11 IMEs.

| **Table S4. IMEs carrying the *fsx1* gene in completely sequenced genomes** | | | | | | | | | | | | | |
| --- | --- | --- | --- | --- | --- | --- | --- | --- | --- | --- | --- | --- | --- |
| **Species with *fsx1*** | **NCBI TaxID** | **PATRIC ID** | **sequence ID^a^** | **ME start k-mer^b^** | **ME end k-mer^b^** | **Length ME k-mer** | **ME start CG^c^** | **ME end CG^c^** | **Length ME CG** | **Species w/o *fsx1*^d^** | **NCBI TaxID** | **PATRIC ID** | **ANI^e^** |
| *Haloplanus natans* DSM 17983 | 926690 | 926690.3 | ATYM01000002 | 1422500 | 1526500 | 104001 | 1422548 | 1535558 | 113011 | *Haloplanus* sp*.* CBA1112 | 1547898 | 1547898.3 | 88.0 |
| *Natrinema altunense* strain AJ2 | 222984 | 222984.5 | JNCS01000001 | 496500 | 593500 | 97001 | 497005 | 591892 | 94888 | *Natrinema altunense* strain 4.1R | 222984 | 222984.10 | 98.0 |
| *Halobonum* sp*.* NJ-3-1 | 2743089 | 2743089.3 | CP058579 | 1918000 | 2078500 | 160501 | 1906722 | 2078630 | 171909 | *Halobonum* sp*.* Gai3-2 | 2743090 | 2743090.3 | 85.0 |
| *Haloferax* sp*.* Q22 | 1526048 | 1526048.3 | LOEP01000012 | 1 | 56511 | 56511 | 1 | 56511 | 56511 | *Haloferax gibbonsii* strain LR2-5 | 35746 | 35746.12 | 94.8 |
| *Haloterrigena* sp. SYSU A121-1 | 2496101 | 2496101.3 | JABURA010000001 | 1761500 | 1860500 | 99001 | 1761690 | 1860499 | 98810 | *Haloterrigena turkmenica* DSM 5511 | 543526 | 543526.13 | 92.4 |
| *Halogeometricum borinquense* strain wsp4 | 60847 | 60847.21 | CP048739 | 2797000 | 2827000 | 30001 | 2763703 | 2853374 | 89672 | *Halogeometricum borinquense* strain wsp3 | 60847 | 60847.22 | 99.6 |
| *Halovivax* sp*.* KZCA124 | 2817025 | 2817025.3 | NZ_CP071597 | 3085000 | 3179500 | 94501 | --**^f^** | -- | -- | -- | -- | -- | -- |

a. Genomic contig carrying the IME.

b. Sequence coordinates of the start and end of the IME identified by k-mer spectrum.

c. Sequence coordinates of the start and end of the IME identified by comparative genomics (CG).

d. Closest species not carrying the *fsx1* gene with a completely sequenced genome (used for CG analysis).

e. Average nucleotide identity between the complete genomes of compared species.

f. No species with a completely sequenced genome was similar enough to *Halovivax* sp*.* KZCA124 to compute an accurate estimate of IME’s insertion sites with CG.

| **Table S5. Synthesized archaeal fusexin genes for fusogenic tests in mammalian cells** | | |
| --- | --- | --- |
| **Synthesized plasmids** | **Accession number/ sequence** | **Species/Assembly source** |
| pBPT01 | WP_007110832 | *Natrinema altunense* |
| pBPT02 | WP_058826362 | *Haloferax* sp*.*Q22 |
| pBPT03 | WP_049937247 | *Haloplanus natans* |
| pBPT04 | SAMEA2619974_10776_4 | MAG |
| pBPT05 | LKMP01000007_1 | Nanohaloarchaea archaeon B1-Br10_U2g21 LB-BRINE-C121 |
| pBPT06 | 3300014206-Ga0172377-10000119-870930-129 | unassembled metagenome |
| pBPT07 | 3300014208-Ga0172379-10000243-871512-158 | unassembled metagenome |
| pBPT08 | 3300014208-Ga0172379-10001592-871560-40 | unassembled metagenome |
| pBPT10 | 3300018015-Ga0187866_1000629\|915963_9 | unassembled metagenome |
| pBPT11 | 3300000868-JGI12330J12834-1000008-299010-8 , **Fsx1** | unassembled metagenome |

| **Table S6. Plasmids used in this study** | | | |
| --- | --- | --- | --- |
| **Plasmid name** | **Description** | **Use** | **Source** |
| pLJFX11B | pLJ6-Fsx1_E_ | For SAXS and SEC-MALS (fig. S3) | this study, amplified from pBPT11 |
| pLJFX11B_T369C | pLJ6-Fsx1_E_-T369C mutant | For crystallographic study (Figs. 1-2; figs. S3-S7) | this study, amplified from pBPT11 |
| pBPT01 | *Natrinema altunense fsx1* synthesized into pGene/V5-His | Inducible expression in mammalian cells (fig. S2) | this study |
| pBPT02 | *Haloferax* sp. Q22 *fsx1* synthesized into pGene/V5-His | Inducible expression in mammalian cells (fig. S2) | this study |
| pBPT03 | *Haloplanus natans fsx1* synthesized into pGene/V5-His | Inducible expression in mammalian cells (fig. S2) | this study |
| pBPT04 | SAMEA2619974 synthesized into pGene/V5-His | Inducible expression in mammalian cells (fig. S2) | this study |
| pBPT05 | LKMP01000007_1 synthesized into pGene/V5-His | Inducible expression in mammalian cells (fig. S2) | this study |
| pBPT06 | 3300014206 synthesized into pGene/V5-His | Inducible expression in mammalian cells (fig. S2) | this study |
| pBPT07 | 3300014208-158 synthesized into pGene/V5-His | Inducible expression in mammalian cells (fig. S2) | this study |
| pBPT08 | 3300014208-40 synthesized into pGene/V5-His | Inducible expression in mammalian cells (fig. S2) | this study |
| pBPT10 | 3300018015 synthesized into pGene/V5-His | Inducible expression in mammalian cells (fig. S2) | this study |
| pBPT11 | *fsx1* synthesized into pGene/V5-His | Inducible expression in mammalian cells (fig. S2) | this study |
| pGene/V5-His | pGene/V5-His | GeneSwitch™ inducible Mammalian Expression | INVITROGEN |
| pSwitch | pSwitch | Regulatory vector for Mifepristone induction | INVITROGEN |
| pOA34 | pGene::EFF-1-V5 | *C. elegans eff-1* fused to a C-terminal V5 tag (EFF-1-V5) in pGene | Avinoam et al., 2011(*11*) |
| pCI H2B-RFP | pCI::H2B-RFP | A CAG promoter (CMV immediate early enhancer and chicken beta actin promoter) and IRES controlled Histone2B-mRFP1 reporter. | Addgene plasmid # 92398(*144*) |
| pCI H2B-GFP | pCI::H2B-GFP | A CAG promoter (CMV immediate early enhancer and chicken beta actin promoter) and IRES controlled Histone2B-EGFP reporter. | Addgene plasmid # 92399(*144*) |
| pCAGIG | pCAGIG | A CAG promoter (CMV immediate early enhancer and chicken beta actin promoter) and IRES controlled EGFP reporter. | Addgene plasmid # 11159(*145*) |
| pNB25 | pCAGIGnes | Intermediate construct to create pNB32 | this study |
| pNB32 | pCI::GFPnes | Content-mixing, Fig. 3A-C | this study |
| pRFPnes | DsRed2 with a nuclear export signal | Content-mixing, Fig. 3A-C | Avinoam et al., 2011(*11*) |
| pXL27 | pCI::Fsx1-V5::H2B-RFP | Content-mixing, Fig. 3A-C | this study |
| pXL28 | pCI::Fsx1-V5::H2B-GFP | Content-mixing, Fig. 3A-C; live imaging of fusion, Fig. 3G | this study |
| pXL29 | pCI::AtHAP2-V5::H2B-RFP | Content-mixing, Fig. 3A-C; Multinucleation assay (fig. S2) | this study |
| pXL30 | pCI::AtHAP2-V5::H2B-GFP | Content-mixing, Fig. 3A-C; Multinucleation assay (fig. S2) | this study |
| pXL49 | pCI::Fsx1-V5::GFPnes | Content-mixing, Fig. 3D-E | this study |
| pNB34 | pCI::EFF-1-V5::GFPnes | Content-mixing, Fig. 3D-E | this study |
| pXL68 | pCI::VSV-G::GFPnes | Content-mixing, Fig. 3D-E | this study |
| pOA19 | pCAGGS::EFF-1-V5 | Surface biotinylation of EFF-1 (fig. S10) | Avinoam et al., 2011(*11*) |
| pXL50 | pCAGGS::Fsx1-V5 | Surface biotinylation of Fsx1 (fig. S10) | this study |
| myr-mCherry | myr-mCherry | mCherry linked to a myristoylated and palmitoylated peptide, live imaging of fusion (Fig. 3G) | Dunsing et al., Sci. Rep. 2018 (*103*) |
| myr-EGFP | myr-EGFP | EGFP linked to a myristoylated and palmitoylated peptide (fig. S2) | Dunsing et al., Sci. Rep. 2018 (*103*) |
| pXL21 | pCI::NaFsx1-V5::H2B-RFP | Multinucleation assay (fig. S2) | this study |
| pXL22 | pCI::NaFsx1-V5::H2B-GFP | Multinucleation assay (fig. S2) | this study |
| pXL23 | pCI::HQ22Fsx1-V5::H2B-RFP | Multinucleation assay (fig. S2) | this study |
| pXL24 | pCI::HQ22Fsx1-V5::H2B-GFP | Multinucleation assay (fig. S2) | this study |
| pXL25 | pCI::HnFsx1-V5::H2B-RFP | Multinucleation assay (fig. S2) | this study, subcloned from pBPT03 with modification of complete signal peptide |
| pXL26 | pCI::HnFsx1-V5::H2B-GFP | Multinucleation assay (fig. S2) | this study, subcloned from pBPT03 with modification of complete signal peptide |
| pXL57 | pCI::Fsx1-ΔFL-AG_4_A::GFPnes | Content-mixing, Fig. 3J | this study |
| **Plasmid name** | **Description** | **Use** | **Source** |
| pXL58 | pCI::Fsx1-ΔFL-AG_4_A::H2B-RFP | Content-mixing, Fig. 3J | this study |
| pXL63 | pCI::Fsx1-ΔDIV-EFF-1-stem::H2B-RFP | Content-mixing, Fig. 3J | this study |
| pXL64 | pCI::Fsx1-ΔDIV-EFF-1-stem::GFPnes | Content-mixing, Fig. 3J | this study |
| pXL108 | pCAGGS::Fsx1-ΔDIV-EFF-1-stem | Surface biotinylation of Fsx1-ΔDIV-EFF-1-stem mutant (fig. S10) | this study |
| pXL82 | pCI::Fsx1-WT-FLAG-3TMs::H2B-RFP | Surface expression tests, FLAG tag inserted before the first TM segment of Fsx1 (fig. S10) | this study |
| pXL86 | pCI::Fsx1-ΔFL-AG_4_A-FLAG-3TMs::H2B-RFP | Surface expression tests, FLAG tag inserted before the first TM segment of Fsx1-ΔFL-AG_4_A mutant (fig. S10) | this study |
| pXL92 | pCI::Fsx1-ΔDIV-EFF-1-stem-FLAG-3TMs::H2B-RFP | Surface expression tests, FLAG tag inserted before the first TM segment of Fsx1-ΔDIV-EFF-1-stem mutant (fig. S10) | this study |
| pOA20 | pCAGGS::AFF-1-FLAG | *C. elegans aff-1* fused to a C-terminal FLAG tag (AFF-1-FLAG) in pCAGGS | Avinoam et al., 2011(*11*) |
| pXL100 | pCI::AFF-1-FLAG::H2B-RFP | Surface expression tests of AFF-1 (fig. S10) | this study |
| pXL106 | pCAGGS-Fsx1-ΔFL-AG_4_A | Surface biotinylation of Fsx1-ΔFL-AG_4_A mutant (fig. S10) | this study |
| pXL123 | pCAGGS-Fsx1ΔTMs-EFF-1 TM | Surface biotinylation of Fsx1ΔTMs-EFF-1 TM mutant (fig. S10) | this study |
| pXL119 | pCI::Fsx1ΔTMs-EFF-1 TM::H2B-RFP | Content-mixing, Fig. 3J; Surface expression tests, FLAG tag inserted before the EFF-1 TM region (fig. S10) | this study |
| pXL120 | pCI::Fsx1ΔTMs-EFF-1 TM::GFPnes | Content-mixing, Fig. 3J | this study |
| pXL115 | pCI::Fsx1ΔTMs-GPI::H2B-RFP | Content-mixing, Fig. 3J; Surface expression tests, FLAG tag inserted before the GPI (fig. S10) | this study |
| pXL116 | pCI::Fsx1ΔTMs-GPI::GFPnes | Content-mixing, Fig. 3J | this study |
| pXL117 | pCI::AtHAP2ΔTM-GPI::H2B-RFP | Content-mixing, Fig. 3J; Surface expression tests, FLAG tag inserted before the GPI (fig. S10) | this study |
| pXL118 | pCI::AtHAP2ΔTM-GPI::GFPnes | Content-mixing, Fig. 3J | this study |

| **Table S7. Primers used in this study** | | |
| --- | --- | --- |
| **Primer name** | **Sequence** | **Description** |
| SNFX11_F | CGTAGCTGAAACCGGTGATTCAATCACGTATAACTCTGG | Forward primer for cloning Fsx1_E_ and T369C mutant into pLJ6 with AgeI |
| SNFX11_R | GGTGATGGTGCTCGAGGGAACCAGAACCTCCGAA | Reverse primer for cloning Fsx1_E_ and T369C mutant into pLJ6 with XhoI |
| SNFX11_T369C_F | ACTGTAtgcGCCACCGTTGAGAATGTC | Forward primer for T369C mutant |
| SNFX11_T369C_R | GGTGGCgcaTACAGTTCCCTCATCTCCCTC | Reverse primer for T369C mutant |
| seq_up | GCTGGTTGTTGTGCTGTCTCATC | Sequencing primer for pLJ6 |
| seq_down | CACCAGCCACCACCTTCTGATAG | Sequencing primer for pLJ6 |
| LXH1 | GATGGTGCGATTGCGGAT | Sequencing primer for pBPT01 |
| LXH2 | CCTACGAGAATGGGCAGA | Sequencing primer for pBPT01 |
| LXH3 | TTGCTGGCAGAGAAATGA | Sequencing primer for pBPT02 |
| LXH4 | TGATGTACCCCGAGTTCA | Sequencing primer for pBPT02 |
| LXH5 | GGATGAAATCTTCAGAAC | Sequencing primer for pBPT03 |
| LXH6 | ACTGTCTCGAAGCCGGTT | Sequencing primer for pBPT03 |
| LXH7 | CAAAATCACCCTCACATC | Sequencing primer for pBPT04 |
| LXH8 | CCTACAATATTAAGTTGTG | Sequencing primer for pBPT04 |
| LXH9 | TGAGTCTGAATGGATTAT | Sequencing primer for pBPT05 |
| LXH10 | TAGGACTACAGCGAAAAT | Sequencing primer for pBPT05 |
| LXH11 | TTGGGGAGGAAATGTAAA | Sequencing primer for pBPT06 |
| LXH12 | TAGAAGAATAAATATTCC | Sequencing primer for pBPT06 |
| LXH13 | TCCTCTTCCCTCGGAGAA | Sequencing primer for pBPT07 |
| LXH14 | CCTACTCAGGTAACGTAA | Sequencing primer for pBPT07 |
| LXH15 | CAGTAACAATAAATGGTG | Sequencing primer for pBPT08 |
| LXH16 | CAGAAGAATAAACATTCC | Sequencing primer for pBPT08 |
| LXH17 | AACAATAGGACAAGCAAA | Sequencing primer for pBPT10 |
| LXH18 | ACCAAAAATATTGTCTGC | Sequencing primer for pBPT10 |
| LXH19 | AGCATACATAGACAACCC | Sequencing primer for pBPT11, Fsx1 |
| LXH20 | ACGTCGATGCCGGAGAAA | Sequencing primer for pBPT11, Fsx1 |
| pGene FW | CTGCTCAACCTTCCTATC | pGene backbone sequencing forward primer |
| pGene REV | TTAGGAAAGGACAGTGGGAGTG | pGene backbone sequencing reverse primer |
| PCA-5 | GGTTCGGCTTCTGGCGTGTGACC | pCI::H2B-RFP/H2B-GFP/GFPnes backbones sequencing forward primer |
| IRES-REV | GCATTCCTTTGGCGAGAG | pCI::H2B-RFP/H2B-GFP/GFPnes backbones sequencing reverse primer |
| pCAGGS FW | GCAACGTGCTGGTTGTTGTGCTGTC | pCAGGS backbone sequencing forward primer |
| pCAGGS RV | TCCCATATGTCCTTCCGAGTGA | pCAGGS backbone sequencing reverse primer |
| LXH24 | CGGGGTACCATGAGACGTGCAGCATTG | Forward primer for cloning Fsx1 and its mutants into pCAGGS vector with KpnI |
| LXH42 | CTAGCTAGCGGTACCATGAGACGTGCAGCATTGATT | Forward primer for cloning Fsx1 into pCI::H2B-RFP/H2B-GFP/GFPnes vectors with NheI and KpnI |
| LXH44 | CTAGCTAGCGGTACCATGGAACCGCCGTTTGAGTGG | Forward primer for cloning EFF-1 into pCI::GFPnes vector with NheI and KpnI |
| LXH45 | CTAGCTAGCGGTACCATGGTGAACGCGATTTTAATG | Forward primer for cloning AtHAP2 into pCI::H2B-RFP/GFP vectors with NheI and KpnI |
| LXH79 | TCCCCCGGGCTAATGGTGATGGTGATGATGACC | Reverse primer for cloning fusexins into pCI vectors with SmaI which binds to 6xHis tag |
| LXH81 | CTAGCTAGCTCAATGGTGATGGTGATGATGACC | Reverse primer for cloning Fsx1 and its mutants into pCAGGS vector with NheI |
| LXH111 | CTAGCTAGCATGAAGTGCCTTTTGTACTTAG | Forward primer for cloning VSV-G into pCI::GFPnes vector with NheI |
| LXH112 | TCCCCCGGGTTACTTTCCAAGTCGGTTCATC | Reverse primer for cloning VSV-G into pCI::GFPnes vector with SmaI |
| **Primer name** | **Sequence** | **Description** |
| LXH39 | CTAGCTAGCGGTACCATGCGGGCGGTGTCTGATTTC | Forward primer for cloning NaFsx1 into pCI::H2B-RFP/GFP vectors with NheI and KpnI |
| LXH40 | CTAGCTAGCGGTACCATGAAAAACGGGTTGAAGGCC | Forward primer for cloning HQ22Fsx1 into pCI::H2B-RFP/GFP vectors with NheI and KpnI |
| LXH134 | CTAGCTAGCATGGTGAAACGAGTGGGTAATTGTTGG  AAGGCCTCAGTAGCGGCATTCTTCCTTCTCATGTTCACTGCATTT | Forward primer for cloning HnFsx1 into pCI::H2B-RFP/GFP vectors with NheI; containing modification of complete signal peptide |
| LXH107 | GCGGAAGGTACAGCAGGTACGCCGGTGGAGGTGGAGCTGATTACGAGATCTATTGTTT | Forward primer for cloning downstream of Fsx1-ΔFL-AG_4_A mutant by overlap PCR |
| LXH108 | AAACAATAGATCTCGTAATCAGCTCCACCTCCACCGGCGTACCTGCTGTACCTTCCGC | Reverse primer for cloning upstream of Fsx1-ΔFL-AG_4_A mutant by overlap PCR |
| LXH101 | ACCGGTATCCAGCAGGAAATCGATCTTGTT | Forward primer 1 for cloning Fsx1-ΔDIV-EFF-1-stem by overlap PCR |
| LXH102 | AACAAGATCGATTTCCTGCTGGATACCGGT | Reverse primer 1 for cloning Fsx1-ΔDIV-EFF-1-stem by overlap PCR |
| LXH103 | ATGATTGCTACGGATCAGGACGATGATTCA | Forward primer 2 for cloning Fsx1-ΔDIV-EFF-1-stem by overlap PCR |
| LXH104 | TGAATCATCGTCCTGATCCGTAGCAATCAT | Reverse primer 2 for cloning Fsx1-ΔDIV-EFF-1-stem by overlap PCR |
| LXH128 | TGTTCGGAGGTTCTGGTTCCGACTACAAGGACGACGATGACAAAGGAGATCTGCTTAC | Forward primer for cloning downstream of Fsx1-WT/mutants-FLAG by overlap PCR |
| LXH129 | GGAACCAGAACCTCCGAACA | Reverse primer for cloning upstream of Fsx1-WT/mutants-FLAG by overlap PCR |
| LXH135 | CTAGCTAGCATGGTACTGTGGCAATGGTCAATAG | Forward primer for cloning *C. elegans* AFF-1-FLAG into pCI::H2B-RFP vector with NheI |
| LXH136 | TCCCCCGGGTTATTTGTCATCGTCGTCCTTGTAGTC | Reverse primer for cloning *C. elegans* AFF-1-FLAG into pCI::H2B-RFP vector with SmaI |
| LXH140 | GACGGGTAGTACCTGAAGTGGTTCCACTTCCTTTATTTGGAGAACCTCCTTTGTCATCGTCGTCCTTGTAGTCGGAACCAGAACCTCCG | Reverse primer 1 for fusing GPI signal from DAF to Fsx1 |
| LXH141 | TCCCCCGGGCTAAGTCAGCAAGCCCATGGTTACTAGCGTCCCAAGCAAACCTGTCAACGTGAAACACGTGTGCCCAGATAGAAGACGGGTAGTACCTGAAGTG | Reverse primer 2 for fusing GPI signal from DAF to Fsx1; Reverse primer for cloning downstream of AtHAP2ΔTM-GPI mutant by overlap PCR |
| LXH142 | AATACGTTTGCTTAAGCTGGGGTGGAAGCGACTACAAGGACGACGAT | Forward primer for cloning downstream of AtHAP2ΔTM-GPI mutant by overlap PCR |
| LXH143 | CCAGCTTAAGCAAACGTATT | Reverse primer for cloning upstream of AtHAP2ΔTM-GPI mutant by overlap PCR |
| LXH144 | GGAGGTTCTGGTTCCGACTACAAGGACGACGATGACAAAATTGTTGTGTATCTC | Forward primer for cloning downstream of Fsx1ΔTMs-EFF-1 TM mutant by overlap PCR which contains one FLAG tag |

**Auxiliary Supplementary Materials**

**Captions for Movies S1 to S2**

**Movie S1** | Time-lapse experiment using spinning disk confocal microscopy reveals merging of two cells expressing myr-mCherry and Fsx1. Time in hours:minutes. Merge of the red and DIC channels is shown.

**Movie S2** | Z-series of the binucleated BHK cell from Fig. 3H. Labeled nuclei (blue) and myr-mCherry (white). Each optical section obtained with spinning disc confocal microscopy is 1 μm apart.

**Captions for Data S1 to S5**

**Data S1_Fsx1s.xlsx.** This table contains a complete list of the Fsx1-encoding genes identified in this study with ID, amino acid sequence and associated environmental and genomic data.

**Data S2_PCGs_IME_ORFs.xlsx.** This table contains a complete list of PCGs’ IME ORFs annotated with identified Pfam, TIGR and arCOGs domains.

**Data S3_11_IME_ORFs.xlsx.** This table contains a complete list of the 11 IME ORFs (as shown in fig. S16) annotated with identified arCOGs domains.

**Data S4_SourceData_Fig.3_and_fig.S2.xlsx.** This file contains the raw data and the statistical information for the functional analyses (i.e. multinucleation and content mixing assays).

**Data S5_SourceData_Synthesized_Fsx1s.docx.** This file contains the sequences of the ten *fsx1s* genes synthesized for expression tests and fusogenic activities.
