## Supplementary material for "Discovery of archaeal Fusexins homologous to eukaryotic HAP2/GCS1 gamete fusion proteins": Data S5

Synthesized plasmid: pBPT01

NZ_JNCS01000001.1:501753-503768 Natrinema altunense strain AJ2 N_altunense_AJ2_contig_1, whole genome shotgun sequence 5’ to 3’

>N.altFusexin1-V56HIS (pBPT01)EDITED

**pBPT01 Edited : Addition of restriction Ez. Sites**

**5’ KpnI, 3’ SacI, SphI, SpeI**

ggtaccATGCGGGCGGTGTCTGATTTCCTGAAGAATAAGTGGGTAGCAGTACCTGCTGTCGCTCTTCTAATCCTCAGTCTCGGTTTTCTGGCGCAGAACTATATTACAGGGTCGTTTGTCAGCGGTGATCAGATAGAGTATACCAGTAATTCGGAGTTCTTCGATGGCGAGGTCTTCGTCATCAACTATATCGGTGACAGGACGACCGACAAGATACACGCGGAGATCCCGGCATCTGAGTTAGCCAGTGCTGCTGACGGAGAAGTCACGAAGGAATTGACGATCGACATCACCTCGCAAGAGACTTACGCCCGGTATAGTACGACATCGACGAGCCTGAGCAGGATTTACGCTTTCAAGGCGATCAATTCGGAGGTGTTTACTGATACGTCTTCTTCGGAGGTTGAGTCTCGGAAGCAGTCTTGGGCGGAGCAGAACTGCTACGATATAGACGGTGATTCGGAGATTGAGCAGGGTGATGATTACACTCAGCGGTTCTGGGTGGATGGTGCGATTGCGGATTACTACAATGCTCAGGTGTACTGTGTTCGGGAAAATGGATACTATGGGAATATCGCGGAGTTGAGTTCGCCTGACAAGGAGTTTGAGGCAGAGGTCGAGGTGAAGGCGGATGGTGAGACACCTCAGTCGACCACTCTAAGTAACTCAGATCTTGGTCAGGGAAGATGCGACAACATCGGGGAGTATGTCAGGGCTTGCTGGACCGGTTTGAATCCGACCGGTGAAGGAGTTCCTGAGCCGTATAATGTCTACGCAATTCACAGCAACACGTTCTCCCCGGAGTGGAGAGTAATCTCAGCGGAGCGGTACTCGTCGTACAACAGCTATGTCGAGAACAATCTGTACAGTCAGATCGAGGCTTGGAAGGAAGGATCTCTGAGTCAGGAAGAGGTTGTCTCGACTGCGAATGATCAGGCGGCTCAAGCAGCTCAAAGAGTTTCCTCTGGCAGCTTTACTGATTACCAGGTGGAGGACTCTTCCTACGAGAATGGGCAGATCCGGTTGAATCCGAGTTATGATATTGCCTGGCCGGAGTTCACGTTCTACGTGGACGGTGCGGACTATATCTCGGTTGAGAAGCCGGTAGGTGAGCCGAGGATAGTGAATGTTGAAGGAGATGAGTTTGGTGAGAAGCAGTCTGGCACTGTTACAGCCGAAGTGGAGAACACGGCGAGCTACGAAGGAAGTTTCTCTGCCAGGGTTTCCAAGTGTAGCGATAGCTTCGGTTATGATTCGTTGCAGGAGACAAAGACTGTGGGTCCCGGTGAGACTGTTTCCTTCGATTTCCGGGTGAGCTTCACTTCGGATAGTTTCGATCAGAAAACGGTCTCTGGTAGCTGTGAGATCGTGGTTTCTGATACCGGTGATTCCTCAAATAGTGATACCGCGGTAGTGGATGTTGAGGCTACTCAGAGTAATGAGTGTACTCCGAACGAGGAGTTCGTCAGGACGATCAACTCAACTCACTCCAGGATCCTCGAGTGTAGCAGTGATGGTTTGACCACGACCGAAGTAGATGTCTGTGGAGACGGTGAGGAAGCAGCCTTCAGAGATAGCTCTTGGACCTGTGTATCTGATGGAACTCTTCCTGGAGGCGGTGACGGATCCGGAGGTTCCGGTGGCGGAGGAGATACGGATGACTGTGTAGTGTTTGCCGTGGGAGATCTGGAGATTGAGGATCCGTTCTGTGCGGATGGTCCGTTGGAAATGCTGTCGAAGATGTTCCACTTGGTGGCAGGGACAGCGGTAGCGTTCTTCACGGGAAGCCTGGGTTACAGGGCTGGTCGTTGGGTTGACGGCGAGTACCAGATTAAGGGAGGTTTCGACCCGTTAAAATCTCGAAGCGTATCCCGCGCTAAGAGAGGAAGATTCCTGATAGGACTGATCGCTGAATTGGTTTCGTTCCTGCTCGGGTTCTATGTGATCCTCTTGGTGCCGATATGGGCACAGTTGATGGTGATCCTGGGATACGTCCTGTTCAAGTATTATACGCCGTTTCTAGAGGGCCCGCGGTTCGAAGGTAAGCCTATCCCTAACCCTCTCCTCGGTCTCGATTCTACGCGTACCGGTCATCATCACCATCACCATTGAgagctcgcatgcactagt

```
>N.altFusexin1-V56HIS (pBPT01)EDITED
From 1 to 2130.
ggtaccATGCGGGCGGTGTCTGATTTCCTGAAGAATAAGTGGGTAGCAGTACCTGCTGTCGCTCTTCTAATCCTCAGTCTCGGTTTTCTG 90
       M  R  A  V  S  D  F  L  K  N  K  W  V  A  V  P  A  V  A  L  L  I  L  S  L  G  F  L  30

GCGCAGAACTATATTACAGGGTCGTTTGTCAGCGGTGATCAGATAGAGTATACCAGTAATTCGGAGTTCTTCGATGGCGAGGTCTTCGTC 180
 A  Q  N  Y  I  T  G  S  F  V  S  G  D  Q  I  E  Y  T  S  N  S  E  F  F  D  G  E  V  F  V  60

ATCAACTATATCGGTGACAGGACGACCGACAAGATACACGCGGAGATCCCGGCATCTGAGTTAGCCAGTGCTGCTGACGGAGAAGTCACG 270
 I  N  Y  I  G  D  R  T  T  D  K  I  H  A  E  I  P  A  S  E  L  A  S  A  A  D  G  E  V  T  90

AAGGAATTGACGATCGACATCACCTCGCAAGAGACTTACGCCCGGTATAGTACGACATCGACGAGCCTGAGCAGGATTTACGCTTTCAAG 360
 K  E  L  T  I  D  I  T  S  Q  E  T  Y  A  R  Y  S  T  T  S  T  S  L  S  R  I  Y  A  F  K  120

GCGATCAATTCGGAGGTGTTTACTGATACGTCTTCTTCGGAGGTTGAGTCTCGGAAGCAGTCTTGGGCGGAGCAGAACTGCTACGATATA 450
 A  I  N  S  E  V  F  T  D  T  S  S  S  E  V  E  S  R  K  Q  S  W  A  E  Q  N  C  Y  D  I  150

GACGGTGATTCGGAGATTGAGCAGGGTGATGATTACACTCAGCGGTTCTGGGTGGATGGTGCGATTGCGGATTACTACAATGCTCAGGTG 540
 D  G  D  S  E  I  E  Q  G  D  D  Y  T  Q  R  F  W  V  D  G  A  I  A  D  Y  Y  N  A  Q  V  180

TACTGTGTTCGGGAAAATGGATACTATGGGAATATCGCGGAGTTGAGTTCGCCTGACAAGGAGTTTGAGGCAGAGGTCGAGGTGAAGGCG 630
 Y  C  V  R  E  N  G  Y  Y  G  N  I  A  E  L  S  S  P  D  K  E  F  E  A  E  V  E  V  K  A  210

GATGGTGAGACACCTCAGTCGACCACTCTAAGTAACTCAGATCTTGGTCAGGGAAGATGCGACAACATCGGGGAGTATGTCAGGGCTTGC 720
 D  G  E  T  P  Q  S  T  T  L  S  N  S  D  L  G  Q  G  R  C  D  N  I  G  E  Y  V  R  A  C  240

TGGACCGGTTTGAATCCGACCGGTGAAGGAGTTCCTGAGCCGTATAATGTCTACGCAATTCACAGCAACACGTTCTCCCCGGAGTGGAGA 810
 W  T  G  L  N  P  T  G  E  G  V  P  E  P  Y  N  V  Y  A  I  H  S  N  T  F  S  P  E  W  R  270

GTAATCTCAGCGGAGCGGTACTCGTCGTACAACAGCTATGTCGAGAACAATCTGTACAGTCAGATCGAGGCTTGGAAGGAAGGATCTCTG 900
 V  I  S  A  E  R  Y  S  S  Y  N  S  Y  V  E  N  N  L  Y  S  Q  I  E  A  W  K  E  G  S  L  300

AGTCAGGAAGAGGTTGTCTCGACTGCGAATGATCAGGCGGCTCAAGCAGCTCAAAGAGTTTCCTCTGGCAGCTTTACTGATTACCAGGTG 990
 S  Q  E  E  V  V  S  T  A  N  D  Q  A  A  Q  A  A  Q  R  V  S  S  G  S  F  T  D  Y  Q  V  330

GAGGACTCTTCCTACGAGAATGGGCAGATCCGGTTGAATCCGAGTTATGATATTGCCTGGCCGGAGTTCACGTTCTACGTGGACGGTGCG 1080
 E  D  S  S  Y  E  N  G  Q  I  R  L  N  P  S  Y  D  I  A  W  P  E  F  T  F  Y  V  D  G  A  360

GACTATATCTCGGTTGAGAAGCCGGTAGGTGAGCCGAGGATAGTGAATGTTGAAGGAGATGAGTTTGGTGAGAAGCAGTCTGGCACTGTT 1170
 D  Y  I  S  V  E  K  P  V  G  E  P  R  I  V  N  V  E  G  D  E  F  G  E  K  Q  S  G  T  V  390

ACAGCCGAAGTGGAGAACACGGCGAGCTACGAAGGAAGTTTCTCTGCCAGGGTTTCCAAGTGTAGCGATAGCTTCGGTTATGATTCGTTG 1260
 T  A  E  V  E  N  T  A  S  Y  E  G  S  F  S  A  R  V  S  K  C  S  D  S  F  G  Y  D  S  L  420

CAGGAGACAAAGACTGTGGGTCCCGGTGAGACTGTTTCCTTCGATTTCCGGGTGAGCTTCACTTCGGATAGTTTCGATCAGAAAACGGTC 1350
 Q  E  T  K  T  V  G  P  G  E  T  V  S  F  D  F  R  V  S  F  T  S  D  S  F  D  Q  K  T  V  450

TCTGGTAGCTGTGAGATCGTGGTTTCTGATACCGGTGATTCCTCAAATAGTGATACCGCGGTAGTGGATGTTGAGGCTACTCAGAGTAAT 1440
 S  G  S  C  E  I  V  V  S  D  T  G  D  S  S  N  S  D  T  A  V  V  D  V  E  A  T  Q  S  N  480

GAGTGTACTCCGAACGAGGAGTTCGTCAGGACGATCAACTCAACTCACTCCAGGATCCTCGAGTGTAGCAGTGATGGTTTGACCACGACC 1530
 E  C  T  P  N  E  E  F  V  R  T  I  N  S  T  H  S  R  I  L  E  C  S  S  D  G  L  T  T  T  510

GAAGTAGATGTCTGTGGAGACGGTGAGGAAGCAGCCTTCAGAGATAGCTCTTGGACCTGTGTATCTGATGGAACTCTTCCTGGAGGCGGT 1620
 E  V  D  V  C  G  D  G  E  E  A  A  F  R  D  S  S  W  T  C  V  S  D  G  T  L  P  G  G  G  540

GACGGATCCGGAGGTTCCGGTGGCGGAGGAGATACGGATGACTGTGTAGTGTTTGCCGTGGGAGATCTGGAGATTGAGGATCCGTTCTGT 1710
 D  G  S  G  G  S  G  G  G  G  D  T  D  D  C  V  V  F  A  V  G  D  L  E  I  E  D  P  F  C  570

GCGGATGGTCCGTTGGAAATGCTGTCGAAGATGTTCCACTTGGTGGCAGGGACAGCGGTAGCGTTCTTCACGGGAAGCCTGGGTTACAGG 1800
 A  D  G  P  L  E  M  L  S  K  M  F  H  L  V  A  G  T  A  V  A  F  F  T  G  S  L  G  Y  R  600

GCTGGTCGTTGGGTTGACGGCGAGTACCAGATTAAGGGAGGTTTCGACCCGTTAAAATCTCGAAGCGTATCCCGCGCTAAGAGAGGAAGA 1890
 A  G  R  W  V  D  G  E  Y  Q  I  K  G  G  F  D  P  L  K  S  R  S  V  S  R  A  K  R  G  R  630

TTCCTGATAGGACTGATCGCTGAATTGGTTTCGTTCCTGCTCGGGTTCTATGTGATCCTCTTGGTGCCGATATGGGCACAGTTGATGGTG 1980
 F  L  I  G  L  I  A  E  L  V  S  F  L  L  G  F  Y  V  I  L  L  V  P  I  W  A  Q  L  M  V  660

ATCCTGGGATACGTCCTGTTCAAGTATTATACGCCGTTTCTAGAGGGCCCGCGGTTCGAAGGTAAGCCTATCCCTAACCCTCTCCTCGGT 2070
 I  L  G  Y  V  L  F  K  Y  Y  T  P  F  L  E  G  P  R  F  E  G  K  P  I  P  N  P  L  L  G  690

CTCGATTCTACGCGTACCGGTCATCATCACCATCACCATTGAgagctcgcatgcactagt 2130
 L  D  S  T  R  T  G  H  H  H  H  H  H  *                    710
```

Synthesized plasmid: pBPT02

>HaloferaxQ22Fusexin1-V5-6HIS pBPT02-edited

**Addition of restriction Ez. Sites**

**5’ KpnI, 3’ SacI, SphI, SpeI**

KpnI edited GGTACC to GGTATC both encode for Tyr

ggtaccATGAAAAACGGGTTGAAGGCCTCGGTAGTAGCACTGTTTTTCCTATTAGCCGCTTCTTCTGCATCAGCAGCGACGAACTCTGTTGACACAGTCACCTACGAGAGCAACAGTGATTTCTTCGACGGACAGGTTCTCCAAGTAGGTTATACCTCTAATTTTGCTACGGACAAGATCAACGTATATCTCTCTGAAAGCAAGATCGAGAGTGAGACTGGAGCGGTAGCGGATGAACCGATTACGTTCGATGTTTCTCACCAGAATACGTACGCCAAGTACCCAACTCAGGATACAGGTTTAGAGTCGATAACTGGTTGGGACGCAGTCAAGAAGACTGTCAGCTCTAAGCAGGAGCTGTGGGACTGGACTACCACGAACTGTGTCGATACGAACGATGCTGGCACGTTCACAGTCGATGGTTACGGGACGACTGATGTAGATGCTCGAGCGAACAGATGGTTCGATACCTGGACAGGGCAGTACAAGTATACAATCTATTGCTGGCAGAGAAATGATTACTACGGTGCTGTAGCTGACATCGGTTCTCCTGACGAAGTCTTCCGGACAGAGTGGAGGCTTCAGGCCGGAGATAAGAATCCTCAGACAGCTGTGATCACTAACGGCGACGGCGGAACAGGAGTGACCTCTAACCTCGGCAGATATGCCAAGGTAAGCTGGGAGGGAAGTCTTTCAACCGGTGAGAATCCTCCGCTCGTCGATGATGAATATGCTCTCCACAGTAATGACTACGAAGGCGGTTGGAGAGTCATAAGCGAATCACGTTACTCCTCGTACCAGAGTTTCGTCCAGAACAATGCTGACGACCTTCTTGGAGAATGGGGTGCAGGCCTAACCACTGAGAGCCACATCGAGAGCGAGATGAACGGAAAAGCGGAAGAAGCTGCTTCGGAGTTTACGGAGTCTCCGCTTTCTGACTCCGAGATCTTGGACTCGAGCTTCCAGAATGGTGCGTTCAAACTGGATATGGAGCGTAGCCTGATGTACCCCGAGTTCAGTGTCTACGTTGATGCGGGTGAGAACGGTTATGCCACGATATCTAAGCCTGTAGGTAAGCCAGAGATCGTCTCTACAAGCGGTGCAGAGTTCGGTGAATTGGGTCAGGGAACGATATCAGTGGAAGCAGAAAATGTCGGTGATAGCGAAGGCAGTTTCTCGGCTCGTACCGATTCCTGCGGCGAGTACTTCAGCGGTAACAGCTTGCAGGATACTCAAAGGGTTTCTCCTGGCGGTACTGCATCGTTTGATTTCCGGGTGACGTTCACTTCGACGAGTATGACGCAGAGTGAGTTCTCTGACCAGTGTGAAGTTGTTGTTGAGGATACTGGGAGTGGGAACGAGGTCTCTGCCAGTGTCTCTGTTACTGCTACTCAGGAAGATGAGTGTACAGAGGGAGATGAGACGAAGAAGGAGAAACAGGTAAACGGTGAAACGGTCGACGTCATCATGTCCTGTACTAATGGCCTGAAACTGGAGGAGGACGAAGTCTGCAGTGCTGATGAGGAAGCTCGATACATTGACGACGATGTCCAGTATGAATGCCGTGACAAGGACTCGCCTCCCGGTGGAGGCGGCGGTGATCCGTGGACTATTCCAGGCCTCGGATGGCAACTAGATACTCCATTTGGCGGTATGGAGAAGGCCTTATCCGGTAACGCTGGTGCATTGACCTGGGCTCAATTGTTGCTGTCGTTCATCGGATTCCTCGCTGGATTCGCCTTAGTAGGGGTTAAGCTCGGTAAGATGGTTGACGGCCTTGCCACGGAGTTCATCCCATTGAGTGATGCCGTCGTGAGACTGGGTATAGGACTTGTAGGAGGCGGGATGGCGTTCATGGCTGTGTATCAGTTAGTGACGAATCCGTTAGGGTTCTTGCTGACTGTTGTCGGCCTGCTTCTAACAGGGTAtCTGTATTTGAAGGGTACTACTCCGGACATTAACTTGCTAGAGGGCCCGCGGTTCGAAGGTAAGCCTATCCCTAACCCTCTCCTCGGTCTCGATTCTACGCGTACCGGTCATCATCACCATCACCATTGAgagctcgcatgcactagt

```
>HaloferaxQ22Fusexin1-V5-6HIS pBPT02-edited
From 1 to 2076.
ggtaccATGAAAAACGGGTTGAAGGCCTCGGTAGTAGCACTGTTTTTCCTATTAGCCGCTTCTTCTGCATCAGCAGCGACGAACTCTGTT 90
       M  K  N  G  L  K  A  S  V  V  A  L  F  F  L  L  A  A  S  S  A  S  A  A  T  N  S  V  30

GACACAGTCACCTACGAGAGCAACAGTGATTTCTTCGACGGACAGGTTCTCCAAGTAGGTTATACCTCTAATTTTGCTACGGACAAGATC 180
 D  T  V  T  Y  E  S  N  S  D  F  F  D  G  Q  V  L  Q  V  G  Y  T  S  N  F  A  T  D  K  I  60

AACGTATATCTCTCTGAAAGCAAGATCGAGAGTGAGACTGGAGCGGTAGCGGATGAACCGATTACGTTCGATGTTTCTCACCAGAATACG 270
 N  V  Y  L  S  E  S  K  I  E  S  E  T  G  A  V  A  D  E  P  I  T  F  D  V  S  H  Q  N  T  90

TACGCCAAGTACCCAACTCAGGATACAGGTTTAGAGTCGATAACTGGTTGGGACGCAGTCAAGAAGACTGTCAGCTCTAAGCAGGAGCTG 360
 Y  A  K  Y  P  T  Q  D  T  G  L  E  S  I  T  G  W  D  A  V  K  K  T  V  S  S  K  Q  E  L  120

TGGGACTGGACTACCACGAACTGTGTCGATACGAACGATGCTGGCACGTTCACAGTCGATGGTTACGGGACGACTGATGTAGATGCTCGA 450
 W  D  W  T  T  T  N  C  V  D  T  N  D  A  G  T  F  T  V  D  G  Y  G  T  T  D  V  D  A  R  150

GCGAACAGATGGTTCGATACCTGGACAGGGCAGTACAAGTATACAATCTATTGCTGGCAGAGAAATGATTACTACGGTGCTGTAGCTGAC 540
 A  N  R  W  F  D  T  W  T  G  Q  Y  K  Y  T  I  Y  C  W  Q  R  N  D  Y  Y  G  A  V  A  D  180

ATCGGTTCTCCTGACGAAGTCTTCCGGACAGAGTGGAGGCTTCAGGCCGGAGATAAGAATCCTCAGACAGCTGTGATCACTAACGGCGAC 630
 I  G  S  P  D  E  V  F  R  T  E  W  R  L  Q  A  G  D  K  N  P  Q  T  A  V  I  T  N  G  D  210

GGCGGAACAGGAGTGACCTCTAACCTCGGCAGATATGCCAAGGTAAGCTGGGAGGGAAGTCTTTCAACCGGTGAGAATCCTCCGCTCGTC 720
 G  G  T  G  V  T  S  N  L  G  R  Y  A  K  V  S  W  E  G  S  L  S  T  G  E  N  P  P  L  V  240

GATGATGAATATGCTCTCCACAGTAATGACTACGAAGGCGGTTGGAGAGTCATAAGCGAATCACGTTACTCCTCGTACCAGAGTTTCGTC 810
 D  D  E  Y  A  L  H  S  N  D  Y  E  G  G  W  R  V  I  S  E  S  R  Y  S  S  Y  Q  S  F  V  270

CAGAACAATGCTGACGACCTTCTTGGAGAATGGGGTGCAGGCCTAACCACTGAGAGCCACATCGAGAGCGAGATGAACGGAAAAGCGGAA 900
 Q  N  N  A  D  D  L  L  G  E  W  G  A  G  L  T  T  E  S  H  I  E  S  E  M  N  G  K  A  E  300

GAAGCTGCTTCGGAGTTTACGGAGTCTCCGCTTTCTGACTCCGAGATCTTGGACTCGAGCTTCCAGAATGGTGCGTTCAAACTGGATATG 990
 E  A  A  S  E  F  T  E  S  P  L  S  D  S  E  I  L  D  S  S  F  Q  N  G  A  F  K  L  D  M  330

GAGCGTAGCCTGATGTACCCCGAGTTCAGTGTCTACGTTGATGCGGGTGAGAACGGTTATGCCACGATATCTAAGCCTGTAGGTAAGCCA 1080
 E  R  S  L  M  Y  P  E  F  S  V  Y  V  D  A  G  E  N  G  Y  A  T  I  S  K  P  V  G  K  P  360

GAGATCGTCTCTACAAGCGGTGCAGAGTTCGGTGAATTGGGTCAGGGAACGATATCAGTGGAAGCAGAAAATGTCGGTGATAGCGAAGGC 1170
 E  I  V  S  T  S  G  A  E  F  G  E  L  G  Q  G  T  I  S  V  E  A  E  N  V  G  D  S  E  G  390

AGTTTCTCGGCTCGTACCGATTCCTGCGGCGAGTACTTCAGCGGTAACAGCTTGCAGGATACTCAAAGGGTTTCTCCTGGCGGTACTGCA 1260
 S  F  S  A  R  T  D  S  C  G  E  Y  F  S  G  N  S  L  Q  D  T  Q  R  V  S  P  G  G  T  A  420

TCGTTTGATTTCCGGGTGACGTTCACTTCGACGAGTATGACGCAGAGTGAGTTCTCTGACCAGTGTGAAGTTGTTGTTGAGGATACTGGG 1350
 S  F  D  F  R  V  T  F  T  S  T  S  M  T  Q  S  E  F  S  D  Q  C  E  V  V  V  E  D  T  G  450

AGTGGGAACGAGGTCTCTGCCAGTGTCTCTGTTACTGCTACTCAGGAAGATGAGTGTACAGAGGGAGATGAGACGAAGAAGGAGAAACAG 1440
 S  G  N  E  V  S  A  S  V  S  V  T  A  T  Q  E  D  E  C  T  E  G  D  E  T  K  K  E  K  Q  480

GTAAACGGTGAAACGGTCGACGTCATCATGTCCTGTACTAATGGCCTGAAACTGGAGGAGGACGAAGTCTGCAGTGCTGATGAGGAAGCT 1530
 V  N  G  E  T  V  D  V  I  M  S  C  T  N  G  L  K  L  E  E  D  E  V  C  S  A  D  E  E  A  510

CGATACATTGACGACGATGTCCAGTATGAATGCCGTGACAAGGACTCGCCTCCCGGTGGAGGCGGCGGTGATCCGTGGACTATTCCAGGC 1620
 R  Y  I  D  D  D  V  Q  Y  E  C  R  D  K  D  S  P  P  G  G  G  G  G  D  P  W  T  I  P  G  540

CTCGGATGGCAACTAGATACTCCATTTGGCGGTATGGAGAAGGCCTTATCCGGTAACGCTGGTGCATTGACCTGGGCTCAATTGTTGCTG 1710
 L  G  W  Q  L  D  T  P  F  G  G  M  E  K  A  L  S  G  N  A  G  A  L  T  W  A  Q  L  L  L  570

TCGTTCATCGGATTCCTCGCTGGATTCGCCTTAGTAGGGGTTAAGCTCGGTAAGATGGTTGACGGCCTTGCCACGGAGTTCATCCCATTG 1800
 S  F  I  G  F  L  A  G  F  A  L  V  G  V  K  L  G  K  M  V  D  G  L  A  T  E  F  I  P  L  600

AGTGATGCCGTCGTGAGACTGGGTATAGGACTTGTAGGAGGCGGGATGGCGTTCATGGCTGTGTATCAGTTAGTGACGAATCCGTTAGGG 1890
 S  D  A  V  V  R  L  G  I  G  L  V  G  G  G  M  A  F  M  A  V  Y  Q  L  V  T  N  P  L  G  630

TTCTTGCTGACTGTTGTCGGCCTGCTTCTAACAGGGTAtCTGTATTTGAAGGGTACTACTCCGGACATTAACTTGCTAGAGGGCCCGCGG 1980
 F  L  L  T  V  V  G  L  L  L  T  G  Y  L  Y  L  K  G  T  T  P  D  I  N  L  L  E  G  P  R  660

TTCGAAGGTAAGCCTATCCCTAACCCTCTCCTCGGTCTCGATTCTACGCGTACCGGTCATCATCACCATCACCATTGAgagctcgcatgc 2070
 F  E  G  K  P  I  P  N  P  L  L  G  L  D  S  T  R  T  G  H  H  H  H  H  H  *              690

actagt 2076
       692
```

Synthesized plasmid: pBPT03

NZ_KE386573.1:2594348-2596255 Haloplanus natans DSM 17983 HALNADRAFT_scaffold1.1, whole genome shotgun sequence

>H.natansFusexin1-V5-6His (pBPT03)EDITED (SacI-KpnI)

**Addition of restriction Ez. Sites**

**5’ KpnI, 3’ SacI, SphI, SpeI**

KpnI: edited GGTACC to GGTACT both encoding for Thr

SacI: GAGCTC to GAGCTT both encoding for Leu

ggtaccATGTTCACTGCATTTGCGGCAGCAGATACTACTTCGGTGGATACTGTCTCGTATAAGAGCAACAGCGAGTTCTTCGACGGAGAAGTTCTTCAGGCTGGTTATGTCTCGAACTTTGCGACTGACAAGATTAACGTCCATCTCGGTGCCAGTGAGATTGAGAGCAAGACCGGAGCTACTGCGGAAGAAGACTTGACTTTGAGTGTTTCTCACCAGAACACCTACGCTCGCTACCCGCTCCAGGAGACCGGTTTGAAGAAGATCTACGGCTGGGAGGGGATGAAGAAGACATTCGATACGAAAGACCAGCTATGGAACTGGGTTACCTCAAACTGTGCTGACTTCACTGAGGGCGGAGTAGCGATCGGAGATAGAACCAGTAAGGTCGAGGCCGCTGCCAAGAGCTGGTACGATTACTGGACAGGAAGTTACAATTACCAGGCCTTCTGCCTGCGGAAGAACGGGTACTACGGAGATGTAGCTGATATCGGCTCCCCGGATGAAATCTTCAGAACCGAGTGGAGGCTGCAAGCAGGCGATAAGAATCCGCAGACTGCTATAATAACAAATGGGGATGGTGGATCAGGTGTTGTCTCCAATCTTGGCCGGTATGCCAAGGTGAAATGGCAGGGTAACTTGGATACAGGTCAGAATCCTCCGCTTGTCGACGATGAACTTGCCATCCACGGTAACAACTATGAGGGCGGTTGGAGAGTGATCAGTGAGCAACGGTACGATAACTACTGGAACTACATCAAGAACGACGGGAACAGGCTTCTCGATAAGTGGAAGTCCGGAGACTATTCTGAAAGCTATATTGAAGGCCTGTTGAACGGAAAGGCCGAGAACGCTCAGAAGAGGTACAGTGAGTCTCCGTTGGCGAATGCTGAGGTCTTGGATTCCAGCTTCCAGAGTGGAGCATTGAAGGCCGATATGGATAGTCGATTGGCTTACCCAGAGTTCAATGTCTATGTTGATGCCGGTGAGAACGGGTACATCACTGTCTCGAAGCCGGTTGGGAAGCCGAAGATTGTTTCATCGAGTGGTGCCAGTTTCGGTGAGCT**t**AGTAGCGGCAGGATCTCTGTCGACGTGAAGAACGTGGGCGGGGCGGAAGGCAGTTTCTCAGCCCGTGCTGGCAGCTGTAGTCAGTACTTCCAGGCGGATGCTCTTCAGAATACGAAAAGGGTTGCTCCCGGTGAGACTGCGTCCTACAGTTTCCGGGTTACGTTCACTTCGACGAGTATGCAGCAGGCCTCGTATACAGGTCGATGCAGTATCACTGTAGAGGATACGGGGAGTGGTCGAGAGGTTTCGACCAGTGTCTCGGTGGAGGCTACGCAACAGTCTGAATGTACTCAGGGCAAGGAGATTGTGAAGCAGAAGAACGGGAACGACGTCATCTACAGTTGTCCTGACGGTCTCAAGATTCAGAAGCAGGATACGTGTACGGGAGAGTTGAAGGCGGTTTTCGTCAACAACGATATCCAATACGATTGTAGGGAGGAAGGTAC**t**GGTGGTGGATCTGGTGGAGGCCTGTTCGGTAAACGGTTCACTCTCCCAATCACCGGTACTCAACTGAGTAATCCTCTGAATGCTTTCGAGAATATCTGGAGCGGTGACGCGAATGCGTTGAACTGGTTGCAGGTGTTTGTAACGTTCATCGCGTTCCTCGGCGGGTTCGCCTTGGTCGGTGTCAAGCTCGGTAAAATTGTAGACGGACTTGCGACAGAGTTCATCCCGGTAAAGGATTCCCACGTTAGACTGGTGATAGGCCTTCTCGGTGGAGGAATGATCGCAACAGCGGTCTATCAATTGGTCACAGATCCGTTAGGCCTCTTGGTCACGGTACTGGGCTTGGTTGTGATGGGTTATCTGTATCTCAGTGCGAGTGCTCCTGAAATCAACCTCCTAGAGGGCCCGCGGTTCGAAGGTAAGCCTATCCCTAACCCTCTCCTCGGTCTCGATTCTACGCGTACCGGTCATCATCACCATCACCATTGAgagctcgcatgcactagt

```
>H.natansFusexin1-V5-6His  (pBPT03)EDITED
From 1 to 2022.
ggtaccATGTTCACTGCATTTGCGGCAGCAGATACTACTTCGGTGGATACTGTCTCGTATAAGAGCAACAGCGAGTTCTTCGACGGAGAA 90
       M  F  T  A  F  A  A  A  D  T  T  S  V  D  T  V  S  Y  K  S  N  S  E  F  F  D  G  E  30

GTTCTTCAGGCTGGTTATGTCTCGAACTTTGCGACTGACAAGATTAACGTCCATCTCGGTGCCAGTGAGATTGAGAGCAAGACCGGAGCT 180
 V  L  Q  A  G  Y  V  S  N  F  A  T  D  K  I  N  V  H  L  G  A  S  E  I  E  S  K  T  G  A  60

ACTGCGGAAGAAGACTTGACTTTGAGTGTTTCTCACCAGAACACCTACGCTCGCTACCCGCTCCAGGAGACCGGTTTGAAGAAGATCTAC 270
 T  A  E  E  D  L  T  L  S  V  S  H  Q  N  T  Y  A  R  Y  P  L  Q  E  T  G  L  K  K  I  Y  90

GGCTGGGAGGGGATGAAGAAGACATTCGATACGAAAGACCAGCTATGGAACTGGGTTACCTCAAACTGTGCTGACTTCACTGAGGGCGGA 360
 G  W  E  G  M  K  K  T  F  D  T  K  D  Q  L  W  N  W  V  T  S  N  C  A  D  F  T  E  G  G  120

GTAGCGATCGGAGATAGAACCAGTAAGGTCGAGGCCGCTGCCAAGAGCTGGTACGATTACTGGACAGGAAGTTACAATTACCAGGCCTTC 450
 V  A  I  G  D  R  T  S  K  V  E  A  A  A  K  S  W  Y  D  Y  W  T  G  S  Y  N  Y  Q  A  F  150

TGCCTGCGGAAGAACGGGTACTACGGAGATGTAGCTGATATCGGCTCCCCGGATGAAATCTTCAGAACCGAGTGGAGGCTGCAAGCAGGC 540
 C  L  R  K  N  G  Y  Y  G  D  V  A  D  I  G  S  P  D  E  I  F  R  T  E  W  R  L  Q  A  G  180

GATAAGAATCCGCAGACTGCTATAATAACAAATGGGGATGGTGGATCAGGTGTTGTCTCCAATCTTGGCCGGTATGCCAAGGTGAAATGG 630
 D  K  N  P  Q  T  A  I  I  T  N  G  D  G  G  S  G  V  V  S  N  L  G  R  Y  A  K  V  K  W  210

CAGGGTAACTTGGATACAGGTCAGAATCCTCCGCTTGTCGACGATGAACTTGCCATCCACGGTAACAACTATGAGGGCGGTTGGAGAGTG 720
 Q  G  N  L  D  T  G  Q  N  P  P  L  V  D  D  E  L  A  I  H  G  N  N  Y  E  G  G  W  R  V  240

ATCAGTGAGCAACGGTACGATAACTACTGGAACTACATCAAGAACGACGGGAACAGGCTTCTCGATAAGTGGAAGTCCGGAGACTATTCT 810
 I  S  E  Q  R  Y  D  N  Y  W  N  Y  I  K  N  D  G  N  R  L  L  D  K  W  K  S  G  D  Y  S  270

GAAAGCTATATTGAAGGCCTGTTGAACGGAAAGGCCGAGAACGCTCAGAAGAGGTACAGTGAGTCTCCGTTGGCGAATGCTGAGGTCTTG 900
 E  S  Y  I  E  G  L  L  N  G  K  A  E  N  A  Q  K  R  Y  S  E  S  P  L  A  N  A  E  V  L  300

GATTCCAGCTTCCAGAGTGGAGCATTGAAGGCCGATATGGATAGTCGATTGGCTTACCCAGAGTTCAATGTCTATGTTGATGCCGGTGAG 990
 D  S  S  F  Q  S  G  A  L  K  A  D  M  D  S  R  L  A  Y  P  E  F  N  V  Y  V  D  A  G  E  330

AACGGGTACATCACTGTCTCGAAGCCGGTTGGGAAGCCGAAGATTGTTTCATCGAGTGGTGCCAGTTTCGGTGAGCTtAGTAGCGGCAGG 1080
 N  G  Y  I  T  V  S  K  P  V  G  K  P  K  I  V  S  S  S  G  A  S  F  G  E  L  S  S  G  R  360

ATCTCTGTCGACGTGAAGAACGTGGGCGGGGCGGAAGGCAGTTTCTCAGCCCGTGCTGGCAGCTGTAGTCAGTACTTCCAGGCGGATGCT 1170
 I  S  V  D  V  K  N  V  G  G  A  E  G  S  F  S  A  R  A  G  S  C  S  Q  Y  F  Q  A  D  A  390

CTTCAGAATACGAAAAGGGTTGCTCCCGGTGAGACTGCGTCCTACAGTTTCCGGGTTACGTTCACTTCGACGAGTATGCAGCAGGCCTCG 1260
 L  Q  N  T  K  R  V  A  P  G  E  T  A  S  Y  S  F  R  V  T  F  T  S  T  S  M  Q  Q  A  S  420

TATACAGGTCGATGCAGTATCACTGTAGAGGATACGGGGAGTGGTCGAGAGGTTTCGACCAGTGTCTCGGTGGAGGCTACGCAACAGTCT 1350
 Y  T  G  R  C  S  I  T  V  E  D  T  G  S  G  R  E  V  S  T  S  V  S  V  E  A  T  Q  Q  S  450

GAATGTACTCAGGGCAAGGAGATTGTGAAGCAGAAGAACGGGAACGACGTCATCTACAGTTGTCCTGACGGTCTCAAGATTCAGAAGCAG 1440
 E  C  T  Q  G  K  E  I  V  K  Q  K  N  G  N  D  V  I  Y  S  C  P  D  G  L  K  I  Q  K  Q  480

GATACGTGTACGGGAGAGTTGAAGGCGGTTTTCGTCAACAACGATATCCAATACGATTGTAGGGAGGAAGGTACtGGTGGTGGATCTGGT 1530
 D  T  C  T  G  E  L  K  A  V  F  V  N  N  D  I  Q  Y  D  C  R  E  E  G  T  G  G  G  S  G  510

GGAGGCCTGTTCGGTAAACGGTTCACTCTCCCAATCACCGGTACTCAACTGAGTAATCCTCTGAATGCTTTCGAGAATATCTGGAGCGGT 1620
 G  G  L  F  G  K  R  F  T  L  P  I  T  G  T  Q  L  S  N  P  L  N  A  F  E  N  I  W  S  G  540

GACGCGAATGCGTTGAACTGGTTGCAGGTGTTTGTAACGTTCATCGCGTTCCTCGGCGGGTTCGCCTTGGTCGGTGTCAAGCTCGGTAAA 1710
 D  A  N  A  L  N  W  L  Q  V  F  V  T  F  I  A  F  L  G  G  F  A  L  V  G  V  K  L  G  K  570

ATTGTAGACGGACTTGCGACAGAGTTCATCCCGGTAAAGGATTCCCACGTTAGACTGGTGATAGGCCTTCTCGGTGGAGGAATGATCGCA 1800
 I  V  D  G  L  A  T  E  F  I  P  V  K  D  S  H  V  R  L  V  I  G  L  L  G  G  G  M  I  A  600

ACAGCGGTCTATCAATTGGTCACAGATCCGTTAGGCCTCTTGGTCACGGTACTGGGCTTGGTTGTGATGGGTTATCTGTATCTCAGTGCG 1890
 T  A  V  Y  Q  L  V  T  D  P  L  G  L  L  V  T  V  L  G  L  V  V  M  G  Y  L  Y  L  S  A  630

AGTGCTCCTGAAATCAACCTCCTAGAGGGCCCGCGGTTCGAAGGTAAGCCTATCCCTAACCCTCTCCTCGGTCTCGATTCTACGCGTACC 1980
 S  A  P  E  I  N  L  L  E  G  P  R  F  E  G  K  P  I  P  N  P  L  L  G  L  D  S  T  R  T  660

GGTCATCATCACCATCACCATTGAgagctcgcatgcactagt 2022
 G  H  H  H  H  H  H  *                    674
```

*Haloplanus natans* Fsx1 modified with **complete signal peptide**

**ATGGTGAAACGAGTGGGTAATTGTTGGAAGGCCTCAGTAGCGGCATTCTTCCTTCTC**ATGTTCACTGCATTTGCGGCAGCAGATACTACTTCGGTGGATACTGTCTCGTATAAGAGCAACAGCGAGTTCTTCGACGGAGAAGTTCTTCAGGCTGGTTATGTCTCGAACTTTGCGACTGACAAGATTAACGTCCATCTCGGTGCCAGTGAGATTGAGAGCAAGACCGGAGCTACTGCGGAAGAAGACTTGACTTTGAGTGTTTCTCACCAGAACACCTACGCTCGCTACCCGCTCCAGGAGACCGGTTTGAAGAAGATCTACGGCTGGGAGGGGATGAAGAAGACATTCGATACGAAAGACCAGCTATGGAACTGGGTTACCTCAAACTGTGCTGACTTCACTGAGGGCGGAGTAGCGATCGGAGATAGAACCAGTAAGGTCGAGGCCGCTGCCAAGAGCTGGTACGATTACTGGACAGGAAGTTACAATTACCAGGCCTTCTGCCTGCGGAAGAACGGGTACTACGGAGATGTAGCTGATATCGGCTCCCCGGATGAAATCTTCAGAACCGAGTGGAGGCTGCAAGCAGGCGATAAGAATCCGCAGACTGCTATAATAACAAATGGGGATGGTGGATCAGGTGTTGTCTCCAATCTTGGCCGGTATGCCAAGGTGAAATGGCAGGGTAACTTGGATACAGGTCAGAATCCTCCGCTTGTCGACGATGAACTTGCCATCCACGGTAACAACTATGAGGGCGGTTGGAGAGTGATCAGTGAGCAACGGTACGATAACTACTGGAACTACATCAAGAACGACGGGAACAGGCTTCTCGATAAGTGGAAGTCCGGAGACTATTCTGAAAGCTATATTGAAGGCCTGTTGAACGGAAAGGCCGAGAACGCTCAGAAGAGGTACAGTGAGTCTCCGTTGGCGAATGCTGAGGTCTTGGATTCCAGCTTCCAGAGTGGAGCATTGAAGGCCGATATGGATAGTCGATTGGCTTACCCAGAGTTCAATGTCTATGTTGATGCCGGTGAGAACGGGTACATCACTGTCTCGAAGCCGGTTGGGAAGCCGAAGATTGTTTCATCGAGTGGTGCCAGTTTCGGTGAGCTtAGTAGCGGCAGGATCTCTGTCGACGTGAAGAACGTGGGCGGGGCGGAAGGCAGTTTCTCAGCCCGTGCTGGCAGCTGTAGTCAGTACTTCCAGGCGGATGCTCTTCAGAATACGAAAAGGGTTGCTCCCGGTGAGACTGCGTCCTACAGTTTCCGGGTTACGTTCACTTCGACGAGTATGCAGCAGGCCTCGTATACAGGTCGATGCAGTATCACTGTAGAGGATACGGGGAGTGGTCGAGAGGTTTCGACCAGTGTCTCGGTGGAGGCTACGCAACAGTCTGAATGTACTCAGGGCAAGGAGATTGTGAAGCAGAAGAACGGGAACGACGTCATCTACAGTTGTCCTGACGGTCTCAAGATTCAGAAGCAGGATACGTGTACGGGAGAGTTGAAGGCGGTTTTCGTCAACAACGATATCCAATACGATTGTAGGGAGGAAGGTACtGGTGGTGGATCTGGTGGAGGCCTGTTCGGTAAACGGTTCACTCTCCCAATCACCGGTACTCAACTGAGTAATCCTCTGAATGCTTTCGAGAATATCTGGAGCGGTGACGCGAATGCGTTGAACTGGTTGCAGGTGTTTGTAACGTTCATCGCGTTCCTCGGCGGGTTCGCCTTGGTCGGTGTCAAGCTCGGTAAAATTGTAGACGGACTTGCGACAGAGTTCATCCCGGTAAAGGATTCCCACGTTAGACTGGTGATAGGCCTTCTCGGTGGAGGAATGATCGCAACAGCGGTCTATCAATTGGTCACAGATCCGTTAGGCCTCTTGGTCACGGTACTGGGCTTGGTTGTGATGGGTTATCTGTATCTCAGTGCGAGTGCTCCTGAAATCAACCTCCTAGAGGGCCCGCGGTTCGAAGGTAAGCCTATCCCTAACCCTCTCCTCGGTCTCGATTCTACGCGTACCGGTCATCATCACCATCACCATTAG

Synthesized plasmid: pBPT04

SAMEA2619974_10776_4

SAMEA.TARA-Fusexin1-V5-6His (pBPT04) EDITED

**Addition of restriction Ez. Sites**

**5’ KpnI, 3’ SacI, SphI, SpeI**

KpnI edited GGTACC to GGTTCC both encoding for Val

ggtaccATGTATAAAAGGAAGATAATCATACCAATATTATTATTTGTAATATTATCCCTAATAGTTAGTGCTTTCTCAACACTATCACTTGATAGGGTAGATTTTTCTACCAGGGGAGATGAGTTTGACCAACAGTGGTTATTATTAATTAGTGAAGATGGTAGAGCTGATAAAGCAGTGGTAACTAAAAAAGCAGAAGAAATTAAAGATGATAAAATCCACGCTAAAAATGATTTAACTATCAAAACTGAGATAGATAAAAATAGTTGTGTGTATACAATCCAAAATATAAACCAACCTATTAGTAGGTTAGATTATACAAAAAAGACTGTATGGAAATTCTGGGATATACCTAATGAGATAAAGAGTTGTGAGCAAAGAGAAGGGTATTATTGGGCTTTTAACCTTGGATTTGATTTTAATGTTTATTGTTTCTTCACTCCAGCTGGTAAAGATTCGTTTGTTGGTAGGATTTCCGATAAAAAGTATGATTTTAATACCAAAATCACCCTCACATCAGGTGGTGAAAGTAAATCAGTAGATATATCAAACTCAAATAGAGTAAGTAATTCAGTTCCAGATTATTTTGTAGCAAAATGGCACGGTAGTCTGTCAACTGGAGCAGAATGTCCAGATATCACTTCCCTATCTCCCATATATGATAGAGATAGTGGTGAATGGAAGTTGGGAGAAAAATCATCAATTTCTAACTATCAATCTTTTCATTCTTCAGGTATGAAAGAATGTCTTAACAGTTATATAATTGCCAGGACTACTGGAGGACTTATAGTAGAAACACCAGATAGTTGTAAGAAAAGGTATAATGATTTAGTAGAAAAAGCACAGTCACCAGAAGTATTTAATATTAAAGGTTCTACTGGTAGTATGACAACACAATCATCAGAGCTTGGTGAGTTAGTGGTTGAAGATGTTCAACCTTTCCAATTCCCAGTAATAAGTTTGAAAATCAATGCAGATTGGATTGGAATTGTCCAACCTATCACAAAACCTACAATATTAAGTTGTGAAAGTAGTAAGTTTAAACAAGGAGAGATAGGTGAGATAGATGTTGAGGTTAAAAATAAGGGAGAGAGTGGTTCAGTTTCCATATCAACTGTTTGTTTATCTCCCTTTAAATCAGTTGGAACAACACCCATAATAAGGTTAAAGAAGGGTGAAAGTAAATCAATTACTGTTCCAATCACCGTTAGTACAAAAGAAGATGTAAGTAAAACATGTAGTGTAACAGTTCAAGATGAATCAGACCCAAGTGTGAGAGTTAGTAAAAGATGTGATGTTAGTGCAAGTGGAGTTGTATTGTGTGAAGCAGGAAAGAAAAGGTGTAGTGGTAGATTTATAGAACAATGTAAAAGTAGTGGTAGTGAATATGGGTTAATTGAAGAGTGTGAATTAGATTGTAAGTTGGATAAGTACGGACAACCATTTTGTCCAGAAGTTACACCCCCACCTCCACCTCCACCAAACGGAAATGGAGATAAATGTGAACCAATATGGGCAATAGCAAAGATAACTATCATACCAGATTTTATTTGTGAGATGGAGAAGGTtCCTTACCTAAAAGAGGGTATTAGTGGTTTGGTTGGATTTGTGATGTTTATAATATTAATATTTATGTTGAAGAATTTAATCGGATTTGAGAACATACCACAAAGGTTAATGGTATTAGGTTTCTCATTGATAATTGCAATATTATTGGGTTTTTTGTTTTATTATTTGTTTTGGTTTGGAGTTGCATTAATAGTGGCGTTAGTTGTGATGTTTTTTGTAATTAAAATAATCCTCGGAAAGGTGGGGTTGCTAGAGGGCCCGCGGTTCGAAGGTAAGCCTATCCCTAACCCTCTCCTCGGTCTCGATTCTACGCGTACCGGTCATCATCACCATCACCATTGAgagctcgcatgcactagt

SAMEA.TARA-Fusexin1-V5-6His (pBPT04) EDITED

```
From 1 to 1929.
ggtaccATGTATAAAAGGAAGATAATCATACCAATATTATTATTTGTAATATTATCCCTAATAGTTAGTGCTTTCTCAACACTATCACTT 90
       M  Y  K  R  K  I  I  I  P  I  L  L  F  V  I  L  S  L  I  V  S  A  F  S  T  L  S  L  30

GATAGGGTAGATTTTTCTACCAGGGGAGATGAGTTTGACCAACAGTGGTTATTATTAATTAGTGAAGATGGTAGAGCTGATAAAGCAGTG 180
 D  R  V  D  F  S  T  R  G  D  E  F  D  Q  Q  W  L  L  L  I  S  E  D  G  R  A  D  K  A  V  60

GTAACTAAAAAAGCAGAAGAAATTAAAGATGATAAAATCCACGCTAAAAATGATTTAACTATCAAAACTGAGATAGATAAAAATAGTTGT 270
 V  T  K  K  A  E  E  I  K  D  D  K  I  H  A  K  N  D  L  T  I  K  T  E  I  D  K  N  S  C  90

GTGTATACAATCCAAAATATAAACCAACCTATTAGTAGGTTAGATTATACAAAAAAGACTGTATGGAAATTCTGGGATATACCTAATGAG 360
 V  Y  T  I  Q  N  I  N  Q  P  I  S  R  L  D  Y  T  K  K  T  V  W  K  F  W  D  I  P  N  E  120

ATAAAGAGTTGTGAGCAAAGAGAAGGGTATTATTGGGCTTTTAACCTTGGATTTGATTTTAATGTTTATTGTTTCTTCACTCCAGCTGGT 450
 I  K  S  C  E  Q  R  E  G  Y  Y  W  A  F  N  L  G  F  D  F  N  V  Y  C  F  F  T  P  A  G  150

AAAGATTCGTTTGTTGGTAGGATTTCCGATAAAAAGTATGATTTTAATACCAAAATCACCCTCACATCAGGTGGTGAAAGTAAATCAGTA 540
 K  D  S  F  V  G  R  I  S  D  K  K  Y  D  F  N  T  K  I  T  L  T  S  G  G  E  S  K  S  V  180

GATATATCAAACTCAAATAGAGTAAGTAATTCAGTTCCAGATTATTTTGTAGCAAAATGGCACGGTAGTCTGTCAACTGGAGCAGAATGT 630
 D  I  S  N  S  N  R  V  S  N  S  V  P  D  Y  F  V  A  K  W  H  G  S  L  S  T  G  A  E  C  210

CCAGATATCACTTCCCTATCTCCCATATATGATAGAGATAGTGGTGAATGGAAGTTGGGAGAAAAATCATCAATTTCTAACTATCAATCT 720
 P  D  I  T  S  L  S  P  I  Y  D  R  D  S  G  E  W  K  L  G  E  K  S  S  I  S  N  Y  Q  S  240

TTTCATTCTTCAGGTATGAAAGAATGTCTTAACAGTTATATAATTGCCAGGACTACTGGAGGACTTATAGTAGAAACACCAGATAGTTGT 810
 F  H  S  S  G  M  K  E  C  L  N  S  Y  I  I  A  R  T  T  G  G  L  I  V  E  T  P  D  S  C  270

AAGAAAAGGTATAATGATTTAGTAGAAAAAGCACAGTCACCAGAAGTATTTAATATTAAAGGTTCTACTGGTAGTATGACAACACAATCA 900
 K  K  R  Y  N  D  L  V  E  K  A  Q  S  P  E  V  F  N  I  K  G  S  T  G  S  M  T  T  Q  S  300

TCAGAGCTTGGTGAGTTAGTGGTTGAAGATGTTCAACCTTTCCAATTCCCAGTAATAAGTTTGAAAATCAATGCAGATTGGATTGGAATT 990
 S  E  L  G  E  L  V  V  E  D  V  Q  P  F  Q  F  P  V  I  S  L  K  I  N  A  D  W  I  G  I  330

GTCCAACCTATCACAAAACCTACAATATTAAGTTGTGAAAGTAGTAAGTTTAAACAAGGAGAGATAGGTGAGATAGATGTTGAGGTTAAA 1080
 V  Q  P  I  T  K  P  T  I  L  S  C  E  S  S  K  F  K  Q  G  E  I  G  E  I  D  V  E  V  K  360

AATAAGGGAGAGAGTGGTTCAGTTTCCATATCAACTGTTTGTTTATCTCCCTTTAAATCAGTTGGAACAACACCCATAATAAGGTTAAAG 1170
 N  K  G  E  S  G  S  V  S  I  S  T  V  C  L  S  P  F  K  S  V  G  T  T  P  I  I  R  L  K  390

AAGGGTGAAAGTAAATCAATTACTGTTCCAATCACCGTTAGTACAAAAGAAGATGTAAGTAAAACATGTAGTGTAACAGTTCAAGATGAA 1260
 K  G  E  S  K  S  I  T  V  P  I  T  V  S  T  K  E  D  V  S  K  T  C  S  V  T  V  Q  D  E  420

TCAGACCCAAGTGTGAGAGTTAGTAAAAGATGTGATGTTAGTGCAAGTGGAGTTGTATTGTGTGAAGCAGGAAAGAAAAGGTGTAGTGGT 1350
 S  D  P  S  V  R  V  S  K  R  C  D  V  S  A  S  G  V  V  L  C  E  A  G  K  K  R  C  S  G  450

AGATTTATAGAACAATGTAAAAGTAGTGGTAGTGAATATGGGTTAATTGAAGAGTGTGAATTAGATTGTAAGTTGGATAAGTACGGACAA 1440
 R  F  I  E  Q  C  K  S  S  G  S  E  Y  G  L  I  E  E  C  E  L  D  C  K  L  D  K  Y  G  Q  480

CCATTTTGTCCAGAAGTTACACCCCCACCTCCACCTCCACCAAACGGAAATGGAGATAAATGTGAACCAATATGGGCAATAGCAAAGATA 1530
 P  F  C  P  E  V  T  P  P  P  P  P  P  P  N  G  N  G  D  K  C  E  P  I  W  A  I  A  K  I  510

ACTATCATACCAGATTTTATTTGTGAGATGGAGAAGGTtCCTTACCTAAAAGAGGGTATTAGTGGTTTGGTTGGATTTGTGATGTTTATA 1620
 T  I  I  P  D  F  I  C  E  M  E  K  V  P  Y  L  K  E  G  I  S  G  L  V  G  F  V  M  F  I  540

ATATTAATATTTATGTTGAAGAATTTAATCGGATTTGAGAACATACCACAAAGGTTAATGGTATTAGGTTTCTCATTGATAATTGCAATA 1710
 I  L  I  F  M  L  K  N  L  I  G  F  E  N  I  P  Q  R  L  M  V  L  G  F  S  L  I  I  A  I  570

TTATTGGGTTTTTTGTTTTATTATTTGTTTTGGTTTGGAGTTGCATTAATAGTGGCGTTAGTTGTGATGTTTTTTGTAATTAAAATAATC 1800
 L  L  G  F  L  F  Y  Y  L  F  W  F  G  V  A  L  I  V  A  L  V  V  M  F  F  V  I  K  I  I  600

CTCGGAAAGGTGGGGTTGCTAGAGGGCCCGCGGTTCGAAGGTAAGCCTATCCCTAACCCTCTCCTCGGTCTCGATTCTACGCGTACCGGT 1890
 L  G  K  V  G  L  L  E  G  P  R  F  E  G  K  P  I  P  N  P  L  L  G  L  D  S  T  R  T  G  630

CATCATCACCATCACCATTGAgagctcgcatgcactagt 1929
 H  H  H  H  H  H  *                    643
```

Synthesized plasmid: pBPT05

LKMP01000007_1 (Nanohaloarchaea B1-Br10_U2g21 LB-BRINE-C121)

>NanohaloarchFusexin1-V5-6His (pBPT05)EDITED

**Addition of restriction Ez. Sites**

**5’ KpnI, 3’ SacI, SphI, SpeI**

ggtaccATGTCTACGTTTTTGATAGTTCCAGCTTCTGTATCTTATGTATCGGAAGTAGACTATATTGAAACAACTTCAGAGTTTTTTGACCAGGGCCACGTAGTTATCGAATTTGTTATTGATGGCGCTGATGAGAAAATACATGCAGATCTAGAACCTAGCCAGTTATCAGATTTATCTGATCAAAATACGAATCAAAGTGTTGAGATTAGCGCGAAAGTATCCGATGTCTATGCCGAGTATGAAGTAGAAGATAGAGGTCTCAGGCCTTTAAAACCTATAGAATACACGAGAACTACTCATTCCACAAAAGAAAAAGCTCAGATTTGGGCCGATAATAACTGCCTTGATCTTGATAATAGCGGAGCACCCAACTATAATACTTATAGAAGAGGATTGGTTAACCCCTGGGGAGACTGGATTGTGAATTGTGCGATACCTAATATGAGCCAATCTACTTTTAACGTTGGTTTCTTACAGAGTCCTCCAGATACGTTATTTGAGTCTGAATGGATTATAGCTGTTGAAGGTGAATCTGCCGAAACAGTTCGATTATCTAATAGTGGCGCAGAACATGGAAGTTTGGAAAGAGTGAATGATAATGTTCAACTCGAGTTTGTTTCATTGAAAGATACCGGGAGTAATGCTCCTAACCCTCAAAATACTCTAGCTGCTTTTTCAAGCACAGATGGCTGGAGATTAATTGAAAAAGAAGAATTTGATAGCTACATAACTTATATTGAAAATGAATCTCCTAAAAAAGTTGAAGAGATGCGAGAACAAGATAATTTCAATATAGAAGACTGGACTGATAAGAAAAACAAAGAAGCCAGAGATAACTTTTTGAGCGCATCAAGAGAATACGAAGAATCTCCTATTCAAGGGTCAAACGTATCTGATGGCAAAAACAAATTTCAAAATGGGATTTTTCAATATAGGCCTGAATCAGAGTTCGAATGGTTTATCTCAGAGTTTAACGTTAGATTAAATGGTGATTTTGTAGGACTACAGCGAAAATTTGCAGATTTAGAGATTATAAATATAGAAGATACCAAAGTTAATGGTCTTTCAAGCGGTTCTGTGGAAGTAAAAGTTGCAAATAATGGCAATGGAACCGGCGCTTTTGAAGTTTCAGCTTCCTGTGATAAATTTGATACAGTTGGAGTAAGTAGTCGCACATCATTAAAATCTGGAGAATCTGACACAGTATTTCTAAATTTGGATACTGACGGTTCTGAAGAGTTCTATGAAACTTGTAATCTCAATATTAAGGACCTTGAAACTACTTCAATCGTTTCTAGCTCATTTACTGCATCTTACACTCCTGAAAAAGAATGTACACCAGGAAGACAGTCAGTCAGAATAGAGAACGGACAAGAAGCAATTTACGAATGTGCTGAAGACGGAATTTCCATATTTGAGGTTGATGCTTGCGAATTGGACGAAAGAGCTAAACCTTACGGTGAAGGATATGAGTGTCAAAAAGTTGAAAGTCTTCGGGAAATTGAAGCACCTCAAGACTGCGAAATTTCCATACTTCCTAGCTTCCTGAATAACTTATTCGCTAATGACTATACCTTTACAAATCCAATTTGCGCCACGGAAAACAGGTTAAATGATTTTAATAATAGTATTGAATCTGCAAAAAACATGTTTACTATGCTATTCATTATAACAGGCGGATTAATAGCGGCAATACTAAGTTGGAAAATCTCTAAGACCTTTACTAAACACTACCTTAGCTTACTTAAACAGCAAAAACTAGAGGGCCCGCGGTTCGAAGGTAAGCCTATCCCTAACCCTCTCCTCGGTCTCGATTCTACGCGTACCGGTCATCATCACCATCACCATTGAgagctcgcatgcactagt

```
>NanohaloarchFusexin1-V5-6His (pBPT05)EDITED
From 1 to 1872.

ggtaccATGTCTACGTTTTTGATAGTTCCAGCTTCTGTATCTTATGTATCGGAAGTAGACTATATTGAAACAACTTCAGAGTTTTTTGAC 90
       M  S  T  F  L  I  V  P  A  S  V  S  Y  V  S  E  V  D  Y  I  E  T  T  S  E  F  F  D  30

CAGGGCCACGTAGTTATCGAATTTGTTATTGATGGCGCTGATGAGAAAATACATGCAGATCTAGAACCTAGCCAGTTATCAGATTTATCT 180
 Q  G  H  V  V  I  E  F  V  I  D  G  A  D  E  K  I  H  A  D  L  E  P  S  Q  L  S  D  L  S  60

GATCAAAATACGAATCAAAGTGTTGAGATTAGCGCGAAAGTATCCGATGTCTATGCCGAGTATGAAGTAGAAGATAGAGGTCTCAGGCCT 270
 D  Q  N  T  N  Q  S  V  E  I  S  A  K  V  S  D  V  Y  A  E  Y  E  V  E  D  R  G  L  R  P  90

TTAAAACCTATAGAATACACGAGAACTACTCATTCCACAAAAGAAAAAGCTCAGATTTGGGCCGATAATAACTGCCTTGATCTTGATAAT 360
 L  K  P  I  E  Y  T  R  T  T  H  S  T  K  E  K  A  Q  I  W  A  D  N  N  C  L  D  L  D  N  120

AGCGGAGCACCCAACTATAATACTTATAGAAGAGGATTGGTTAACCCCTGGGGAGACTGGATTGTGAATTGTGCGATACCTAATATGAGC 450
 S  G  A  P  N  Y  N  T  Y  R  R  G  L  V  N  P  W  G  D  W  I  V  N  C  A  I  P  N  M  S  150

CAATCTACTTTTAACGTTGGTTTCTTACAGAGTCCTCCAGATACGTTATTTGAGTCTGAATGGATTATAGCTGTTGAAGGTGAATCTGCC 540
 Q  S  T  F  N  V  G  F  L  Q  S  P  P  D  T  L  F  E  S  E  W  I  I  A  V  E  G  E  S  A  180

GAAACAGTTCGATTATCTAATAGTGGCGCAGAACATGGAAGTTTGGAAAGAGTGAATGATAATGTTCAACTCGAGTTTGTTTCATTGAAA 630
 E  T  V  R  L  S  N  S  G  A  E  H  G  S  L  E  R  V  N  D  N  V  Q  L  E  F  V  S  L  K  210

GATACCGGGAGTAATGCTCCTAACCCTCAAAATACTCTAGCTGCTTTTTCAAGCACAGATGGCTGGAGATTAATTGAAAAAGAAGAATTT 720
 D  T  G  S  N  A  P  N  P  Q  N  T  L  A  A  F  S  S  T  D  G  W  R  L  I  E  K  E  E  F  240

GATAGCTACATAACTTATATTGAAAATGAATCTCCTAAAAAAGTTGAAGAGATGCGAGAACAAGATAATTTCAATATAGAAGACTGGACT 810
 D  S  Y  I  T  Y  I  E  N  E  S  P  K  K  V  E  E  M  R  E  Q  D  N  F  N  I  E  D  W  T  270

GATAAGAAAAACAAAGAAGCCAGAGATAACTTTTTGAGCGCATCAAGAGAATACGAAGAATCTCCTATTCAAGGGTCAAACGTATCTGAT 900
 D  K  K  N  K  E  A  R  D  N  F  L  S  A  S  R  E  Y  E  E  S  P  I  Q  G  S  N  V  S  D  300

GGCAAAAACAAATTTCAAAATGGGATTTTTCAATATAGGCCTGAATCAGAGTTCGAATGGTTTATCTCAGAGTTTAACGTTAGATTAAAT 990
 G  K  N  K  F  Q  N  G  I  F  Q  Y  R  P  E  S  E  F  E  W  F  I  S  E  F  N  V  R  L  N  330

GGTGATTTTGTAGGACTACAGCGAAAATTTGCAGATTTAGAGATTATAAATATAGAAGATACCAAAGTTAATGGTCTTTCAAGCGGTTCT 1080
 G  D  F  V  G  L  Q  R  K  F  A  D  L  E  I  I  N  I  E  D  T  K  V  N  G  L  S  S  G  S  360

GTGGAAGTAAAAGTTGCAAATAATGGCAATGGAACCGGCGCTTTTGAAGTTTCAGCTTCCTGTGATAAATTTGATACAGTTGGAGTAAGT 1170
 V  E  V  K  V  A  N  N  G  N  G  T  G  A  F  E  V  S  A  S  C  D  K  F  D  T  V  G  V  S  390

AGTCGCACATCATTAAAATCTGGAGAATCTGACACAGTATTTCTAAATTTGGATACTGACGGTTCTGAAGAGTTCTATGAAACTTGTAAT 1260
 S  R  T  S  L  K  S  G  E  S  D  T  V  F  L  N  L  D  T  D  G  S  E  E  F  Y  E  T  C  N  420

CTCAATATTAAGGACCTTGAAACTACTTCAATCGTTTCTAGCTCATTTACTGCATCTTACACTCCTGAAAAAGAATGTACACCAGGAAGA 1350
 L  N  I  K  D  L  E  T  T  S  I  V  S  S  S  F  T  A  S  Y  T  P  E  K  E  C  T  P  G  R  450

CAGTCAGTCAGAATAGAGAACGGACAAGAAGCAATTTACGAATGTGCTGAAGACGGAATTTCCATATTTGAGGTTGATGCTTGCGAATTG 1440
 Q  S  V  R  I  E  N  G  Q  E  A  I  Y  E  C  A  E  D  G  I  S  I  F  E  V  D  A  C  E  L  480

GACGAAAGAGCTAAACCTTACGGTGAAGGATATGAGTGTCAAAAAGTTGAAAGTCTTCGGGAAATTGAAGCACCTCAAGACTGCGAAATT 1530
 D  E  R  A  K  P  Y  G  E  G  Y  E  C  Q  K  V  E  S  L  R  E  I  E  A  P  Q  D  C  E  I  510

TCCATACTTCCTAGCTTCCTGAATAACTTATTCGCTAATGACTATACCTTTACAAATCCAATTTGCGCCACGGAAAACAGGTTAAATGAT 1620
 S  I  L  P  S  F  L  N  N  L  F  A  N  D  Y  T  F  T  N  P  I  C  A  T  E  N  R  L  N  D  540

TTTAATAATAGTATTGAATCTGCAAAAAACATGTTTACTATGCTATTCATTATAACAGGCGGATTAATAGCGGCAATACTAAGTTGGAAA 1710
 F  N  N  S  I  E  S  A  K  N  M  F  T  M  L  F  I  I  T  G  G  L  I  A  A  I  L  S  W  K  570

ATCTCTAAGACCTTTACTAAACACTACCTTAGCTTACTTAAACAGCAAAAACTAGAGGGCCCGCGGTTCGAAGGTAAGCCTATCCCTAAC 1800
 I  S  K  T  F  T  K  H  Y  L  S  L  L  K  Q  Q  K  L  E  G  P  R  F  E  G  K  P  I  P  N  600

CCTCTCCTCGGTCTCGATTCTACGCGTACCGGTCATCATCACCATCACCATTGAgagctcgcatgcactagt 1872
 P  L  L  G  L  D  S  T  R  T  G  H  H  H  H  H  H  *                    624
```

Synthesized plasmid: pBPT06

3300014206-Ga0172377-10000119-870930-129 (Methanomicrobia)

3300014206Sub.Fusexin1-V5-6His (pBPT06)EDITED

**Addition of restriction Ez. Sites**

**5’ KpnI, 3’ SacI, SphI, SpeI**

ggtaccATGAACAAAATGAGAAGTACGATAAAAATATCAATATTTTTTTTATTGATATTCGCACTAACAATACCAATACAAAATGTTAGTGCAGAAGTAGATGGATTTACATTGAAAGCTGCATCTCTGTCACATCCAGTATCAAACAATAAAGATTTAGAAGATGCAAACTTGACAATTCTTGTTGGAACTGGAGGAGGGCAATCACTACATGGAAAATTTGGAAATGAACAACTAACTCAATGGATTCCAAATTTATTATTCAAGTATCCACTGATATTTGAAATGAGTCCTATAATAGAAACTGCAGACTATCCAATAAGAAATCAAGCACATAGCGAATTAAAAAGACTTGAATATGAAATAAAAGGAAACTGGTTATTTTCAGAAAAATGTAATTCATCAATGTTATATTGTTTCGATGTCGGATCTGGAAAAATCATTGAAATAAAAGAAGTCGATGTTGGAAGATATGGATCAATAGATAATCCAAATTTATTTTGGGGAGGAAATGTAAAAGTAACAATAAATGGGATTCCATACTCAAAAGATATTTCAAGTAACATAGAAAGTGTACAATTTAATTCTATAACAGGAGATTTTGTTGCACTAATAAGATGGGTTGGTTCTACTGTAACAGGACAAGGACTTCCTGATCAACACAATAATGTTCCTATATATAATAAAGACAACGAAAAATGGTCAATATCAAATATTGCAACTTTCAGAGATTACGAACGATCATTACTAACAACTGAAATAGAGTTGAATGCTAAACAAGAACAAAAGAAGGCATTTGATAATAAATTCTATGAATATGCAGCAGAAGAATATTTTGATAAGTATTCGATATGTAAAGATGATCAATGTTCACCGATTGTAAGTATTATAAATACACACAATGCAAAGCTAACAATATTGTTAAATGAACGAAAGAAAATTGGTTATGAATCAGACATATCATCTAATCAACTAACTACAAAAGATGGAAATGTTTTACTTGTATTAAATAGAAGAATAAATATTCCTGAATTTACAATAATAACAACTGCTAAGAGTGTAGGAGCATACATACCAACTGGAGAATTCCAGATAATAGATATAACATCACCAACTTTTAGTTCTGGAGATAGTAATGGAAAAGTTGATATAAAAATAAGAAATAATGGAGATTATAAAGGAACTGCTACTGCAACTTTAATAGATGAAACTGATACATTTAAACAAATATCGAATTCAAAATCGTTAGAAACGACACTCGATCCTGGTAAAGAAGGAACTATGACAATGTACATATCAAGTGGTACAATATCGAGTGATATTACCAAAACTGCTAAGATAGATGTATATGATATAAATAGTCCAAACAATGTAACATCAAAAGAATTCACTGCATCTGTAACAACACCAAAAATGTGTCAACCAGATTCTCAAAGATTAGACAATAAAATTGTATACACATGTAATACAGATGGAACTGCTGAATATGTAACATTAGATTGTTCAAATGGATTACCAAACTATGAAAATAATACATATACATGCACTCAAATCGAATATAATAAAGGTTTGGTGATGACAAATGTCAAAAATCCAATCAAAGTAGTCGAGAAACAAAACAAAACTAAAAAAGAAGATAAAAAAGATGAGTCATCAACTGCATACAATTTTTTAAAATGGATGACAGGTTTGGTGTTATTATACTTTGCAATCGCCTTTATAAGAAAACATAATAACAAAGATCAAATAATCACAAAAGAAAACACAAGATTATCGATAATACAATCGTCCTTAGTGTTAGGAATATTTTATATATATTTACAATTTAAGTTTCTAACAGAAACACTTGGTTCGACATTCTGGATTATAGTTATAATATCTGGAATATTGGTTGGACTTGTTGACATGTATCTAAAAATGACAATACATATGAATATTCCAAAAATATATGTGCTAACAGCAGGTATAATATTATTTATAATTCTATCTCAAATAGCTGTTGGAATAAAAGAAACTGCTTGTAATAGTTGGACAACAAAATGGTTAGCAAATTGGATGTTTGACACATGCAAGGAATGGAGTATAAATGACTATCTACCAGAATGGCTGAAACTAGAGGGCCCGCGGTTCGAAGGTAAGCCTATCCCTAACCCTCTCCTCGGTCTCGATTCTACGCGTACCGGTCATCATCACCATCACCATTGAgagctcgcatgcactagt

```
3300014206Sub.Fusexin1-V5-6His (pBPT06)EDITED

From 1 to 2223.
ggtaccATGAACAAAATGAGAAGTACGATAAAAATATCAATATTTTTTTTATTGATATTCGCACTAACAATACCAATACAAAATGTTAGT 90
       M  N  K  M  R  S  T  I  K  I  S  I  F  F  L  L  I  F  A  L  T  I  P  I  Q  N  V  S  30

GCAGAAGTAGATGGATTTACATTGAAAGCTGCATCTCTGTCACATCCAGTATCAAACAATAAAGATTTAGAAGATGCAAACTTGACAATT 180
 A  E  V  D  G  F  T  L  K  A  A  S  L  S  H  P  V  S  N  N  K  D  L  E  D  A  N  L  T  I  60

CTTGTTGGAACTGGAGGAGGGCAATCACTACATGGAAAATTTGGAAATGAACAACTAACTCAATGGATTCCAAATTTATTATTCAAGTAT 270
 L  V  G  T  G  G  G  Q  S  L  H  G  K  F  G  N  E  Q  L  T  Q  W  I  P  N  L  L  F  K  Y  90

CCACTGATATTTGAAATGAGTCCTATAATAGAAACTGCAGACTATCCAATAAGAAATCAAGCACATAGCGAATTAAAAAGACTTGAATAT 360
 P  L  I  F  E  M  S  P  I  I  E  T  A  D  Y  P  I  R  N  Q  A  H  S  E  L  K  R  L  E  Y  120

GAAATAAAAGGAAACTGGTTATTTTCAGAAAAATGTAATTCATCAATGTTATATTGTTTCGATGTCGGATCTGGAAAAATCATTGAAATA 450
 E  I  K  G  N  W  L  F  S  E  K  C  N  S  S  M  L  Y  C  F  D  V  G  S  G  K  I  I  E  I  150

AAAGAAGTCGATGTTGGAAGATATGGATCAATAGATAATCCAAATTTATTTTGGGGAGGAAATGTAAAAGTAACAATAAATGGGATTCCA 540
 K  E  V  D  V  G  R  Y  G  S  I  D  N  P  N  L  F  W  G  G  N  V  K  V  T  I  N  G  I  P  180

TACTCAAAAGATATTTCAAGTAACATAGAAAGTGTACAATTTAATTCTATAACAGGAGATTTTGTTGCACTAATAAGATGGGTTGGTTCT 630
 Y  S  K  D  I  S  S  N  I  E  S  V  Q  F  N  S  I  T  G  D  F  V  A  L  I  R  W  V  G  S  210

ACTGTAACAGGACAAGGACTTCCTGATCAACACAATAATGTTCCTATATATAATAAAGACAACGAAAAATGGTCAATATCAAATATTGCA 720
 T  V  T  G  Q  G  L  P  D  Q  H  N  N  V  P  I  Y  N  K  D  N  E  K  W  S  I  S  N  I  A  240

ACTTTCAGAGATTACGAACGATCATTACTAACAACTGAAATAGAGTTGAATGCTAAACAAGAACAAAAGAAGGCATTTGATAATAAATTC 810
 T  F  R  D  Y  E  R  S  L  L  T  T  E  I  E  L  N  A  K  Q  E  Q  K  K  A  F  D  N  K  F  270

TATGAATATGCAGCAGAAGAATATTTTGATAAGTATTCGATATGTAAAGATGATCAATGTTCACCGATTGTAAGTATTATAAATACACAC 900
 Y  E  Y  A  A  E  E  Y  F  D  K  Y  S  I  C  K  D  D  Q  C  S  P  I  V  S  I  I  N  T  H  300

AATGCAAAGCTAACAATATTGTTAAATGAACGAAAGAAAATTGGTTATGAATCAGACATATCATCTAATCAACTAACTACAAAAGATGGA 990
 N  A  K  L  T  I  L  L  N  E  R  K  K  I  G  Y  E  S  D  I  S  S  N  Q  L  T  T  K  D  G  330

AATGTTTTACTTGTATTAAATAGAAGAATAAATATTCCTGAATTTACAATAATAACAACTGCTAAGAGTGTAGGAGCATACATACCAACT 1080
 N  V  L  L  V  L  N  R  R  I  N  I  P  E  F  T  I  I  T  T  A  K  S  V  G  A  Y  I  P  T  360

GGAGAATTCCAGATAATAGATATAACATCACCAACTTTTAGTTCTGGAGATAGTAATGGAAAAGTTGATATAAAAATAAGAAATAATGGA 1170
 G  E  F  Q  I  I  D  I  T  S  P  T  F  S  S  G  D  S  N  G  K  V  D  I  K  I  R  N  N  G  390

GATTATAAAGGAACTGCTACTGCAACTTTAATAGATGAAACTGATACATTTAAACAAATATCGAATTCAAAATCGTTAGAAACGACACTC 1260
 D  Y  K  G  T  A  T  A  T  L  I  D  E  T  D  T  F  K  Q  I  S  N  S  K  S  L  E  T  T  L  420

GATCCTGGTAAAGAAGGAACTATGACAATGTACATATCAAGTGGTACAATATCGAGTGATATTACCAAAACTGCTAAGATAGATGTATAT 1350
 D  P  G  K  E  G  T  M  T  M  Y  I  S  S  G  T  I  S  S  D  I  T  K  T  A  K  I  D  V  Y  450

GATATAAATAGTCCAAACAATGTAACATCAAAAGAATTCACTGCATCTGTAACAACACCAAAAATGTGTCAACCAGATTCTCAAAGATTA 1440
 D  I  N  S  P  N  N  V  T  S  K  E  F  T  A  S  V  T  T  P  K  M  C  Q  P  D  S  Q  R  L  480

GACAATAAAATTGTATACACATGTAATACAGATGGAACTGCTGAATATGTAACATTAGATTGTTCAAATGGATTACCAAACTATGAAAAT 1530
 D  N  K  I  V  Y  T  C  N  T  D  G  T  A  E  Y  V  T  L  D  C  S  N  G  L  P  N  Y  E  N  510

AATACATATACATGCACTCAAATCGAATATAATAAAGGTTTGGTGATGACAAATGTCAAAAATCCAATCAAAGTAGTCGAGAAACAAAAC 1620
 N  T  Y  T  C  T  Q  I  E  Y  N  K  G  L  V  M  T  N  V  K  N  P  I  K  V  V  E  K  Q  N  540

AAAACTAAAAAAGAAGATAAAAAAGATGAGTCATCAACTGCATACAATTTTTTAAAATGGATGACAGGTTTGGTGTTATTATACTTTGCA 1710
 K  T  K  K  E  D  K  K  D  E  S  S  T  A  Y  N  F  L  K  W  M  T  G  L  V  L  L  Y  F  A  570

ATCGCCTTTATAAGAAAACATAATAACAAAGATCAAATAATCACAAAAGAAAACACAAGATTATCGATAATACAATCGTCCTTAGTGTTA 1800
 I  A  F  I  R  K  H  N  N  K  D  Q  I  I  T  K  E  N  T  R  L  S  I  I  Q  S  S  L  V  L  600

GGAATATTTTATATATATTTACAATTTAAGTTTCTAACAGAAACACTTGGTTCGACATTCTGGATTATAGTTATAATATCTGGAATATTG 1890
 G  I  F  Y  I  Y  L  Q  F  K  F  L  T  E  T  L  G  S  T  F  W  I  I  V  I  I  S  G  I  L  630

GTTGGACTTGTTGACATGTATCTAAAAATGACAATACATATGAATATTCCAAAAATATATGTGCTAACAGCAGGTATAATATTATTTATA 1980
 V  G  L  V  D  M  Y  L  K  M  T  I  H  M  N  I  P  K  I  Y  V  L  T  A  G  I  I  L  F  I  660

ATTCTATCTCAAATAGCTGTTGGAATAAAAGAAACTGCTTGTAATAGTTGGACAACAAAATGGTTAGCAAATTGGATGTTTGACACATGC 2070
 I  L  S  Q  I  A  V  G  I  K  E  T  A  C  N  S  W  T  T  K  W  L  A  N  W  M  F  D  T  C  690

AAGGAATGGAGTATAAATGACTATCTACCAGAATGGCTGAAACTAGAGGGCCCGCGGTTCGAAGGTAAGCCTATCCCTAACCCTCTCCTC 2160
 K  E  W  S  I  N  D  Y  L  P  E  W  L  K  L  E  G  P  R  F  E  G  K  P  I  P  N  P  L  L  720

GGTCTCGATTCTACGCGTACCGGTCATCATCACCATCACCATTGAgagctcgcatgcactagt 2223
 G  L  D  S  T  R  T  G  H  H  H  H  H  H  *                    741
```

Synthesized plasmid: pBPT07

3300014208-Ga0172379-10000243-871512-158(Methanomicrobia?)

>3300014208Sub.Fusexin1-V5-6His (pBPT07)EDITED

**Addition of restriction Ez. Sites**

**5’ KpnI, 3’ SacI, SphI, SpeI**

ggtaccATGAACCCAATAAAACCCATATTTCTATCAATACTCATAATCGCTATCTTAATTCCCTCCTCTGTCGCCTCCACTACAGCAGGATCAGGATTTACTGTAATGTCAATATCCCCATCTAAAATAATTTCCAACGATGAAGACTTAAAAACTGCCAATTTCATCGTAACAGTCGTAGCAAACGGAAATTCTCAATCTATCATCGGAGAACTCCTCACCGCCAATGACATCAATGCTAAACTCTCCGAATCAGGCTCCTCAGTTCGAGTTTCCTCCCCCTTTAAAATTGAAATGGGTTCAGTCAAAGAAACCCTAACTTACCCCATCCGTAACACAGGCGCAATTCGTAAATACGAATATGCATATTCATCTGAAGTTAATTTTGTTTCGGAAGCAACATGTCTATCAGGCTATTCATATTGTTATACAGTCGAGGCAGGTTCCACATGGTGGGGAAAAACATACAGAAGTCTCAATATTCGCCAAGCAAACAGCGCCGTCTCCTCTTCCCTCGGAGAACTCGATAATCCAAACATTAATTGGCAGGGCGATATAGTTCTAACAGCAGGACTAACTCGTGTATCAAAAACAATAGGTTCAATCGAATCCCTTGGAGCAGCAGATTTCTATAACAGTAACGAGTGGATAGCTCAAGCTACATGGACAGGTTCTTTACTCACAGGGCAAGCAACCCCCAATCCTTCTCTCTATAAAGCAACCTTTACTCCCGCCTTTAATCAATGGAGAATCTCCTCAAATTCCTACTATAATGACTATGCATCCTCCCTATCTGCAAAACAAGCATACTTCCCCTCAATGTTAAGCCAATATACTAAACAAGTGTGTCTTGATTCATACGGTTGTCCATCAGAAACTCCCCTAACAAACGCAATCAGCAATCATAATAGCATTGTAGACATCCTCTTATCTCAAAACACTAAAGTAACATACACCTCCTCTACAGGAGAAACTCTAGCCCAATCAACATCAGGAACTTCCTACTCAGGTAACGTAATCGAGACAACCAATCGTCGTATCTCCAATCCTGTAATCGTGTTCAAAGTCAAAGCATTATCCTTATCTCCCTACATTCCCGTCGGAAAACCACGCATCGAATCAATAACAACACCCGAATTCTCTAGCGGTGACAACAATGGAGTAGCAACAGTTCAATTCAAAAATATCGGTACTGAATCAGCAACATTCTCAGCAACATTCTTTGATCCCTCAGGAACTTTCCACCCCTCCTCTAATCAAGAATCCTCAAAAACTACATGCCCTGCCCTTTCCCAATGCATCATAAACGTCTTTATCTCCAGTGACTCCCCTACCAATGATCGTATCAGTAAAACTGCTACAATGACAGTCTATGACATAAACAAACCTTCCAATTCTGATACAAAAACCTTCGACATAATAATGTCACCTCCTAAAACCTGTATCCCTAGTTCCACTCGCACCTCAGGAAAACTCATCTATACCTGTAACGCCGACGGTTCAGGGGAAACAGTAATAGAATGCACAGGAACAATCTCCCTTCAAAACAGCAAATACATTTGCATAGGTGGTTCAGTAACTCCCAAACCCGGAACAACACCAGTACCGAAAACAATATTCGCAACCCTATCTGACTTCTGGAGTTCACCCCAGCTAATACTAACTGTTCTAGTCCTTCTAATACTATCGTCTCTAATCTACATCAAATACAAAAAACTAGAGGGCCCGCGGTTCGAAGGTAAGCCTATCCCTAACCCTCTCCTCGGTCTCGATTCTACGCGTACCGGTCATCATCACCATCACCATTGAgagctcgcatgcactagt

```
>3300014208Sub.Fusexin1-V5-6His (pBPT07)EDITED
From 1 to 1827.
ggtaccATGAACCCAATAAAACCCATATTTCTATCAATACTCATAATCGCTATCTTAATTCCCTCCTCTGTCGCCTCCACTACAGCAGGA 90
       M  N  P  I  K  P  I  F  L  S  I  L  I  I  A  I  L  I  P  S  S  V  A  S  T  T  A  G  30

TCAGGATTTACTGTAATGTCAATATCCCCATCTAAAATAATTTCCAACGATGAAGACTTAAAAACTGCCAATTTCATCGTAACAGTCGTA 180
 S  G  F  T  V  M  S  I  S  P  S  K  I  I  S  N  D  E  D  L  K  T  A  N  F  I  V  T  V  V  60

GCAAACGGAAATTCTCAATCTATCATCGGAGAACTCCTCACCGCCAATGACATCAATGCTAAACTCTCCGAATCAGGCTCCTCAGTTCGA 270
 A  N  G  N  S  Q  S  I  I  G  E  L  L  T  A  N  D  I  N  A  K  L  S  E  S  G  S  S  V  R  90

GTTTCCTCCCCCTTTAAAATTGAAATGGGTTCAGTCAAAGAAACCCTAACTTACCCCATCCGTAACACAGGCGCAATTCGTAAATACGAA 360
 V  S  S  P  F  K  I  E  M  G  S  V  K  E  T  L  T  Y  P  I  R  N  T  G  A  I  R  K  Y  E  120

TATGCATATTCATCTGAAGTTAATTTTGTTTCGGAAGCAACATGTCTATCAGGCTATTCATATTGTTATACAGTCGAGGCAGGTTCCACA 450
 Y  A  Y  S  S  E  V  N  F  V  S  E  A  T  C  L  S  G  Y  S  Y  C  Y  T  V  E  A  G  S  T  150

TGGTGGGGAAAAACATACAGAAGTCTCAATATTCGCCAAGCAAACAGCGCCGTCTCCTCTTCCCTCGGAGAACTCGATAATCCAAACATT 540
 W  W  G  K  T  Y  R  S  L  N  I  R  Q  A  N  S  A  V  S  S  S  L  G  E  L  D  N  P  N  I  180

AATTGGCAGGGCGATATAGTTCTAACAGCAGGACTAACTCGTGTATCAAAAACAATAGGTTCAATCGAATCCCTTGGAGCAGCAGATTTC 630
 N  W  Q  G  D  I  V  L  T  A  G  L  T  R  V  S  K  T  I  G  S  I  E  S  L  G  A  A  D  F  210

TATAACAGTAACGAGTGGATAGCTCAAGCTACATGGACAGGTTCTTTACTCACAGGGCAAGCAACCCCCAATCCTTCTCTCTATAAAGCA 720
 Y  N  S  N  E  W  I  A  Q  A  T  W  T  G  S  L  L  T  G  Q  A  T  P  N  P  S  L  Y  K  A  240

ACCTTTACTCCCGCCTTTAATCAATGGAGAATCTCCTCAAATTCCTACTATAATGACTATGCATCCTCCCTATCTGCAAAACAAGCATAC 810
 T  F  T  P  A  F  N  Q  W  R  I  S  S  N  S  Y  Y  N  D  Y  A  S  S  L  S  A  K  Q  A  Y  270

TTCCCCTCAATGTTAAGCCAATATACTAAACAAGTGTGTCTTGATTCATACGGTTGTCCATCAGAAACTCCCCTAACAAACGCAATCAGC 900
 F  P  S  M  L  S  Q  Y  T  K  Q  V  C  L  D  S  Y  G  C  P  S  E  T  P  L  T  N  A  I  S  300

AATCATAATAGCATTGTAGACATCCTCTTATCTCAAAACACTAAAGTAACATACACCTCCTCTACAGGAGAAACTCTAGCCCAATCAACA 990
 N  H  N  S  I  V  D  I  L  L  S  Q  N  T  K  V  T  Y  T  S  S  T  G  E  T  L  A  Q  S  T  330

TCAGGAACTTCCTACTCAGGTAACGTAATCGAGACAACCAATCGTCGTATCTCCAATCCTGTAATCGTGTTCAAAGTCAAAGCATTATCC 1080
 S  G  T  S  Y  S  G  N  V  I  E  T  T  N  R  R  I  S  N  P  V  I  V  F  K  V  K  A  L  S  360

TTATCTCCCTACATTCCCGTCGGAAAACCACGCATCGAATCAATAACAACACCCGAATTCTCTAGCGGTGACAACAATGGAGTAGCAACA 1170
 L  S  P  Y  I  P  V  G  K  P  R  I  E  S  I  T  T  P  E  F  S  S  G  D  N  N  G  V  A  T  390

GTTCAATTCAAAAATATCGGTACTGAATCAGCAACATTCTCAGCAACATTCTTTGATCCCTCAGGAACTTTCCACCCCTCCTCTAATCAA 1260
 V  Q  F  K  N  I  G  T  E  S  A  T  F  S  A  T  F  F  D  P  S  G  T  F  H  P  S  S  N  Q  420

GAATCCTCAAAAACTACATGCCCTGCCCTTTCCCAATGCATCATAAACGTCTTTATCTCCAGTGACTCCCCTACCAATGATCGTATCAGT 1350
 E  S  S  K  T  T  C  P  A  L  S  Q  C  I  I  N  V  F  I  S  S  D  S  P  T  N  D  R  I  S  450

AAAACTGCTACAATGACAGTCTATGACATAAACAAACCTTCCAATTCTGATACAAAAACCTTCGACATAATAATGTCACCTCCTAAAACC 1440
 K  T  A  T  M  T  V  Y  D  I  N  K  P  S  N  S  D  T  K  T  F  D  I  I  M  S  P  P  K  T  480

TGTATCCCTAGTTCCACTCGCACCTCAGGAAAACTCATCTATACCTGTAACGCCGACGGTTCAGGGGAAACAGTAATAGAATGCACAGGA 1530
 C  I  P  S  S  T  R  T  S  G  K  L  I  Y  T  C  N  A  D  G  S  G  E  T  V  I  E  C  T  G  510

ACAATCTCCCTTCAAAACAGCAAATACATTTGCATAGGTGGTTCAGTAACTCCCAAACCCGGAACAACACCAGTACCGAAAACAATATTC 1620
 T  I  S  L  Q  N  S  K  Y  I  C  I  G  G  S  V  T  P  K  P  G  T  T  P  V  P  K  T  I  F  540

GCAACCCTATCTGACTTCTGGAGTTCACCCCAGCTAATACTAACTGTTCTAGTCCTTCTAATACTATCGTCTCTAATCTACATCAAATAC 1710
 A  T  L  S  D  F  W  S  S  P  Q  L  I  L  T  V  L  V  L  L  I  L  S  S  L  I  Y  I  K  Y  570

AAAAAACTAGAGGGCCCGCGGTTCGAAGGTAAGCCTATCCCTAACCCTCTCCTCGGTCTCGATTCTACGCGTACCGGTCATCATCACCAT 1800
 K  K  L  E  G  P  R  F  E  G  K  P  I  P  N  P  L  L  G  L  D  S  T  R  T  G  H  H  H  H  600

CACCATTGAgagctcgcatgcactagt 1827
 H  H  *                    609
```

Synthesized plasmid: pBPT08

3300014208-Ga0172379-10001592-871560-40 (Methanomicrobia)

>3300014208Sub.Fusexin1-V5-6His (pBPT08)EDITED

**Addition of restriction Ez. Sites**

**5’ KpnI, 3’ SacI, SphI, SpeI**

ggtaccATGATGAACAAAATGAGAAGCACGATAAAAATATCAATATTTTTTTTATTGATATTCACACTAACAATACCAACTGTTAGTGCAGAAACAGACGGATTTACATTAAAAGCTGCAACAATATCACATCCAGTATCAAACAATAAAGATTTAGAAGATGCAAACATGACACTTCTTGTTGGATCTGGCGGAGGACAATCTTTATATGGAAAATTTGGAAATGAACAATTTACACAATGGATTCCAAATCTATTGTTTAAATATCCACTTGTGTTTGAACTTGATTCGATAAAAGAAACTGCAGATTATCAAATCAGAAATCAAGGAGACTTAAAGAAACTTGAATATACTATAAGAGGAAACTGGTTCTATTCTGCTAAATGCAATCCAGATATACCATATTGTTTTGATGTCGGTTCTGGAAAAATAATTGAAGTAAGATACGTTGACGTTGGAAGATATGGTTCACTCGACACGCCAAATTTAAATTTTAAAGGAAAAGCAACAGTAACAATAAATGGTGTTCCTTATACAAAAGAAATTTCAAATATAATAAGTAGCATTCAGTTCAATTCAATAACAGGAGAATTTGTTGCTTTAATAAAATGGGATGGTTCTTCTGTAACAGGGCAAGGATTACCAGATCAACATAATTACGTTCCAGTGAATAAAAAAGATAGCGGAGAATGGACTCTGTCAACATTATCTTTATATAAAGAATATGATGATTCACTTCTTAAAACAGAAATAGAATTAAACAACAAAAAAGAACAAAAGAGAAAGTTTGATAATAAATTCTATGAATATGCTGCAGAAAAATATGTAGATAAATACTCTATATGTAAAGATAACGAATGTTCACCAATTGTAAACATTGTAAATGAACATAATGCAAAATTAACAAAATTGTTAGATTCAAGAGAAAAAATTGGTTATGAATCATCTATATCATCCGTACAAAATTCATATAAAGATGGAAATGTTTTAGTTGTCTTAAACAGAAGAATAAACATTCCTGAATTTACAATAATAACAACTGCTAAAAGTGTAGGAGCATACATCCCTGTCGGTGAATTTCAAATAATCGATATAACATCACCAACATTCAGTTCAGGAGATAATAATGGAAAGGTTGATGTAAAAGTAAGAAACATTGGAAACTATAGAGGAACATTCTCATTAACTTTAATAGATGAGACTAACACATTTAAACAAATATCGAATTCAAAATCGCTCGAAACATCACTCGAACCTGGAAAAGAAACAATAATCACAATGTATATATCAAGTGGTATAATATCAAGCGATATTACAAAAACTGCAAAGATAGAAGTTTATGACATAAATAATCCGAATAATTTAACATCGAAAGACTTTACTGTATCAGTAACAACACCAAAAATATGTCAACCTGATTCACAAAGATTAGATAACAAAAAGGTGTATACATGTTCTAATGATGGTATGAGAGAATATCTAACATTAGATTGTTCAAATGGAATTATAGACTATAATAACAATACTTATGATTGTACACAAATAGAATACAATAAAGATTTGGTTATAACAAATGCTGAAACACCAATCAAAATAGTCGAAGTACAAAACAAAACAAAACCAGAAGAACCTAAAGAAGAAACACCGAGTTTAATATACTGGCTAGTTGGATTTGTTATACTTAATATATATATAGCATATATAAGAAAATACAAAAATAAAGATGAAACGATAACTTCTGAAAATATACAAAAATCAGTAATTCTATCTGGACTAATAATAGGAACACTCTATTTGTTAATACAATTTACTTATCTGATAGAAACACTAGGTAGTATAATCTGGTTGATATTTATAATGGCTGTAGTAATAATAGGACTTATTAATGTATATTTAAAAGTAATATATCACATGAACATACCAAAAATATATATTATAACAGCAAGTATAATATTGTTTATAATCCTATCAAATATAGCAATTGGAATAAAAGAAACCGCTTGTGATAGTTGGTCAACGAGATGGTTAGCAAGTAAACTATTTGATACTTGTAAAGAATGGACTATAACAGATTACCTACCAGAATGGTTGAAACTAGAGGGCCCGCGGTTCGAAGGTAAGCCTATCCCTAACCCTCTCCTCGGTCTCGATTCTACGCGTACCGGTCATCATCACCATCACCATTGAgagctcgcatgcactagt

```
>3300014208Sub.Fusexin1-V5-6His (pBPT08)EDITED
From 1 to 2205.
ggtaccATGATGAACAAAATGAGAAGCACGATAAAAATATCAATATTTTTTTTATTGATATTCACACTAACAATACCAACTGTTAGTGCA 90
       M  M  N  K  M  R  S  T  I  K  I  S  I  F  F  L  L  I  F  T  L  T  I  P  T  V  S  A  30

GAAACAGACGGATTTACATTAAAAGCTGCAACAATATCACATCCAGTATCAAACAATAAAGATTTAGAAGATGCAAACATGACACTTCTT 180
 E  T  D  G  F  T  L  K  A  A  T  I  S  H  P  V  S  N  N  K  D  L  E  D  A  N  M  T  L  L  60

GTTGGATCTGGCGGAGGACAATCTTTATATGGAAAATTTGGAAATGAACAATTTACACAATGGATTCCAAATCTATTGTTTAAATATCCA 270
 V  G  S  G  G  G  Q  S  L  Y  G  K  F  G  N  E  Q  F  T  Q  W  I  P  N  L  L  F  K  Y  P  90

CTTGTGTTTGAACTTGATTCGATAAAAGAAACTGCAGATTATCAAATCAGAAATCAAGGAGACTTAAAGAAACTTGAATATACTATAAGA 360
 L  V  F  E  L  D  S  I  K  E  T  A  D  Y  Q  I  R  N  Q  G  D  L  K  K  L  E  Y  T  I  R  120

GGAAACTGGTTCTATTCTGCTAAATGCAATCCAGATATACCATATTGTTTTGATGTCGGTTCTGGAAAAATAATTGAAGTAAGATACGTT 450
 G  N  W  F  Y  S  A  K  C  N  P  D  I  P  Y  C  F  D  V  G  S  G  K  I  I  E  V  R  Y  V  150

GACGTTGGAAGATATGGTTCACTCGACACGCCAAATTTAAATTTTAAAGGAAAAGCAACAGTAACAATAAATGGTGTTCCTTATACAAAA 540
 D  V  G  R  Y  G  S  L  D  T  P  N  L  N  F  K  G  K  A  T  V  T  I  N  G  V  P  Y  T  K  180

GAAATTTCAAATATAATAAGTAGCATTCAGTTCAATTCAATAACAGGAGAATTTGTTGCTTTAATAAAATGGGATGGTTCTTCTGTAACA 630
 E  I  S  N  I  I  S  S  I  Q  F  N  S  I  T  G  E  F  V  A  L  I  K  W  D  G  S  S  V  T  210

GGGCAAGGATTACCAGATCAACATAATTACGTTCCAGTGAATAAAAAAGATAGCGGAGAATGGACTCTGTCAACATTATCTTTATATAAA 720
 G  Q  G  L  P  D  Q  H  N  Y  V  P  V  N  K  K  D  S  G  E  W  T  L  S  T  L  S  L  Y  K  240

GAATATGATGATTCACTTCTTAAAACAGAAATAGAATTAAACAACAAAAAAGAACAAAAGAGAAAGTTTGATAATAAATTCTATGAATAT 810
 E  Y  D  D  S  L  L  K  T  E  I  E  L  N  N  K  K  E  Q  K  R  K  F  D  N  K  F  Y  E  Y  270

GCTGCAGAAAAATATGTAGATAAATACTCTATATGTAAAGATAACGAATGTTCACCAATTGTAAACATTGTAAATGAACATAATGCAAAA 900
 A  A  E  K  Y  V  D  K  Y  S  I  C  K  D  N  E  C  S  P  I  V  N  I  V  N  E  H  N  A  K  300

TTAACAAAATTGTTAGATTCAAGAGAAAAAATTGGTTATGAATCATCTATATCATCCGTACAAAATTCATATAAAGATGGAAATGTTTTA 990
 L  T  K  L  L  D  S  R  E  K  I  G  Y  E  S  S  I  S  S  V  Q  N  S  Y  K  D  G  N  V  L  330

GTTGTCTTAAACAGAAGAATAAACATTCCTGAATTTACAATAATAACAACTGCTAAAAGTGTAGGAGCATACATCCCTGTCGGTGAATTT 1080
 V  V  L  N  R  R  I  N  I  P  E  F  T  I  I  T  T  A  K  S  V  G  A  Y  I  P  V  G  E  F  360

CAAATAATCGATATAACATCACCAACATTCAGTTCAGGAGATAATAATGGAAAGGTTGATGTAAAAGTAAGAAACATTGGAAACTATAGA 1170
 Q  I  I  D  I  T  S  P  T  F  S  S  G  D  N  N  G  K  V  D  V  K  V  R  N  I  G  N  Y  R  390

GGAACATTCTCATTAACTTTAATAGATGAGACTAACACATTTAAACAAATATCGAATTCAAAATCGCTCGAAACATCACTCGAACCTGGA 1260
 G  T  F  S  L  T  L  I  D  E  T  N  T  F  K  Q  I  S  N  S  K  S  L  E  T  S  L  E  P  G  420

AAAGAAACAATAATCACAATGTATATATCAAGTGGTATAATATCAAGCGATATTACAAAAACTGCAAAGATAGAAGTTTATGACATAAAT 1350
 K  E  T  I  I  T  M  Y  I  S  S  G  I  I  S  S  D  I  T  K  T  A  K  I  E  V  Y  D  I  N  450

AATCCGAATAATTTAACATCGAAAGACTTTACTGTATCAGTAACAACACCAAAAATATGTCAACCTGATTCACAAAGATTAGATAACAAA 1440
 N  P  N  N  L  T  S  K  D  F  T  V  S  V  T  T  P  K  I  C  Q  P  D  S  Q  R  L  D  N  K  480

AAGGTGTATACATGTTCTAATGATGGTATGAGAGAATATCTAACATTAGATTGTTCAAATGGAATTATAGACTATAATAACAATACTTAT 1530
 K  V  Y  T  C  S  N  D  G  M  R  E  Y  L  T  L  D  C  S  N  G  I  I  D  Y  N  N  N  T  Y  510

GATTGTACACAAATAGAATACAATAAAGATTTGGTTATAACAAATGCTGAAACACCAATCAAAATAGTCGAAGTACAAAACAAAACAAAA 1620
 D  C  T  Q  I  E  Y  N  K  D  L  V  I  T  N  A  E  T  P  I  K  I  V  E  V  Q  N  K  T  K  540

CCAGAAGAACCTAAAGAAGAAACACCGAGTTTAATATACTGGCTAGTTGGATTTGTTATACTTAATATATATATAGCATATATAAGAAAA 1710
 P  E  E  P  K  E  E  T  P  S  L  I  Y  W  L  V  G  F  V  I  L  N  I  Y  I  A  Y  I  R  K  570

TACAAAAATAAAGATGAAACGATAACTTCTGAAAATATACAAAAATCAGTAATTCTATCTGGACTAATAATAGGAACACTCTATTTGTTA 1800
 Y  K  N  K  D  E  T  I  T  S  E  N  I  Q  K  S  V  I  L  S  G  L  I  I  G  T  L  Y  L  L  600

ATACAATTTACTTATCTGATAGAAACACTAGGTAGTATAATCTGGTTGATATTTATAATGGCTGTAGTAATAATAGGACTTATTAATGTA 1890
 I  Q  F  T  Y  L  I  E  T  L  G  S  I  I  W  L  I  F  I  M  A  V  V  I  I  G  L  I  N  V  630

TATTTAAAAGTAATATATCACATGAACATACCAAAAATATATATTATAACAGCAAGTATAATATTGTTTATAATCCTATCAAATATAGCA 1980
 Y  L  K  V  I  Y  H  M  N  I  P  K  I  Y  I  I  T  A  S  I  I  L  F  I  I  L  S  N  I  A  660

ATTGGAATAAAAGAAACCGCTTGTGATAGTTGGTCAACGAGATGGTTAGCAAGTAAACTATTTGATACTTGTAAAGAATGGACTATAACA 2070
 I  G  I  K  E  T  A  C  D  S  W  S  T  R  W  L  A  S  K  L  F  D  T  C  K  E  W  T  I  T  690

GATTACCTACCAGAATGGTTGAAACTAGAGGGCCCGCGGTTCGAAGGTAAGCCTATCCCTAACCCTCTCCTCGGTCTCGATTCTACGCGT 2160
 D  Y  L  P  E  W  L  K  L  E  G  P  R  F  E  G  K  P  I  P  N  P  L  L  G  L  D  S  T  R  720

ACCGGTCATCATCACCATCACCATTGAgagctcgcatgcactagt 2205
 T  G  H  H  H  H  H  H  *                    735
```

Synthesized plasmid: pBPT10

3300018015-Ga0187866_1000629|915963_9

3300018015Sub.Fusexin1-V5-6His (pBPT10) EDITED

**Addition of restriction Ez. Sites**

**5’ KpnI, 3’ SacI, SphI, SpeI**

SpeI edited ATACTAGT to ATTCTAGT both encoding for Ileu

ggtaccATGAGAAAAAAGAATAGGAACATAATAGCAGGAGTTATtCTAGTTGTTGCTATCTTGATTATTCTTGTTTCCTTCGGAGCATTCAAGCTTCCTTTTTCTGTAATCAATTCTGGATTTTCAACATTATCATTAAGCAGTATAAATGTAAATTCAAATGACCCTGTCCTCGGAGGGCAGACTTGGTTTCTTACAGTTTCTCAAAATGGAGCTTCACAATCAGCGTATGTTGATAATTCTGCTGTCTTAAAAGATGACACTACAGGTGCAACATCAAATCCAATGACAATTTCATTAACCCTTAATAAGAATTATGCTTCTTACAATATATTCAACCAAGGAGTTCCTATAAGGCATCTGGAATACACCACAACATCTTACACTGCAATAAATCTTTTGACTGGAACAAGCGGATGCAACACTGCAGTCTGGAACAATTATCAAATATTTCAGCAAGTATCTTATTGCTTTAATTATGTGACAGATGGATATTATGGAACAATAGGACAAGCAAATCAAGTATTTCAGTCAACAGTAAATGTTCAAGGTTCAGGTGGCTCTGATTCCTGCACAATTTCAAACACAGGAGCAAATACATGCTATTCCCAAGCTGGAAATTTATATGCTTCATGGGTTGGAAGCCTTGTTTCTGGGGCATCTGCTCCTGACCCAAGTTCTCAAGGAATAGTTGCAGTTTATAATACTGCTACAGGAGGATGGAAGACAGCAAATCAAAACTCATATCAGACATATTCACAGCAGAACATAAATTCCTGCATAACTTCTGGAGGAACTAATTGCTTTAATCAGTATAATAATTATGAAAGTGCATTTATGAATGGTTATTCTTTCACATCCACTGGAGGAAATGTTGCATACACGCAAGGAACACAAAATAGTGGGCAGGTTATTTTAAATCTTCCACAGCAATTCAACTTCCCTGTAATTACAATGGGGATAAAGGCAAGTATGGTTGGAATAAATATTCTTTCTGGAAAACCAAAAATATTGTCTGCAACCCCGCAAGTATTCCAGACAGGACAAACAGGAACAATTCAAGCCACTGTTGAAAATGTTGGAACAGCAACAGGAAGCTTTAATGTGGGAGCAACATGCACAAATGGATTTTCTCAGACAGGAAATTCATTGACAATTTCTTCTCTTGCCCCAAATTCCCAGAACACAGTTTATATTCCTATAACCGCAAATGTTGTAAGCGGAACAAATGAAGGAACATGCACAGTAACAGCTACTGACATAAGCAATTCATTGAATTCAGACACTAAGACAGTTTCAGTAAGTGCAAATGCAATAATAATCTGCACAAACGGTCAAATATCAACTAATGGCAATTTGATACAGCAATGCACAAACAATGTTTGGGTAACTATCAAATCATGTAATCCAGCAAATGAAACTGCTCAATTGACAGGCAGTGGAGCAAGTTGTGTTCCTAAAAACTCACCAGGAACTTGCGGATTCTTAGGAGTAGGATGCTTGTGGAATTCAATATTTGGAAGCGGAGGATTCTTTGATAACTTATTTTCAGGAATAGGAAATTTCTTCAGCATAGTAAAATTAATTGCAAGTATTTTGGCAGGTATATTTGCATTCGTCTTTAGCTTAGATTTGTTTAACAAGATAAGTTTTGTAAAAAATCATAAATGGTTGCTTTGGATTATATCATTGATAATTGGATTGCTTTTCTTCTGGCTTGTAACAACAGTATTTTGGATAGGAGTAATTTTATTTGTAGTATATCTTATATTTAAATCAATTATTGGAGCTACTCCAATTGGACAAGCATTTAAATTATTAAAAAAACTAGAGGGCCCGCGGTTCGAAGGTAAGCCTATCCCTAACCCTCTCCTCGGTCTCGATTCTACGCGTACCGGTCATCATCACCATCACCATTGAgagctcgcatgcactagt

3300018015Sub.Fusexin1-V5-6His (pBPT10) EDITED

```
From 1 to 1938.
ggtaccATGAGAAAAAAGAATAGGAACATAATAGCAGGAGTTATtCTAGTTGTTGCTATCTTGATTATTCTTGTTTCCTTCGGAGCATTC 90
       M  R  K  K  N  R  N  I  I  A  G  V  I  L  V  V  A  I  L  I  I  L  V  S  F  G  A  F  30

AAGCTTCCTTTTTCTGTAATCAATTCTGGATTTTCAACATTATCATTAAGCAGTATAAATGTAAATTCAAATGACCCTGTCCTCGGAGGG 180
 K  L  P  F  S  V  I  N  S  G  F  S  T  L  S  L  S  S  I  N  V  N  S  N  D  P  V  L  G  G  60

CAGACTTGGTTTCTTACAGTTTCTCAAAATGGAGCTTCACAATCAGCGTATGTTGATAATTCTGCTGTCTTAAAAGATGACACTACAGGT 270
 Q  T  W  F  L  T  V  S  Q  N  G  A  S  Q  S  A  Y  V  D  N  S  A  V  L  K  D  D  T  T  G  90

GCAACATCAAATCCAATGACAATTTCATTAACCCTTAATAAGAATTATGCTTCTTACAATATATTCAACCAAGGAGTTCCTATAAGGCAT 360
 A  T  S  N  P  M  T  I  S  L  T  L  N  K  N  Y  A  S  Y  N  I  F  N  Q  G  V  P  I  R  H  120

CTGGAATACACCACAACATCTTACACTGCAATAAATCTTTTGACTGGAACAAGCGGATGCAACACTGCAGTCTGGAACAATTATCAAATA 450
 L  E  Y  T  T  T  S  Y  T  A  I  N  L  L  T  G  T  S  G  C  N  T  A  V  W  N  N  Y  Q  I  150

TTTCAGCAAGTATCTTATTGCTTTAATTATGTGACAGATGGATATTATGGAACAATAGGACAAGCAAATCAAGTATTTCAGTCAACAGTA 540
 F  Q  Q  V  S  Y  C  F  N  Y  V  T  D  G  Y  Y  G  T  I  G  Q  A  N  Q  V  F  Q  S  T  V  180

AATGTTCAAGGTTCAGGTGGCTCTGATTCCTGCACAATTTCAAACACAGGAGCAAATACATGCTATTCCCAAGCTGGAAATTTATATGCT 630
 N  V  Q  G  S  G  G  S  D  S  C  T  I  S  N  T  G  A  N  T  C  Y  S  Q  A  G  N  L  Y  A  210

TCATGGGTTGGAAGCCTTGTTTCTGGGGCATCTGCTCCTGACCCAAGTTCTCAAGGAATAGTTGCAGTTTATAATACTGCTACAGGAGGA 720
 S  W  V  G  S  L  V  S  G  A  S  A  P  D  P  S  S  Q  G  I  V  A  V  Y  N  T  A  T  G  G  240

TGGAAGACAGCAAATCAAAACTCATATCAGACATATTCACAGCAGAACATAAATTCCTGCATAACTTCTGGAGGAACTAATTGCTTTAAT 810
 W  K  T  A  N  Q  N  S  Y  Q  T  Y  S  Q  Q  N  I  N  S  C  I  T  S  G  G  T  N  C  F  N  270

CAGTATAATAATTATGAAAGTGCATTTATGAATGGTTATTCTTTCACATCCACTGGAGGAAATGTTGCATACACGCAAGGAACACAAAAT 900
 Q  Y  N  N  Y  E  S  A  F  M  N  G  Y  S  F  T  S  T  G  G  N  V  A  Y  T  Q  G  T  Q  N  300

AGTGGGCAGGTTATTTTAAATCTTCCACAGCAATTCAACTTCCCTGTAATTACAATGGGGATAAAGGCAAGTATGGTTGGAATAAATATT 990
 S  G  Q  V  I  L  N  L  P  Q  Q  F  N  F  P  V  I  T  M  G  I  K  A  S  M  V  G  I  N  I  330

CTTTCTGGAAAACCAAAAATATTGTCTGCAACCCCGCAAGTATTCCAGACAGGACAAACAGGAACAATTCAAGCCACTGTTGAAAATGTT 1080
 L  S  G  K  P  K  I  L  S  A  T  P  Q  V  F  Q  T  G  Q  T  G  T  I  Q  A  T  V  E  N  V  360

GGAACAGCAACAGGAAGCTTTAATGTGGGAGCAACATGCACAAATGGATTTTCTCAGACAGGAAATTCATTGACAATTTCTTCTCTTGCC 1170
 G  T  A  T  G  S  F  N  V  G  A  T  C  T  N  G  F  S  Q  T  G  N  S  L  T  I  S  S  L  A  390

CCAAATTCCCAGAACACAGTTTATATTCCTATAACCGCAAATGTTGTAAGCGGAACAAATGAAGGAACATGCACAGTAACAGCTACTGAC 1260
 P  N  S  Q  N  T  V  Y  I  P  I  T  A  N  V  V  S  G  T  N  E  G  T  C  T  V  T  A  T  D  420

ATAAGCAATTCATTGAATTCAGACACTAAGACAGTTTCAGTAAGTGCAAATGCAATAATAATCTGCACAAACGGTCAAATATCAACTAAT 1350
 I  S  N  S  L  N  S  D  T  K  T  V  S  V  S  A  N  A  I  I  I  C  T  N  G  Q  I  S  T  N  450

GGCAATTTGATACAGCAATGCACAAACAATGTTTGGGTAACTATCAAATCATGTAATCCAGCAAATGAAACTGCTCAATTGACAGGCAGT 1440
 G  N  L  I  Q  Q  C  T  N  N  V  W  V  T  I  K  S  C  N  P  A  N  E  T  A  Q  L  T  G  S  480

GGAGCAAGTTGTGTTCCTAAAAACTCACCAGGAACTTGCGGATTCTTAGGAGTAGGATGCTTGTGGAATTCAATATTTGGAAGCGGAGGA 1530
 G  A  S  C  V  P  K  N  S  P  G  T  C  G  F  L  G  V  G  C  L  W  N  S  I  F  G  S  G  G  510

TTCTTTGATAACTTATTTTCAGGAATAGGAAATTTCTTCAGCATAGTAAAATTAATTGCAAGTATTTTGGCAGGTATATTTGCATTCGTC 1620
 F  F  D  N  L  F  S  G  I  G  N  F  F  S  I  V  K  L  I  A  S  I  L  A  G  I  F  A  F  V  540

TTTAGCTTAGATTTGTTTAACAAGATAAGTTTTGTAAAAAATCATAAATGGTTGCTTTGGATTATATCATTGATAATTGGATTGCTTTTC 1710
 F  S  L  D  L  F  N  K  I  S  F  V  K  N  H  K  W  L  L  W  I  I  S  L  I  I  G  L  L  F  570

TTCTGGCTTGTAACAACAGTATTTTGGATAGGAGTAATTTTATTTGTAGTATATCTTATATTTAAATCAATTATTGGAGCTACTCCAATT 1800
 F  W  L  V  T  T  V  F  W  I  G  V  I  L  F  V  V  Y  L  I  F  K  S  I  I  G  A  T  P  I  600

GGACAAGCATTTAAATTATTAAAAAAACTAGAGGGCCCGCGGTTCGAAGGTAAGCCTATCCCTAACCCTCTCCTCGGTCTCGATTCTACG 1890
 G  Q  A  F  K  L  L  K  K  L  E  G  P  R  F  E  G  K  P  I  P  N  P  L  L  G  L  D  S  T  630

CGTACCGGTCATCATCACCATCACCATTGAgagctcgcatgcactagt 1938
 R  T  G  H  H  H  H  H  H  *                    646
```

Synthesized plasmid: pBPT11

3300000868-JGI12330J12834-1000008-299010-8 (Halobacteria), **(Fsx1 characterized)**

3300000868- Sub.Fusexin1-V5-6His (pBPT11)EDITED

**Addition of restriction Ez. Sites**

**5’ KpnI, 3’ SacI, SphI, SpeI**

SacI GAGCTC to GAGCCC both encoding for Ala

Start codon GTG to ATG

ggtaccATGAGACGTGCAGCATTGATTCTGGCTTTCGTACTGTTCATCGGTCTTTCCTCGGCTACTGTTACCTCGGCTGATTCAATCACGTATAACTCTGGAACCAGCGAGTTCTTCGACGGAGATGTCTTTGCAATAGAGGTAACCGCTGACCAGTCGACTGATGAGATAGATATCTACCTCGGTGCTTCTGAGCTTTCGGAAAAGACGGACGGCGAGGTCAACCAGGACCTCAGTATTGAGTTCACGCACCAGGATTCCAAGCTGAAGTATTCTACTTCGACTTCTGACGAGCTACGGGACATTGTCACTCTTACAACTTACTACGAGGACGGGTTTGACACGGAGCAGGATGCTATAGACGCCATCAAATCAGACTGTTACGATCTGAATCAGAACGGGAATGGAAGCGGAAGGTACAGCAGGTACTACTCAGTTACATCTCCTGTCTATGATTACGAGATCTATTGTTTCCAGAAGAACGAGAAACTGGCTACTCCAGCATACATAGACAACCCGGATGAGATCTTCACTGCAAAGGCTGAGCTACAGGCCGGTGACAAGACGATTCAGTCAGCTACATTGTCTAACGGTGACGCCGGAGACGGAACAGTCACTGACCTCGGTGACTCAAAGATTAGCTGGAACGGGAACCTGGATTTAGGAGCTTCTGAACCGGAGAATAGCCGAGTCATTGCTTTGTACTCGAATGATTTTGAGAACGGGTGGAGGATCGGCAATAAGCAGTCGTACGAGGACTACAAGACGTTCATCGGTGGCGGGGATGCCTACGATCTCCTGATTGACTGGCAGGACGGGACTTACACCGCCTCTGAAGTAGAGGATGAGCTGGTGAATACTGATGCGAACCAGGCTGTTGAGGAGGCCAGTTCTAGCACGACAGACTTGGTGAACGCGAAGGTTAAGGACTCGTCTCTTGACACCGGTTCGTTCGTGTATGATACCCCTGAGCTGCTTTCCTACCCGTCCTTCACAGTCTACGTCGATGCCGGAGAAAACGGATACATCGAGGTAACGAAGCCGACTGGTGACCCCGATATCATCTCCACTTCATCTACAGAAATTAAGGAGGGAGATGAGGGAACTGTAACCGCCACCGTTGAGAATGTCGGAGATGGTGAGGGCGAGTTCTCTGGAAGGCTTTCCAGCTGCGGTGAAGGCTTCAGTATTGTCGACGACCAGAATACGAAGAATGTGGGTGCCGGAGAATCGGTGACATACAGTTTTGATGTCGCTTTCAGTTCTGTCAGCAGTGAGTCGAAGGAGATCTCGGGCAGCTGCACGTTCGAGGTGAACGGCGTCGAGTCCAGTGATTCCACCAGTGTTTCTGTGACCGGTATCCAGCAGAGTGAGTGTAATCCAGGCGATCAGAGACGGGAAAAGAATGAGAATGACCGCTGGGAGATCTACACCTGCCAGGACAACGGCCTGACCTACGAATATGATGTAACTTGTGCAGAGGACGAGAAGGCTGTAGCTCAGGGAGATAACCAGTTTTCCTGTGAGAAACAGGACGATGATTCAGGTGGCGGAGACAACACTGGATCTGACTCCGGACTGTTCTCGAATCTGTTCGGAGGTTCTGGTTCCGGAGATCTGCTTACACAGGTTCACACAGCTCTGTCTATCCTCGCGGGGCTTGTTGCAGGCTTCTTCGGTTATAGAGGAGC**c**CGATGGATTCACGGCGAGACTGATATAAAAGGAGGCTTCAAGCTGGAGTCCCGGAACGTAAGCCGTGTTAAGAGAGGTAGTCCTGTAGCCGGTATTGTCGGAGCTGTACTCGGGTTTGTCGTAGGATACGGTGTTGCCTCAGTCTTCCATCCAGTAGTTCAGATTATAGTGGTTCTGGGAATTGCTGTTGGCTTGTACTATTTCAGGCTAGAGGGCCCGCGGTTCGAAGGTAAGCCTATCCCTAACCCTCTCCTCGGTCTCGATTCTACGCGTACCGGTCATCATCACCATCACCATTGAgagctcgcatgcactagt

```
3300000868- Sub.Fusexin1-V5-6His (pBPT11)EDITED
From 1 to 2010.
ggtaccaTGAGACGTGCAGCATTGATTCTGGCTTTCGTACTGTTCATCGGTCTTTCCTCGGCTACTGTTACCTCGGCTGATTCAATCACG 90
       M  R  R  A  A  L  I  L  A  F  V  L  F  I  G  L  S  S  A  T  V  T  S  A  D  S  I  T  30

TATAACTCTGGAACCAGCGAGTTCTTCGACGGAGATGTCTTTGCAATAGAGGTAACCGCTGACCAGTCGACTGATGAGATAGATATCTAC 180
 Y  N  S  G  T  S  E  F  F  D  G  D  V  F  A  I  E  V  T  A  D  Q  S  T  D  E  I  D  I  Y  60

CTCGGTGCTTCTGAGCTTTCGGAAAAGACGGACGGCGAGGTCAACCAGGACCTCAGTATTGAGTTCACGCACCAGGATTCCAAGCTGAAG 270
 L  G  A  S  E  L  S  E  K  T  D  G  E  V  N  Q  D  L  S  I  E  F  T  H  Q  D  S  K  L  K  90

TATTCTACTTCGACTTCTGACGAGCTACGGGACATTGTCACTCTTACAACTTACTACGAGGACGGGTTTGACACGGAGCAGGATGCTATA 360
 Y  S  T  S  T  S  D  E  L  R  D  I  V  T  L  T  T  Y  Y  E  D  G  F  D  T  E  Q  D  A  I  120

GACGCCATCAAATCAGACTGTTACGATCTGAATCAGAACGGGAATGGAAGCGGAAGGTACAGCAGGTACTACTCAGTTACATCTCCTGTC 450
 D  A  I  K  S  D  C  Y  D  L  N  Q  N  G  N  G  S  G  R  Y  S  R  Y  Y  S  V  T  S  P  V  150

TATGATTACGAGATCTATTGTTTCCAGAAGAACGAGAAACTGGCTACTCCAGCATACATAGACAACCCGGATGAGATCTTCACTGCAAAG 540
 Y  D  Y  E  I  Y  C  F  Q  K  N  E  K  L  A  T  P  A  Y  I  D  N  P  D  E  I  F  T  A  K  180

GCTGAGCTACAGGCCGGTGACAAGACGATTCAGTCAGCTACATTGTCTAACGGTGACGCCGGAGACGGAACAGTCACTGACCTCGGTGAC 630
 A  E  L  Q  A  G  D  K  T  I  Q  S  A  T  L  S  N  G  D  A  G  D  G  T  V  T  D  L  G  D  210

TCAAAGATTAGCTGGAACGGGAACCTGGATTTAGGAGCTTCTGAACCGGAGAATAGCCGAGTCATTGCTTTGTACTCGAATGATTTTGAG 720
 S  K  I  S  W  N  G  N  L  D  L  G  A  S  E  P  E  N  S  R  V  I  A  L  Y  S  N  D  F  E  240

AACGGGTGGAGGATCGGCAATAAGCAGTCGTACGAGGACTACAAGACGTTCATCGGTGGCGGGGATGCCTACGATCTCCTGATTGACTGG 810
 N  G  W  R  I  G  N  K  Q  S  Y  E  D  Y  K  T  F  I  G  G  G  D  A  Y  D  L  L  I  D  W  270

CAGGACGGGACTTACACCGCCTCTGAAGTAGAGGATGAGCTGGTGAATACTGATGCGAACCAGGCTGTTGAGGAGGCCAGTTCTAGCACG 900
 Q  D  G  T  Y  T  A  S  E  V  E  D  E  L  V  N  T  D  A  N  Q  A  V  E  E  A  S  S  S  T  300

ACAGACTTGGTGAACGCGAAGGTTAAGGACTCGTCTCTTGACACCGGTTCGTTCGTGTATGATACCCCTGAGCTGCTTTCCTACCCGTCC 990
 T  D  L  V  N  A  K  V  K  D  S  S  L  D  T  G  S  F  V  Y  D  T  P  E  L  L  S  Y  P  S  330

TTCACAGTCTACGTCGATGCCGGAGAAAACGGATACATCGAGGTAACGAAGCCGACTGGTGACCCCGATATCATCTCCACTTCATCTACA 1080
 F  T  V  Y  V  D  A  G  E  N  G  Y  I  E  V  T  K  P  T  G  D  P  D  I  I  S  T  S  S  T  360

GAAATTAAGGAGGGAGATGAGGGAACTGTAACCGCCACCGTTGAGAATGTCGGAGATGGTGAGGGCGAGTTCTCTGGAAGGCTTTCCAGC 1170
 E  I  K  E  G  D  E  G  T  V  T  A  T  V  E  N  V  G  D  G  E  G  E  F  S  G  R  L  S  S  390

TGCGGTGAAGGCTTCAGTATTGTCGACGACCAGAATACGAAGAATGTGGGTGCCGGAGAATCGGTGACATACAGTTTTGATGTCGCTTTC 1260
 C  G  E  G  F  S  I  V  D  D  Q  N  T  K  N  V  G  A  G  E  S  V  T  Y  S  F  D  V  A  F  420

AGTTCTGTCAGCAGTGAGTCGAAGGAGATCTCGGGCAGCTGCACGTTCGAGGTGAACGGCGTCGAGTCCAGTGATTCCACCAGTGTTTCT 1350
 S  S  V  S  S  E  S  K  E  I  S  G  S  C  T  F  E  V  N  G  V  E  S  S  D  S  T  S  V  S  450

GTGACCGGTATCCAGCAGAGTGAGTGTAATCCAGGCGATCAGAGACGGGAAAAGAATGAGAATGACCGCTGGGAGATCTACACCTGCCAG 1440
 V  T  G  I  Q  Q  S  E  C  N  P  G  D  Q  R  R  E  K  N  E  N  D  R  W  E  I  Y  T  C  Q  480

GACAACGGCCTGACCTACGAATATGATGTAACTTGTGCAGAGGACGAGAAGGCTGTAGCTCAGGGAGATAACCAGTTTTCCTGTGAGAAA 1530
 D  N  G  L  T  Y  E  Y  D  V  T  C  A  E  D  E  K  A  V  A  Q  G  D  N  Q  F  S  C  E  K  510

CAGGACGATGATTCAGGTGGCGGAGACAACACTGGATCTGACTCCGGACTGTTCTCGAATCTGTTCGGAGGTTCTGGTTCCGGAGATCTG 1620
 Q  D  D  D  S  G  G  G  D  N  T  G  S  D  S  G  L  F  S  N  L  F  G  G  S  G  S  G  D  L  540

CTTACACAGGTTCACACAGCTCTGTCTATCCTCGCGGGGCTTGTTGCAGGCTTCTTCGGTTATAGAGGAGCcCGATGGATTCACGGCGAG 1710
 L  T  Q  V  H  T  A  L  S  I  L  A  G  L  V  A  G  F  F  G  Y  R  G  A  R  W  I  H  G  E  570

ACTGATATAAAAGGAGGCTTCAAGCTGGAGTCCCGGAACGTAAGCCGTGTTAAGAGAGGTAGTCCTGTAGCCGGTATTGTCGGAGCTGTA 1800
 T  D  I  K  G  G  F  K  L  E  S  R  N  V  S  R  V  K  R  G  S  P  V  A  G  I  V  G  A  V  600

CTCGGGTTTGTCGTAGGATACGGTGTTGCCTCAGTCTTCCATCCAGTAGTTCAGATTATAGTGGTTCTGGGAATTGCTGTTGGCTTGTAC 1890
 L  G  F  V  V  G  Y  G  V  A  S  V  F  H  P  V  V  Q  I  I  V  V  L  G  I  A  V  G  L  Y  630

TATTTCAGGCTAGAGGGCCCGCGGTTCGAAGGTAAGCCTATCCCTAACCCTCTCCTCGGTCTCGATTCTACGCGTACCGGTCATCATCAC 1980
 Y  F  R  L  E  G  P  R  F  E  G  K  P  I  P  N  P  L  L  G  L  D  S  T  R  T  G  H  H  H  660

CATCACCATTGAgagctcgcatgcactagt 2010
 H  H  H  *                    670
```
